# Capturing the start point of the virus-cell interaction with high-speed 3D single-particle tracking

**DOI:** 10.1101/2021.12.17.473224

**Authors:** Courtney Johnson, Jack Exell, Yuxin Lin, Jonathan Aguilar, Kevin D. Welsher

## Abstract

The early stages of the virus-cell interaction have long evaded observation by existing microscopy methods due to the rapid diffusion of virions in the extracellular space and the large 3D cellular structures involved. Here we present an active-feedback single-particle tracking method with simultaneous volumetric imaging of the live cell environment to address this knowledge gap to present unprecedented detail to the extracellular phase of the infectious cycle. We report previously unobserved phenomena in the early stages of the virus-cell interaction, including skimming contact events at the millisecond timescale, orders of magnitude change in diffusion coefficient upon binding, and cylindrical and linear diffusion modes along cellular protrusions. Finally, we demonstrate how this new method can move single-particle tracking from simple monolayer culture towards more tissue-like conditions by tracking single virions in tightly packed epithelial cells. This multi-resolution method presents new opportunities for capturing fast, 3D processes in biological systems.

**One-Sentence Summary:** Active-feedback 3D single-particle tracking enables an unprecedented look at the early stages of virus-cell interactions.

## Main Text

The ongoing SARS-CoV-2 pandemic has demonstrated with frightening clarity the need for fundamental research in physical virology to exploit and counter the mechanisms of viral infection. The first point of contact with the host organism occurs in the extracellular space of the epithelia, whose cells form a tightly packed arrangement protected by an extended mucus layer and glycocalyx (*1-5*). The structure of the extracellular matrix (ECM) has been shown to be critical to the viral lifecycle, undergoing changes in structure and composition upon introduction of viral pathogens (*6*) and hosting biofilm-like viral assemblies for cell-to-cell transmission (*7*).

Single-particle tracking (SPT) methods have emerged as a powerful tool in our understanding of viral infection (*8-12*). These methods have uncovered virion binding mechanisms to the cell surface (*13, 14*), distinguished internalization pathways (*15-19*), identified the cellular location of envelope fusion (*20-26*), characterized cytoskeletal trafficking (*14, 19-22, 27-33*), and demonstrated how viruses hijack filopodia to efficiently infect neighboring cells (*19, 23, 34-39*).

Despite these advances, it has thus far not been possible to follow virions starting in the extracellular space with sufficient detail to interrogate this important phase of viral infection. This is because extracellular diffusion occurs across depth ranges exceeding 10 μm and at diffusive speeds 2-3 orders of magnitude greater than the highly confined processes followed by conventional single particle tracking methods. Even the most advanced image-based SPT methods such as spinning disk confocal and light sheet microscopy (*40, 41*) are unable to meet these challenges as they suffer from limitations caused by attempting to simultaneously track and image disparately scaled objects on a single platform.

One way to understand these limitations is to consider the sampling of trajectory points in terms of the number of localizations per second (loc/s), which for image-based tracking methods is given by the volumetric imaging rate (Fig. 1A). As volumetric data is acquired frame-by-frame, the temporal resolution scales poorly as the axial extent of the process in question grows. Yet shrinking the volume Z size reduces the likelihood that the virion will remain in the volumetric field of view, shortening the observation duration and thus, absolute number of localizations acquired. Additionally, as camera exposure times decrease, photon limitations when trying to image single particles become a limiting factor. Given a modest value of 16 z-slices spaced over an axial range sufficient to capture extracellular diffusion, even the fastest light sheet systems can theoretically acquire as many as 6.25 loc/s at best. Overcoming these fundamental limitations in speed and axial range motivated the development of active-feedback tracking methods which focus exclusively on the tracked particle (*42*). Such methods can localize particles with high spatial and temporal resolution, but the cost to obtaining this enhanced temporal resolution is loss of environmental context. While such methods have been useful in studying particle dynamics in isolation, this missing context has precluded the application of such methods to studying particle-environment interactions.

**Fig. 1.**
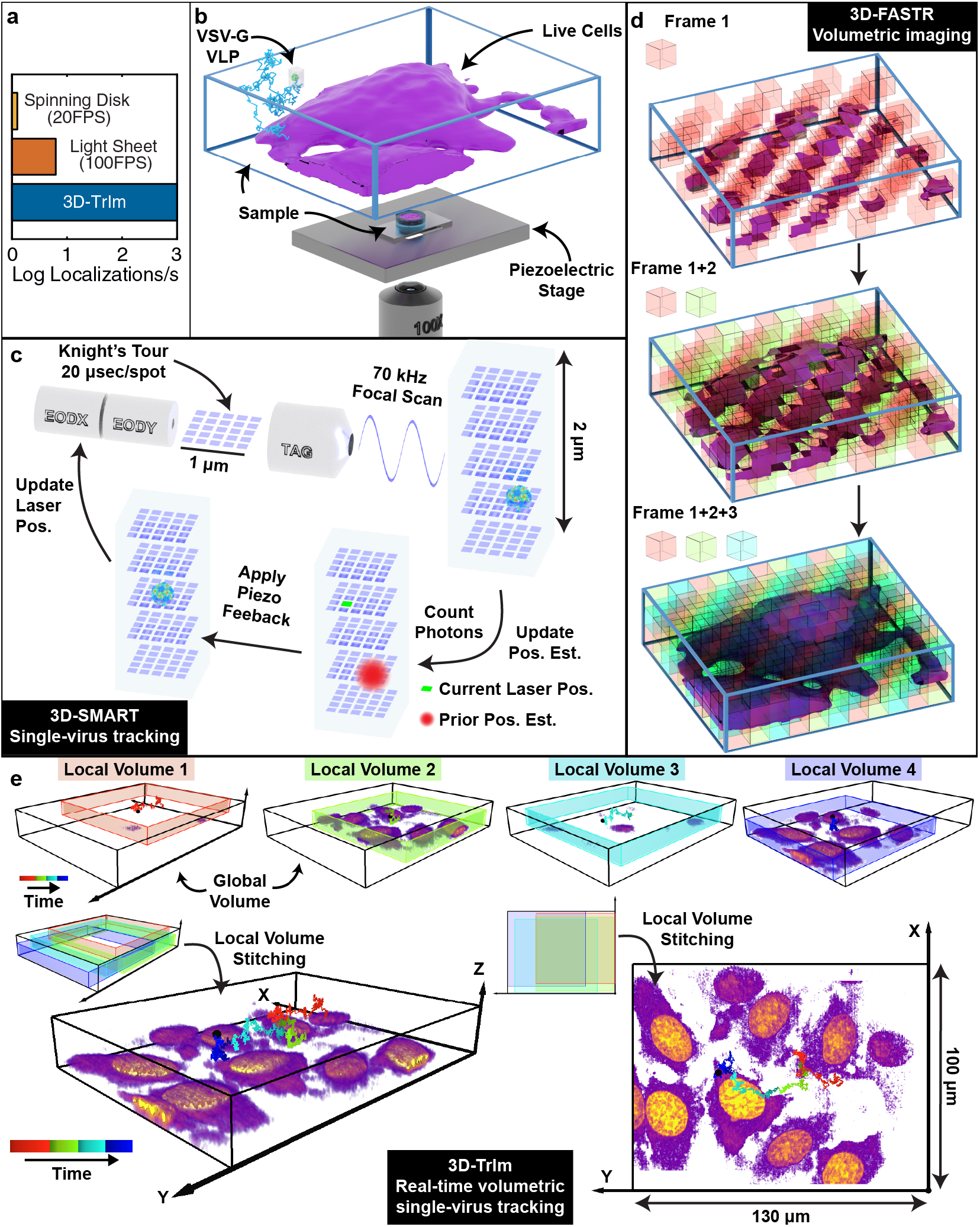
3D Tracking and Imaging (3D-TrIm). (**A**) Sampling rate comparison among spinning disk, light-sheet, and 3D-TrIm. (**B**) Experimental setup. Fluorescently labeled VLPs are added to live cells plated on a coverslip. The sample is placed on a heated sample holder mounted on a piezoelectric stage. (**C**) Overview of 3D-SMART tracking of single viruses. The electro-optic deflector (EOD) and tunable acoustic gradient (TAG) lens rapidly scan the local particle area. Photon arrival times and the current laser position are used to calculate the position of the virus within the scan area. Using the measured position, the piezoelectric stage moves to recenter the virus within the scan area. (**D**) Concept of 3D-FASTR volumetric imaging. By outfitting a traditional 2-photon LSM with an ETL, a repeatable, tessellated 3D sampling pattern can be generated during each frame-time. Over a set number of frame-times, the entire volume is sampled. (**E**) Construction of global volumes in 3D-TrIm. As the virus diffuses, 3D-SMART moves the sample, and the 3D-FASTR imaging system collects sequential volumes from different areas around the particle. These time-resolved local volumes can be used to generate an integrated global volume (see also: movie S1).

To address this gap, we present 3D Tracking and Imaging Microscopy (3D-TrIm), a multi-modal instrument that integrates a real-time active-feedback tracking microscope (*43-45*) with a volumetric imaging system (*46*) capable of simultaneously tracking the high-speed dynamics of extracellular virions while imaging the surrounding 3D, live-cell environment (Fig. 1A). This multi-modal approach provides unique value which cannot be extracted by its independent components alone, and as demonstrated here provides unprecedented glimpses into the initial moments of the virus-cell interaction. Using 3D-TrIm, we acquired over 3000 viral trajectories and demonstrate the benefits of how this contextualized tracking uses localizations in time analogously to super-resolution methods. This “super-temporal-resolution” enables not just the formation of trajectories, but the detection of changes in diffusive regimes and the tracing out of nanoscale structural features. Finally, 3D-TrIm was applied to advance SPT from monolayer cell culture to tightly packed, 3D epithelial culture models, providing a new window into this critical stage of viral infection.

## Results

Capturing extracellular viral dynamics requires the ability to track a rapidly diffusing target in three dimensions. To accomplish this, we apply the recently developed 3D single-molecule active real-time tracking (3D-SMART) (*43, 44*). 3D-SMART is an active-feedback microscopy method that uses real-time position information to “lock on” to moving fluorescent targets. Critically, 3D-SMART can capture particles diffusing at up to 10 μm^2^/s with only a single fluorophore label (*43*), making it an ideal choice for capturing diffusing virions (*45*).

The implementation of 3D-SMART in the current work is similar to previous reports. Briefly, a rapid 3D laser scan excites photons from the fluorescently labeled diffusing viral particle (Fig. 1B, movie S1). Photons are collected on a single-photon counting avalanche photodiode (SPCM-APD), which also acts as a pinhole to provide confocal detection and prevent detection of out-of-focus virions. The photon arrival times are used to calculate the real-time position every 20 µs which is then used in an integral feedback loop to move the sample via a piezoelectric stage, effectively fixing the moving target in the focus of the microscope objective. In the event that a second virion diffuses into the tracking volume, a spike in intensity is recorded. Trajectories are split at these “jump” points so that each trajectory represents a single virion. 3D-SMART produces a three-dimensional particle trajectory with 1000 loc/s, limited here by the 1 ms mechanical response time of the stage, with localization precision up to ∼ 20 nm in XY and ∼ 80 nm in Z (fig. S1) and duration only limited by the travel range of the piezoelectric stage (75 µm × 75 µm × 50 µm; XYZ).

While 3D-SMART is ideally suited for capturing the fast dynamics of extracellular viral particles over vast axial ranges, alone it lacks environmental context. A complementary rapid volumetric imaging method is needed to capture the live-cell environment. The choice of imaging system is non-trivial as the active-feedback nature of 3D-SMART requires a moving stage which places two constraints that must be considered. First, imaging with camera-based methods, such as light-sheet microscopy, would result in motion blur across a single exposure time; the chosen imaging modality must be faster than the 1 ms response time of the piezoelectric stage. Second, the stage cannot be stepped to perform “z-stacking” for volumetric imaging.

To meet these criteria, we implement 3D Fast Acquisition by z-Translating Raster (3D-FASTR) microscopy to provide rapid volumetric imaging around the tracked viral particle (*46*). 3D-FASTR uses a two-photon laser scanning microscope outfitted with an electrically tunable lens (ETL). The short pixel dwell time of the raster scan (∼ 1 µsec) prevents motion blur, and the ETL performs remote focusing of the imaging laser for 3D imaging. In contrast to a conventional z-stack, 3D-FASTR performs a continuous focal scan across an 8 µm range. At optimized scan frequencies, the combination of the XY raster scan and the Z focal scan evenly samples voxels in a tessellating pattern which tiles to scan the entire volume (Fig. 1C, movie S1). At short acquisition times, unsampled voxels will have many sampled neighbors such that the volumetric imaging rate can be increased 2-to-4-fold over conventional image stacking methods through interpolation (*46*). The ability to rapidly image volumes without moving the sample makes 3D-FASTR the perfect complement to 3D-SMART.

To integrate the two microscopes into a combined platform, excitation and emission for both tracking and imaging are routed through a single shared objective lens (100×, NA = 1.4) on an inverted microscope stand outfitted with a 3D piezoelectric stage. The piezoelectric stage is mounted on a secondary motorized stage that remains static during acquisition but can be moved between trajectories to capture different areas of the sample. (fig. S2). Registration of the 3D-SMART and 3D-FASTR systems is required to generate combined 3D-TrIm datasets (fig. S3-4). The piezoelectric stage represents a shared spatial grid between the two microscopes; the real-time stage position is a measure of the tracked particle’s location that is tagged to the imaging XYZ voxel position. The motion of the piezoelectric stage translates the 3D imaging field of view, expanding the total imaging volume size based on how far the virion diffuses. This effect is most dramatic in Z where the 8 µm 3D-FASTR imaging range moves across the 50 µm Z stage range; These tagged positions are then used with calibration information (fig. S5-6) to place each image voxel in the shared global volume space. This per-voxel registration allows for an arbitrary volume acquisition rate; voxels can be accumulated over the duration of an entire trajectory to form a single global volume or spread across multiple local volumes over time (shown here at ∼ 17 sec/volume) to construct sequences over the course of the trajectory (Fig. 1D).

For this pioneering study into the early stages of the virus-cell interaction, we investigated the journey of vesicular stomatitis virus (VSV)-G-pseudotyped lentiviruses from the extracellular solution to the cell surface and beyond. We chose this specific virus-like particle (VLP) due to its wide cellular tropism. A behavior attributed to the ubiquitous host entry partner of VSV-G, the low-density lipoprotein receptor (LDLR) (*47*), which has established VSV-G as the standard for gene delivery (*48*). We utilized an efficient and well-characterized internal virus labeling approach whereby fluorescent proteins fused to the HIV-1 viral protein R (Vpr) are packaged into budding lentiviruses (*49*). Successful incorporation of eGFP.Vpr into single virions was confirmed by tracking VSV-G eGFP.Vpr in the absence of cells and by immunofluorescence experiments (fig. S7-9). Based on a photobleaching analysis that indicates a single eGFP-Vpr has an intensity of ∼ 2 kHz, each VLP tracked in this study has somewhere between 10 and 100 eGFP-Vpr packaged within its capsid (fig. S10). Critically, these VLPs were shown to remain infective (fig. S11).

To initiate data collection, the 3D-TrIm microscope locates diffusive virions by searching a plane approximately 5 μm above the apical cell surface so that viruses are captured prior to virus-cell first contacts. When a VLP enters the 3D-SMART tracking volume, a significant increase in detected photons above the background level is observed on the single-photon counter, which triggers the active-feedback tracking loop and 3D-FASTR image acquisition.

### 3D-TrIm provides super-temporal-resolution of contextualized particle trajectories

To demonstrate the impact of this new method, we acquired extracellular trajectories of our virion using both 3D-TrIm and a state-of-the-art Andor Dragonfly spinning-disk-confocal (SDC) microscope. The side-by-side data shown in Fig. 2 below demonstrate the vast difference in the amount of data available from 3D-TrIm compared to SDC data. The SDC microscope shows the trajectories of multiple extracellular virions, the majority of which remain in the trackable field of view long enough to obtain only a few sample points. Such sample points are characterized as long, straight lines averaging the behavior between frame exposures. Tracking and imaging are coupled together so the field of view is confined to a fixed area and only 1 localization can be made per volume acquired. The overlaid volumes show that over 3 acquired volumes there are 3 localizations. In contrast, the 3D-TrIm trajectory of a single virion features thousands of sample points, capturing the contour of motion instead of straight lines. This localization rate and field of view are completely decoupled from the volume rate. Instead, over the course of 3 volumes there are thousands of localizations per volume and the imaging field of view moves with the particle, reducing bleaching. Notably, the utility of our super-resolved tracking is not just in the ability to form trajectories, but to extract quantitative and qualitative meaning from the data. This super-temporally-resolved data can extract not just a single approximated diffusion coefficient, but detect changes in diffusion speed and mode due to interactions.

**Fig. 2.**
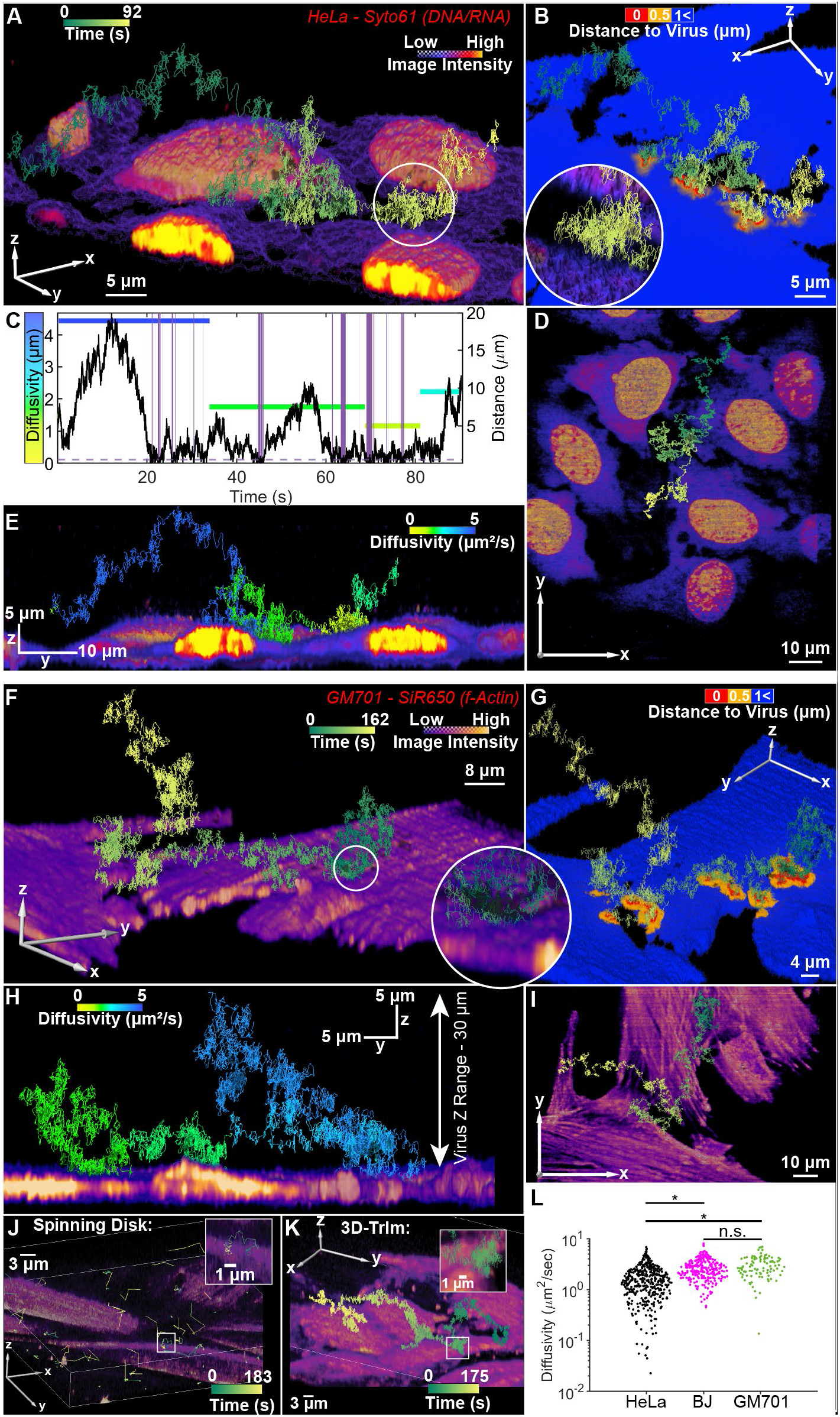
Minute-long high-resolution 3D tracking and volumetric live-cell imaging reveals transient VSV-G VLP contacts with the cell surface. (**A** and **B**) 3D reconstruction of a single VSV-G VLP trajectory with live HeLa cells stained with SYTO61 from a 4D data set. Cells are color-coded by intensity and distance from the cell surface to the virus trajectory in (A and B), respectively. **Circular inset**: enlarged view of “skimming” event. (**C**) Correlation between diffusivity and cell-to-virus distance. Skimming events are highlighted in purple. Skimming event shown in circular inset of (B) occurs between 68-78 sec. (**D**) Top-down (xy) complete imaging area with trajectory. (**E**) Lateral (yz) view with trajectory color-coded by diffusion coefficient segments calculated by change-point analysis. (**F**-**J**) Identical to (A-E) except VSV-G trajectory co-registered with live GM701 fibroblast cells stained with SiR-Actin (movie S2). Skimming event shown in circular inset of (F) occurs between 37.5-47.5 sec. (**J**) Extracellular viral diffusion collected on an Andor DragonFly spinning disk confocal microscope. The same area was sampled continuously. Each volume has a depth of 8 μm split into 16 z planes. Virus particles (eGFP-Vpr VSVG) and live cells (HeLa, SiR650-Actin stain) were imaged simultaneously with a camera exposure time of 40 msec. (**K**) Extracellular viral diffusion collected on 3D-TrIm microscope. A single virus particle (eGFP-Vpr VSVG) was tracked continuously (1 msec sampling shown) and the surrounding area was imaged 5 μm below the particle and 3 μm above, to give an approximate 8 μm volume of cellular environment (HeLa, SiR650-Actin stain). The resulting trajectories over the entire acquisition period are shown and color-coded by time. (**L**) Diffusivity of VSV-G VLP trajectory segments identified by change-point analysis. (HeLa, BJ, and GM701 n = 413, 255, and 100 segments, from 66, 105, and 47 trajectories, respectively). Statistical significance was assessed by 2-sample *t*-test, on the p-value at p < 0.01.

### 3D-TrIm reveals that the extracellular dynamics of virions are cell-type and distance-dependent

An example trajectory of these early interactions is shown in Fig. 2A-C, which features live HeLa cells fluorescently labeled with the nucleic acid stain SYTO61 (additional examples: fig. S15-19). Taking advantage of the co-registered individual local volumes (fig. S12), a global 3D render of the cells was constructed using the maximum intensity of each voxel over time. The viral trajectory is constructed from the piezoelectric stage coordinates, and visualized here at the 3 ms sampling intervals used for diffusivity calculations. The single VSV-G VLP was initially captured in the search plane and diffused for ∼ 90 seconds, reaching a maximum height of nearly 20 μm above the cell surface. The virion closely approaches and intermittently touches the surface of several cells over the course of the trajectory.

These periods of transient contact (“skimming”) were quantified in two ways. First, changepoint analysis was applied to identify changes in the diffusive state of the VLP as it traverses the volume (*50, 51*). The extracted diffusion coefficients are shown as color-coded lines in Fig. 2D. Second, the distance between the VLP and the cell surface was calculated using segmented volume data (fig. S13). This VLP-cell distance is visualized in Fig. 2B, a color-mapped volume with yellow/red indicating approach within 1 μm/0.5 μm (comparable to the axial extent of the 3D-FASTR point-spread function, fig. S14), respectively. This value of 0.5 μm was used as the threshold value for determining contact. The trajectory reveals multiple and repeated contact events at various surface locations (highlighted in purple in Fig. 2D). While the HeLa cell shown is LDLR positive, these skimming events are not restricted to receptor-positive cells but are also observed in GM701 fibroblasts (low LDLR expression level) (Fig. 2F-I and movie S2).

These transient contacts can be quantified to yield their occurrence (fig. S21A), frequency (fig. S21B), and dwell time (fig. S22) for each cell type. These contacts can be as short as several milliseconds in duration but numerous per trajectory, without promoting long-duration viral attachment. Exponential fitting of contact event durations on HeLa cells yielded two populations with dwell times of 23 ± 1 and 102 ± 5 msec (fig. S22A). The shorter dwell time is consistent with free diffusion, while the longer dwell time indicates an interaction of the VLP with the cell surface or extracellular matrix. These dwell times were longer for HeLa (102 ± 5 msec) compared to the low-LDLR GM701 fibroblast cells (88 ± 8 msec, fig. S22B), though they exhibited identical fast components. Since these two diffusive states are not correlated with the presence (BJ) or absence (GM701) of LDLR, it can be concluded that these initial interactions are not related to receptor priming but potentially related to cell morphology (fig. S22C).

The increased dwell time near the cell surface also correlated with slower diffusion in the presence of HeLa cells. A trajectory-wide analysis of “skimming” VSV-G VLPs showed 80% of trajectories acquired in the presence of HeLa cells have diffusion coefficients within the range of 0.5 to 3.4 µm^2^/s (fig. S23), consistent with diffusive rates in the presence of cells in previous reports (*13, 14, 51*). Interestingly, VLPs exhibited increased diffusivity of 1.3 to 4.5 µm^2^/s for thinner fibroblast cells, revealing a cell-type dependence on extracellular diffusivity (Fig. 2K, 2-sample *t*-test, p < 0.01). The unique correlative imaging and high-speed tracking capabilities of the 3D-TrIm microscope enabled a quantitative examination of the relationship between VLP diffusion and distance from the cell surface. This correlation analysis revealed a distance-dependent relationship, with VLPs slowing down as they near the cell surface. Statistically-significant inhibition of diffusion was observed within 2.5 µm of the cell surface for all cell types tested (fig. S23). Again, compared to the fibroblasts, VLPs in the presence of HeLa trended towards lower diffusion coefficients near the cell surface (fig. S23). This difference again suggests morphological effects imposed by the more irregular shaped HeLa compared to flatter, more uniform fibroblasts.

While all three cell types showed a trend towards slower diffusion near the cell surface, each still displayed a significant fraction of VLPs with segments of diffusion > 1 µm^2^/s at less than 1 µm from the cell surface. Again, this effect was cell-type dependent, with BJ cells showing a much higher percentage of fast diffusing VLPs near the surface (BJ: 33.6% ± 11.8%, HeLa: 16.8% ± 3.9%, GM701; 16.7% ± 7.3%). This fast diffusion near the cell surface suggests a complex interplay between the extracellular environment and VLP dynamics. Such short-lived contact events are resolvable with 3D-TrIm due to the unique combination of super-temporally-resolved tracking combined with 3D Imaging that can discern cellular contact on the millisecond timescale.

### 3D-TrIm captures the transitional dynamics of binding

In addition to the transient contacts observed above, long-term binding events of individual VSV-G VLPs on the cell surface were also captured by 3D-TrIm. Unlike skimming, these binding events display dramatic diffusivity changes upon contact. In the first example (Fig. 3A-D, fig. S24, and movie S3), the virus initially undergoes several skimming events, marked by close approach to the surface of multiple cells with subtle changes in the diffusion coefficient. At ∼ 70 s, as the virus-cell distance reaches a minimum, viral diffusivity drops by two orders of magnitude and remains bound for several minutes (indicated by the pink box in Fig. 3D). In such cases where motion of the particle is confined, the observation duration is limited by photobleaching of the virion to ∼5 minutes at the reported laser power. Photobleaching of the cells occurs over a longer duration due to sampling a larger area less frequently.

**Fig. 3.**
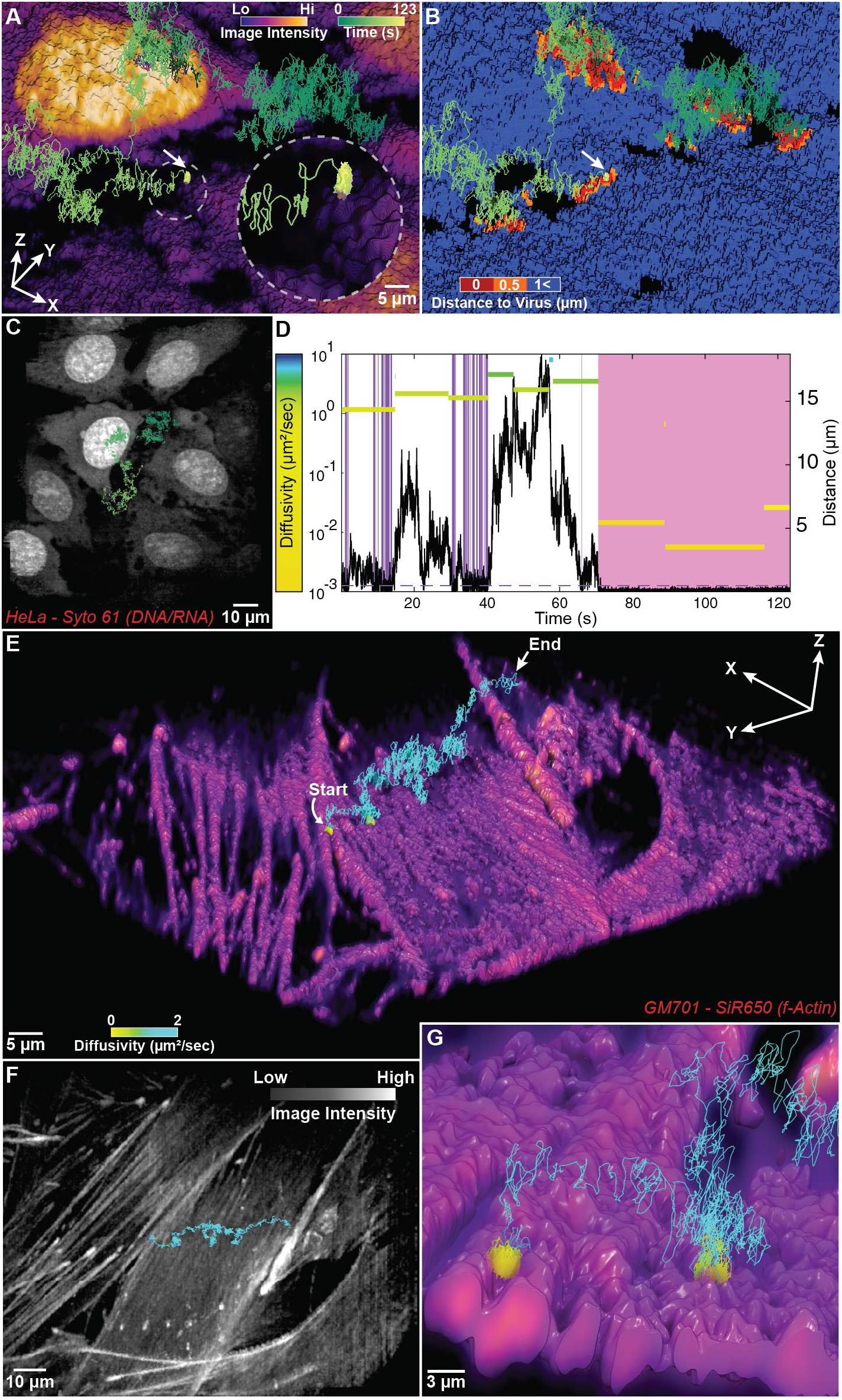
VSV-G Binding. **(A** and **B)** 3D TrIm render of live nucleic acid-labeled HeLa cells. (**A**) Intensity volume render (see also: movie S3). Grey circular inset shows moment of binding. (**B**) Virus-Cell distance render. (**C**) Maximum intensity projection (MIP) with trajectory overlay. (**D**) Trajectory diffusivity and virus-cell distance time trace. Skimming events are highlighted in purple, binding events are highlighted in pink. (**E-G**) 3D TrIm render of live actin-labeled GM701 cells with trajectory color-coded by diffusivity (see also: movie S4). (**F**) MIP with trajectory overlay. (**G**) Close-in view of VLP binding sites.

It was also observed that bound viral particles detach after landing and diffuse away or even bind again elsewhere. Fig. 3E-G shows a VLP initially bound between two protrusions on an actin-stained GM701 fibroblast (see also fig. S25, movie S4). This bound state (D ∼ 0.04 µm^2^/s, Fig. 3E, yellow) persists for ∼ 23 sec, before detaching to free diffusion (D > 2 μm^2^/s). After only ∼ 4 sec of free diffusion, viral diffusivity drops to less than 0.1 µm^2^/s upon contact and remains bound for several seconds before detaching again to free diffusion and ultimately leaving the trackable volume (additional example: fig. S26). Combined with the msec-scale dwell times above, these multiple and long-term binding events suggest that cell-VLP attachment events cover a large range of timescales.

### 3D-TrIm traces out nanoscale features on cellular protrusions

In addition to the ability of 3D-TrIm to resolve milliseconds-long contact and sample changes in diffusive regime, 3D-TrIm’s highly sampled trajectories can trace out nanoscale structures on the cell surface. Similar to how super-localization of emitters enables nanoscale resolution of cellular structures in super-resolution methods, the high 3D precision and rapid sampling of 3D-SMART turns the virus into a nanoscale pen that draws out local features smaller than the diffraction limit that can be contextualized through our simultaneous larger-scale imaging.

One example is the interaction between VLPs and cylindrical protrusions on the cell membrane. Using 3D-TrIm, we observed viral particles binding directly to these structures from the extracellular space (Fig. 4A-E, fig. S27 and movie S5). While not stained and therefore not visible in the image, the virus’s path along the surface creates a high-resolution map, carving out the nanoscale cylindrical morphology of the protrusion surface (Fig. 4D, E). The feature traced by this VLP protrudes ∼ 1 μm vertically from the cell surface (Fig. 4D). A best-fit cylinder gives a radius of 105 ± 8.4 nm (Fig. 4E), consistent with the observed size of filopodial protrusions, and notably, demonstrating the ability for 3D-TrIm to super-resolve tracked features beyond the diffraction limit. These structures were dynamic and resulted in changing cylindrical structure within the viral trajectory over time (Fig. 4F-I), with radii tapering for more distal parts of the structure. VLPs on these structures were consistently measured to have diffusivity values ∼ 0.01 μm^2^/s (fig. S27-S29), nearly identical to other types of membrane diffusion observed (fig. S32). Viral trajectories collected with 3D-TrIm were also able to super-resolve hemispherical membrane blebs (Fig. 4J, K, Fig. S33), highlighting again how high sampling rate 3D tracking can result in super-resolution structural tracing.

**Fig. 4.**
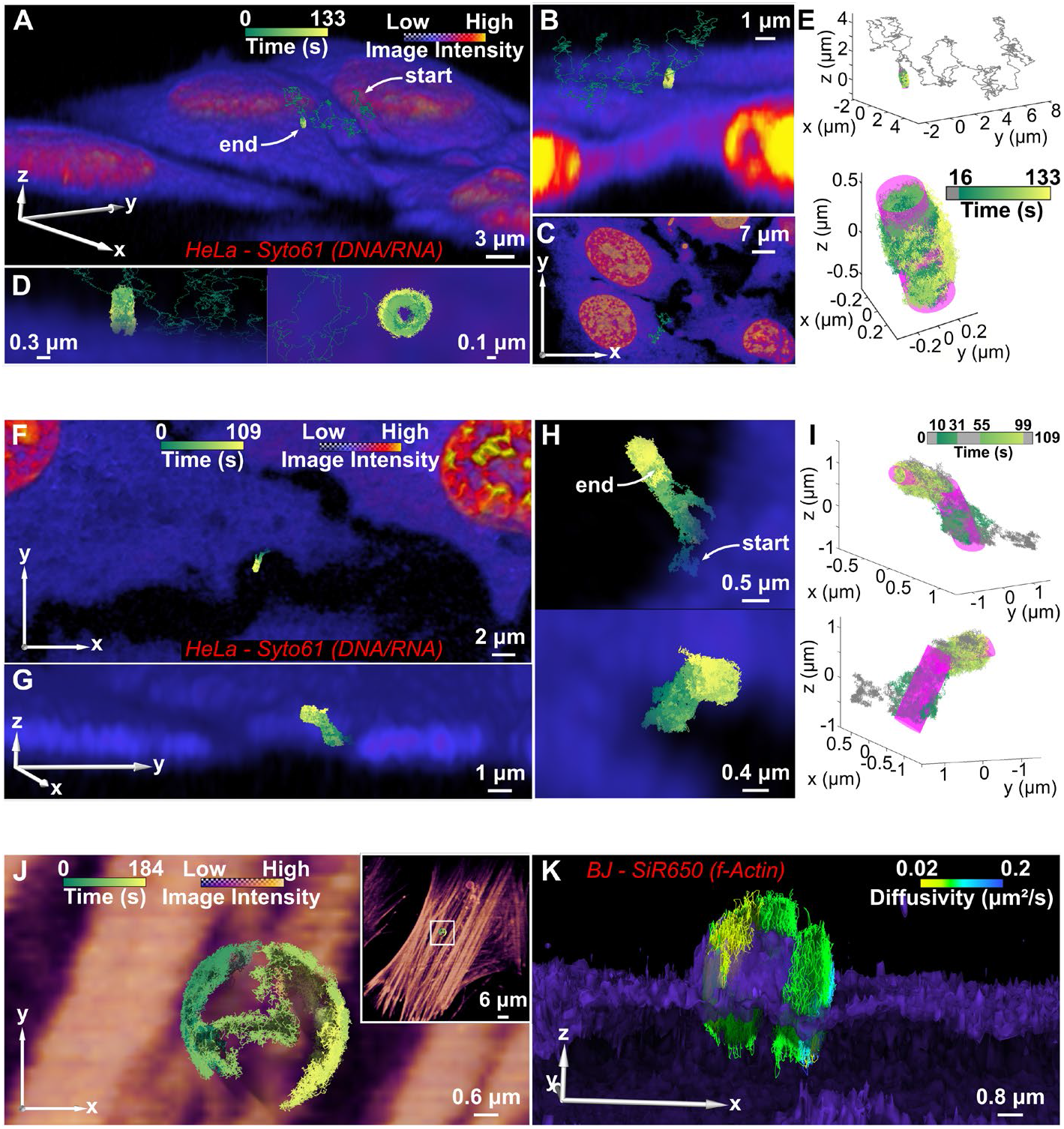
Virus interaction with cellular protrusions. (**A**) VSV-G landing on protrusion from the surface of a live SYTO61-stained HeLa cell (see also: movie S5). (**B**) Exploded view with slices through x and y axes. (**C**) Top-down (xy) complete imaging area with trajectory. (**D**) Magnified views showing the hollow cylinder the VLP draws out while diffusing around the protrusion. (**E**) Cylindrical fitting of the bound portion of the trajectory in (A-D) (16-133 sec), radius = 105 ± 8.4 nm. (**F**) Additional high-resolution trajectory of VLP on a protrusion captured after landing. (**G**) Lateral view, VLP tracking captures change in protrusion shape. (**H**) Enlarged views showing VLP progression after starting from a proximal position on the membrane. (**I**) Cylindrical fitting of base (10-31 sec) and extremity (55-99 sec) of the trajectory in (F-H), radius = 153 ± 8.7 nm and 77 ± 8.4 nm, respectively. Average protrusion radius of 125 ± 16 nm (n = 5). (**J**) Top-down view of VSV-G traveling diffusing on membrane bleb of actin-stained BJ fibroblast cells. **Inset**: enlarged view. (**K**) 3D volume rendering for data shown in (J), except trajectory colored by diffusion coefficient.

### Single viral dynamics in the epithelia: beyond monolayer cell culture models

As is typical of SPT experiments, the previous examples were captured in the presence of monolayer cultures, which have been shown to differ from more realistic tissue models (*52*). Here, we demonstrate the potential for 3D-TrIm to operate in complex environments at considerable depths in systems that more closely approximate the infection routes of viruses *in vivo*. A system of particular relevance to the extracellular dynamics of viruses (and to respiratory viruses in particular) is the epithelia, which is protected by a thick mucus layer (*1, 2*) and a size-excluding pericilliary layer (PCL) (*53*). For well-differentiated epithelial cells to form tightly packed arrangements *in vitro*, they must be grown on a semipermeable membrane support to allow access to the basolateral layer and provide a more realistic growth environment. The thick (>10 µm), tightly-packed epithelial layer cannot be grown directly on a coverslip, making observation of dynamics in these critical systems impossible in conventional microscopy methods. In contrast, 3D-TrIm’s large axial range enables unprecedented high-speed single-particle tracking in these more biologically relevant tissue models.

Epithelial model systems were prepared by growing HT29-MTX cells on a semipermeable membrane support and inverted so that the cells were suspended above the coverslip surface (fig. S34). The diffusion of viruses into this tightly packed layer can be captured from this vantage with msec temporal resolution (Fig. 5A-D, fig. S35-36 and movie S6). The VLP experiences confinement near the surface of multiple cells for nearly 3 minutes. However, it is important to note the difference between sub-diffusion/confinement and diffusivity. The average diffusion coefficient for VSV-G traveling in this complex environment is 1.29 ± 0.44 µm^2^/s, in the range of freely diffusing virus particles and at least 2 orders of magnitude higher than bound or internalized processes (fig. S9), suggesting the extracellular matrix does not present a significant physical barrier. Critically, despite this confinement, the virion manages to diffuse more than 10 µm across the Z axis during the trajectory.

**Fig. 5.**
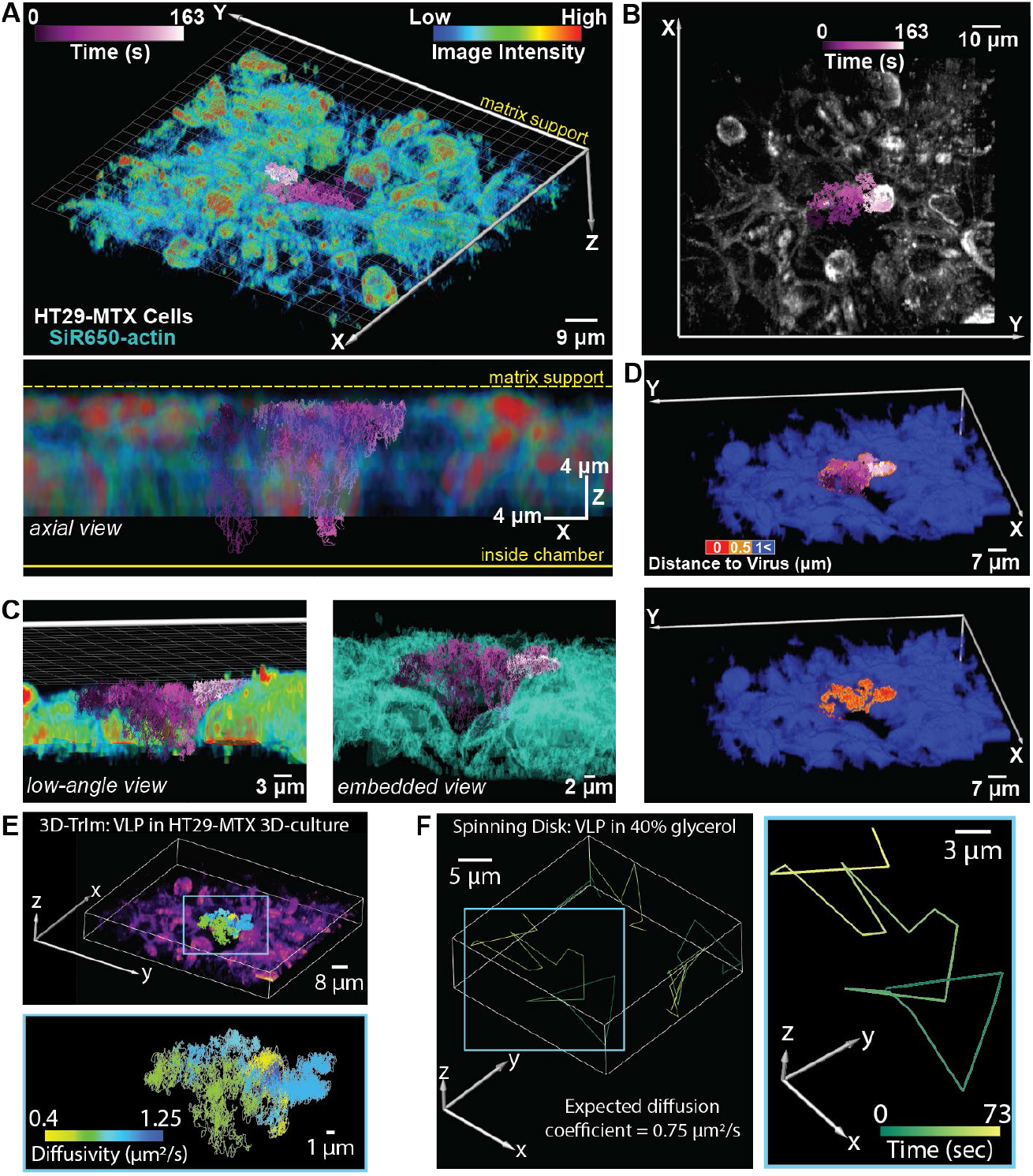
VSV-G VLP diffusing through multi-layered epithelial cells. (**A**) **Top:** 3D reconstruction from a 4D data set covering 10 local volumes, at 10 FPV (see also: movie S6) of suspended HT29-MTX cells grown on inverted matrix support and labelled with SYTO61. Co-registered VSV-G VLP trajectory color-coded by time. **Bottom:** XZ view. (**B**) Top-down (xy) MIP. (**C**) **Right**: magnified cut-through view showing VSV-G VLP confined within vacancy as it diffuses around the edge of a cell. **Left**: Isosurface render of cells highlighting sample density. (**D**) **Top:** the distance of the trajectory to the cell surface is projected as a distance map on the cell volume, with the trajectory color-coded by time overlayed. **Bottom:** Volume with trajectory omitted, showing areas close contact on the cell surface. (**E**)**Top:** The resulting trajectory is color-coded by diffusion coefficient, using MSD and change-point analysis. **Bottom:** Trajectory with volume omitted, showing highly sampled trajectory across a large range in diffusivity. (**F**) The same fluorescently-labeled virus-like particles (eGFP.Vpr VSVG) were tracked in 40 % glycerol (v/v) on an Andor DragonFly spinning disk confocal microscope, with a camera exposure time of 40 msec. A single volume (∼8 μm split into 16 z planes) was sampled continuously. The resulting trajectories over the entire acquisition period are shown and color-coded by time. (right).

This large accessible depth range coupled with near-free diffusivity precludes study with even advanced image-based tracking. To demonstrate the benefit of 3D-TrIm in studying this diffusive regime, we suspended VLPs in 40% glycerol, which based on our solution-phase tracking should reduce the particle diffusion to ∼0.75 µm^2^/s. Fig. 5F shows that when imaged at this similar diffusivity, the slower volume sampling of the spinning disk makes it impossible to extract details of local diffusion from the undersampled trajectory. In contrast, the highly sampled trajectory from 3D-TrIm reveals significant (and previously invisible) changes in virion dynamics as it hugs the cell surface (Fig. 5E).

This rapid but trapped diffusion is in stark contrast to viruses near monolayer cultured cells, where the rapid diffusion limits the dwell time near the cell surface. Notably, this example demonstrates the capability for 3D-TrIm to track at low signal levels (35 kHz) in complex environments, which is a significant advantage over other prior active feedback tracking methods, which require several hundred kHz for successful tracking (*51, 54*).

## Discussion

There are several features of 3D-TrIm that make it extendable to future studies. First, not restricted to capturing rapid extracellular dynamics, 3D-TrIm’s high-resolution tracking ability extends to the later, internalized stages of the infectious cycle (movie S7 and fig. S37-38). These data demonstrate that the VLPs in this study exhibit the hallmarks of infectious virions in viable cells, undergoing normal cellular processes like endocytosis and intracellular trafficking. 3D-TrIm is fully compatible with live-cell studies and active biological processes.

Second, 3D-TrIm can observe single-particle dynamics within the same area over long periods to expand beyond the single-particle nature intrinsic to active-feedback tracking methods (*42*). Such prolonged live-cell imaging is made possible by a combination of continual laser scanning and viral motion, which reduces the overall laser dwell time at any single position. Multiple-trajectory registration enables 3D-TrIm to accumulate a population of trajectories sufficient to perform statistical analyses on the dynamics and activity of virions in a single area (fig. S39-40 and movie S8). We demonstrate this capability in fig. S39B, which shows the population average (27 of 38 trajectories) of VLP position and diffusivity, revealing different VLPs frequent similar areas of the cell surface and extracellular matrix. Further experiments may uncover what factors influence the heterogenous dwell time of virions to these particular regions of the surface.

The data collected here by the 3D-TrIm microscope represent a dramatic step forward for SPT at high speeds in complex systems. Prior to this study, the immobilization and detachment behaviors of single virions could be observed by epifluorescence microscopy (*13, 55*). However, here 3D-TrIm gives a continuous and uninterrupted timeline of the viral particle through the entire process, complete with volumetric imaging and environmentally contextualized diffusion analysis. While in the current study, 3D-TrIm is able to track viral particles for minutes at a time, following one particle through the entire infection process will require 10s of minutes of observation time. To address this in the future, we have recently developed an information-efficient sampling approach which extends the observation time dramatically by only sampling high-information areas around the particle (*56*). This should allow tracking of the VLP all the way from the extracellular to the perinuclear space.

The skimming events observed here are the first of their kind and demonstrate how cell morphology can affect viral diffusion and how diffusion is inhibited at the cell surface. For membrane-bound VLPs, while the phenomena of viruses utilizing actin-rich protrusions as tracks to facilitate transport along the plasma membrane has been well-described (*19, 23, 34-39*), the distinct transport modes revealed here give insight into the different ways that VLPs interact with cellular protrusions. Finally, we demonstrated never before reported rapid diffusion of single VLPs in a tightly packed epithelial layer, paving the way towards high-speed SPT in more realistic biological systems. Future studies with 3D-TrIm will be able to probe various unanswered questions in viral dynamics, including the “molecular walker” hypothesis (*57*) and the seeming impenetrability of the PCL for particles larger than 40 nm in diameter (*53*). This work is also readily extendable to the study of psuedotyped SARS-CoV-2 VLPs, which have already been shown to mimic many of the properties of the wild-type virion (*58*). Importantly, the application of this new technique can be extended to any system where fast dynamics of nanoscale objects occur over large volumetric scales, including delivery of nanoscale drug candidates to the lungs (*59*) and through leaky tumor vasculature (*60*).

## Supporting information

Movie S1. 3D-TrIm Method Animation Demo, related to Fig. 1.

Movie S2. VSV-G Exploring the extracellular matrix, related to Fig. 2

Movie S3. VSV-G Searching Behavior, related to Fig. 3A-C.

Movie S4. VSV-G Searching Behavior, related to Fig. 3E-G.

Movie S5. Virus Interaction with Filopodia, related to Fig. 4.

Movie S6. VSV-G diffusing through multi-layered epithelial cells, related to Fig. 5.

Movie S7. Internalized Viral Trafficking, related to Fig. S37.

Movie S8. Multi-Trajectory Overlay, related to Fig. S39.

## Acknowledgments

The authors acknowledge the Duke Viral Vector Core for assistance with virus-like particle generation and the Duke Cell Culture Facility for access to cell lines used in this study. The authors would also like to thank LSM Tech for help customizing the laser scanning microscope for integration into the 3D-TrIm setup. The authors are also grateful to Duke Office of Information Technology (OIT) and Duke Research Computing for facilitating access to Amira 3D 2021.1 for data rendering.

## Author contributions

Conceptualization: CJ, JE, KDW

Methodology: CJ, JE, KDW

Investigation: CJ, JE, YL

Data Curation: CJ, JE, YL, JA, KDW

Visualization: CJ, JE, YL, JA, KDW

Funding acquisition: KDW

Writing – original draft: CJ, JE, KDW

Writing – review & editing: CJ, JE, KDW

## Data and materials availability

Code to analyze single virus trajectories (MATLAB) and render them (Amira) is available at https://github.com/welsherlab/3dtrim. Raw data available from the corresponding author upon reasonable request.

## Supplementary Materials

Materials and Methods

Figs. S1 to S38

References (*61–71*)

Movies S1 to S8

