## Supplemental Text and Figures for "Capturing the start point of the virus-cell interaction with high-speed 3D single-particle tracking"

**This PDF file includes:**

Materials and Methods
Figs. S1 to S40
Captions for Movies S1 to S8

**Other Supplementary Materials for this manuscript include the following:**

Movies S1 to S8

### 1 Materials and Methods

#### 1.1 3D-TrIm Instrument Overview

The 3D-TrIm instrument design is shown in fig. S2 and consists of 3D-SMART tracking excitation optics and 3D-FASTR imaging excitation optics coupled through a commercial confocal microscope (Zeiss LSM 410, modified by LSM Tech), piezoelectric stage, and microscope objective to join both setups together.

##### 1.1.1 3D-SMART Overview

High-speed particle tracking is achieved using a rapidly scanning laser focus over a narrow field of view and single-photon counting detectors to calculate photon-by-photon position updates in real-time. This is accomplished using a pair of electro-optic deflectors (**EOD**) to scan the laser across a 25-point Knight's tour pattern (*44, 61*), covering a 1 μm$^2$ area in the XY-plane, holding at each spot for 20 μsec (Fig. 1B). A tunable acoustic gradient (**TAG**) lens (*62*) sinusoidally deflects the laser's focus at ~ 70 kHz over a 2 μm range around the focal volume (Fig. 1B). Photon arrivals at different laser positions are fed into a Kalman filter (*63*) on a field-programmable gate array (FPGA). The real-time localization is then used as the input to an integral-feedback controller, which controls a piezoelectric nanopositioner to recenter the particle in the objective focal volume.

##### 1.1.2 3D-SMART Excitation Optics

The optical pathway for 3D-SMART tracking excitation uses a continuous-wave solid-state 488 nm laser source (FCD488-30, JDSU) which is expanded 1.25× and spatially filtered with a 75 μm pinhole. The linear polarization of the beam is selected by a Glan-Thompson Polarizer (GTH5-A, Thorlabs) and rotated by a half-wave plate (WPH05M-488, Thorlabs). A pair of electro-optic deflectors (**EOD**) then deflect the polarized beam with a second half-wave plate in between to rotate the polarization orthogonal to the input. The deflected beam is further expanded 3.33× before focal modulation through an acoustic lens (**TAG lens**). This modulated focus is then relayed to the sample using three lenses that will preserve the input magnification. The beam is reflected into the underside of the LSM by a multiband dichroic mirror (**DCM2)** (Chroma ZT405/488/635rpc) and then reflected vertically toward the objective, collimated through a 200 mm lens (LA1708-A) situated in the underside of a slider containing a 700 nm shortpass dichroic (**DCM1**). The beam exits the slider and is focused by the objective lens.

##### 1.1.3 3D-FASTR Overview

In conventional laser scanning microscopes, a laser beam is raster-scanned point-by-point to form an image frame. Extension into the third dimension is achieved by translating the sample and performing serial frame acquisition at different depths. As the stage is used for active-feedback tracking of the particle, we instead used an electrically tunable lens (**ETL**) to create a continuous optical translation of the focus to provide an imaging depth range of up to 8 μm. This focal sweep completes several periods within the period of the frame raster scan, creating scan patterns across the Y-Z image plane which vary each frame-time. When the relative frequency between the ETL and the 2D frame scan are timed correctly (fig. S13F-H), a unique subset of voxel locations in the volume space are scanned each frame before the pattern repeats (Fig. 1C). This method, called 3D-FASTR, increases the volumetric imaging rate 4-8x beyond that provided by conventional volumetric microscopy methods because the resulting gaps between scanned voxels can be interpolated (*46*). The 1 µsec dwell time is common to most existing laser scanning microscopes and generates more than enough signal-to-noise to capture the morphology and dynamics of fluorescently labelled live cells.

The development of 3D-FASTR results in an imaging system capable of 3D imaging in motion, but since its original publication, the optical configuration of 3D-FASTR was re-designed to obtain a constant, diffraction-limited point spread function throughout the focal range. These optical improvements also resulted in a triangular focal translation without the need for arbitrary drive signals. Improvements to 3D-FASTR optical performance are discussed in section 1.5.1.

###### 1.1.4 3D-FASTR Excitation Optics

Two-photon excitation was enabled by incorporating a tunable-wavelength (tuned to 800 nm), pulsed Yb 100 fs, 80 MHz laser source (Chameleon Discovery, Coherent), and expanded 2.5× using a Galilean telescope (ACN254-050-B, Thorlabs; AC254-125-B-ML, Thorlabs). Variable focusing of the expanded beam was achieved using an electrically tunable lens (EL-10-30-C-NIR-LD-MV, Optotune) placed approximately 1200 mm behind the rear aperture of the LSM System. A 450 mm lens (**IL1**) (11665, Edmund Optics) positioned 935 mm after the ETL enabled nearly linear focal translation with respect to current (fig. S13D), eliminating the need for any external modulation described in (*46*). After passing through the confocal scan unit, the beam's focal range is adjusted by a 1000 mm lens (**IL2**) (LA1464-B, Thorlabs) placed on the imaging excitation side of the slider. The beam is then reflected vertically off a 700 nm shortpass dichroic mirror (**DCM1**) to the rear aperture of the objective lens (100x Plan-Apo 1.4 NA, Zeiss).

###### 1.1.5 Imaging Variable Focus Detection

The beam is sampled before variable focusing using a 10:90 beamsplitter (**BSR**) (BS025, Thorlabs). The sampled beam is focused using a 75 mm lens (AC254-075-B-ML, Thorlabs) onto a Si photodiode (**PDR**) (DET10A, Thorlabs) terminated with a 56 kΩ resistor. This photodiode is used as a power reference to normalize the signal. Two additional 10:90 beamsplitters (**BS1**, **BS2**) sample the beam after focal modulation at extrema of the ETL focal range, producing peak photodiode signals at focal length minima (**PD1**) and maxima (**PD2**). Because the two measurement photodiode signals peak out-of-phase with respect to focal shift, their difference produces a monotonic response curve with respect to ETL focal power as reported previously in (*46*).

###### 1.1.6 Shared Emission Pathway

A key factor to integrating the tracking and imaging modules in 3D-TrIm is minimization of crosstalk between the channels. In this implementation, tracking fluorescence collection occurs in the 500-600 nm window and Imaging fluorescence is collected in the 600-700 nm window.

Emitted fluorescence from both tracking and imaging passes back through the dichroic mirror (**DCM1**) and 200 mm lens on the tracking excitation side of the slider. Both emitted beams pass through the multiband entrance dichroic (**DCM2**) and focus through an 85 mm lens positioned approximately 360 mm from the objective lens. Reflected imaging laser excitation is removed through a multiphoton blocking filter (FF01-750/SP-25, Semrock), and the tracking and imaging emissions are separated with a second dichroic mirror (**DCM3**) (Di03-R594-t1, Semrock). The imaging emission is focused on to a non-descanned detector (**PMT**) using a 50 mm lens (LA1131-A, Thorlabs) positioned 50 mm beyond the previous 85 mm lens.

After DCM3, the tracking emission is relayed by a 300 mm lens (AC254-300-A-ML, Thorlabs), placed 625 mm beyond the previous 85 mm lens and focused, onto an avalanche photodiode (**APD**) by an 80 mm lens (AC254-080-A-ML, Thorlabs). This detection setup results in a beam magnification that nearly fills the APD sensor area and prevents interference from the scanned imaging laser.

##### 1.2 Tracking and Imaging Integration Overview

The 3D-SMART active-feedback tracking and 3D-FASTR imaging are coupled into a single platform through the chromatically separated excitation and emission pathways described above but unified by the master spatial grid of the piezoelectric stage. The stage coordinates define the physical space and can be used as an absolute reference between tracking and imaging spaces. As the 3D-SMART system natively outputs the trajectory in stage-space coordinates, only the location of the 3D-FASTR image must be adjusted to generate fully registered tracking and imaging volume spaces. Multiple calibrations were used to obtain the imaging parameters needed to convert voxel locations from image-space into physical stage-space and are described below.

###### 1.2.1 Tracking and Imaging Relative Center Positions

The lateral center of the tracking volume sits at a constant location relative to the imaging volume, determined by the relative alignment of the tracking and imaging excitation beams. If the two beams are perfectly co-aligned, the trackcenter pixel will be in the middle of the 512x512 image frame at (256,256). It is not necessary to perform this co-alignment, so long as the trackcenter is known as described below.

Fig. S6 shows the process used to register the relative location between the tracking volume in the image space. We used 200 nm multicolor microspheres (Tetraspeck, T7280, ThermoFisher) which produce a signal on both the tracking and imaging detectors, and were sparsely concentrated to a density of 1 sphere/imaging field of view. 2D Imaging data (fig. S6A) of the bead are acquired for ~100 frames while the bead is held in the focus of the tracking volume and the ETL focus is held constant at the plane of the axial track center. We assume that the position of the single tracked microsphere corresponds to the center of the tracking volume, and apply a Gaussian fit to find the peak center (fig. S6B). We repeat this procedure for each acquired frame and take the mean value for x/y as the pixel center and line center in the image frame. Fig. S6C-D show the distribution of Gaussian peaks.

The axial center is determined by the difference in collimation between the tracking volume and ETL volume. As tracking is initiated by acquiring the virus in the extracellular space above the cells, we expect the cells will enter the axial imaging range of the ETL scan from the bottom. To maximize the amount of cellular image sampling, we bias the focus of the tracking volume to be higher relative to the center of the imaging volume to more frequently image areas beneath the particle by moving the 400 mm lens farther than confocal distance, leading to a slightly converging behavior of the tracking beam. The net impact is with the tracking beam serving as the 0 μm reference, the axial depth range of the ETL scan extends 5 μm below the particle, and only 3 μm above it. This balances the need to sample the cell more aggressively on approach while maintaining 3D cellular imaging when particles are bound or internalized.

##### 1.2.2 ETL Calibration

The focal depth of each pixel in the image space must be known to assign imaging pixels into 3D volume space. We convert the photodiode signals to a focal depth in the image space using a calibration performed with the same multicolor 200 nm microspheres as previous which are visible to both tracking and imaging detectors. The microsphere is tracked for ~ 10 sec to obtain the focal position of the bead, given by the mean of the axial stage position which is used as the reference (0 μm) focal plane. After acquiring the reference plane, a series of image stacks and corresponding photodiode signals are acquired with the ETL held to a constant current. This process is repeated until the entire ETL current range is sampled. The resulting curve shape of intensity vs. focal depth is fit to an interpolant, the peak of which is taken to be the focal plane. This peak focal position and the average photodiode signal are taken for each sampled ETL current. The resultant curve of ETL relative focal shift vs. photodiode ratio difference (fig. S13E) is fit to a rational polynomial equation and used to convert photodiode signal to image space focal position.

##### 1.2.3 Pixel Size Determination

The image pixel size must be known to convert the pixel positions from pixel-space to stage-space. The pixel size was measured using a field of 6 μm microspheres imaged and then translated in 1 μm steps along a single axis of the piezoelectric stage after at least 1 frame-time of acquisition (1.07 sec). Fig. S5A shows the field of beads in image frame-space of frame 17 colored in green. The stage is translated and imaged each step and fig. S5B shows the resulting field at frame 71. The magenta-colored beads are visibly shifted from their original green location, apparent when the two frames are overlaid in fig. S5C. The distance each bead moves between frames is $\Delta_{pixel}$. This process was repeated across the 75 μm range of the stage and performed twice for each axis. The locations of the microspheres in the frame space were tracked using a circular Hough Transformation and distance minimization algorithm to identify each bead in every frame as shown in fig. S5D-E for frames 17 and 71. The relative change in image pixel or line centroid location versus the relative change in X or Y stage position yields a linear fit corresponding to the image size of the pixels in px/μm (fig. S5F-G). The slope direction for each axis is a function of the difference in orientation between the laser scan and the stage and the sign must be known and accounted for to convert the image from frame-space to stage-space.

##### 1.3 TrIm Data Acquisition Workflow

Tracking and imaging acquisition is initiated by the 3D-SMART laser pattern scanning through the tracking volume. When the number of detected photons exceeds a pre-determined threshold based on the laser power and observed background, a particle is assumed to be located within the tracking volume. This particle detection event will initiate the stage feedback loop, open the imaging laser shutter, and trigger data recording. Tracking and imaging data continues to be recorded until the tracking intensity drops below a threshold value based on the background level.

We note here that the primary limit to trajectory duration in diffusive scenarios is not photobleaching, but that trajectories most frequently ended when reaching the boundary of the piezoelectric stage. For confined trajectories, acquisition continues until bleaching causes the virus intensity to fall to a level similar to the background.

###### 1.3.1 Automated Acquisition Overview

Due to the serial nature of single particle tracking acquisition, obtaining a quantity of trajectories sufficient for population-level statistical analysis can be tedious and time-consuming. We developed an automated acquisition protocol to act as an autopilot so that 3D-TrIm experiments could be conducted without user input over experiment durations up to 12 hours. The cells were maintained at 37 °C and buffered (Live Cell Imaging Solution, Invitrogen A14291DJ) with 2% fetal bovine serum (FBS) (Millipore Sigma, #F2442) to preserve viability through the duration of the experiment.

The autopilot algorithm searches for particles and considers any particle-detection event as a trajectory regardless of duration. After a set duration, either time or quantity based, an area-migration event is triggered, which moves the motorized stage to a new location in an S pattern to minimize overall distance traveled. The algorithm then continues to search for particles in the new area and repeats this process until it is manually terminated, or an imposed time limit has been reached. A typical autopilot session results in acquisition of ~500 trajectories.

##### 1.4 TrIm Data Processing and Visualization Workflow

The goal of data processing in 3D-TrIm is to register tracking and imaging datasets in the same stage-space, which requires conversion of image data from pixel-space to stage-space. The pixel-based nature of 3D-FASTR acquisition results in image data without fixed temporal boundaries: any number of 3D volumes can be flexibly created from the total acquired image data, limited only by the desired balance between image quality and speed.

In the static-stage 3D-FASTR imaging case, the precise timing of the relative frame and ETL frequencies enables the out-of-sequence sampling to scan a volume completely without multiply sampling a voxel location more than once. The theoretical time to scan a complete volume, $T_{vol}$, is equivalent to the product of the frame time and number of z-slices in the volume. In 3D-TrIm, however, the motion of the stage across Z expands the volume beyond the initial size given by the ETL depth range alone. We observed axial diffusion ranges of up to 25 μm in a single $T_{vol}$ period. To account for increased volume size, we use the static-case $T_{vol}$ value as our base volume acquisition time.

Once the temporal volume boundaries are determined, the voxel positions are transformed from image frame position ($P_{image}$) into the stage coordinate space position ($V_{rel}$) in each axis using the calibrated value for the trackcenter ($P_{center}$) and the pixel size (Ψ, μm/px) according to the equation below which shows this transformation for the x-axis positions.

$$V_{X,rel.} = \left(P_{X,image} - P_{X,center}\right) * \psi_X \tag{1}$$

This value is then offset by the position of the stage, S, to yield the absolute position in physical space ($V_{X,abs.}$).

$$V_{X,abs.} = S_X \pm V_{X,rel.} \tag{2}$$

The sign is determined by the relative orientation between the piezo stage and laser scan as obtained through the slope of the pixel size calibration described above in section 1.2.3.

This process is simplified for the Z axis because the ETL calibration is zero-referenced to the axial trackcenter in microns, such that only the stage offset in equation 2 is necessary for voxel placement. After conversion, the stage-space voxel positions must be re-discretized to form the global voxel grid. This is accomplished by using the stage-space extrema as the centers of the resulting minimum and maximum voxels. The locations of these voxel boundaries are determined per trajectory based on the size of the global volume space.

After voxel assignment, each local volume is assessed for sampling to prevent over-interpolation of undersampled image planes and cropped such that the exterior planes of the volume are at least 20% sampled. The MATLAB (MathWorks) function `inpaintn` (*64*) is used to perform the inpainting operation. After interpolation, the local volume intensity is rescaled by setting the saturation point at the intensity equal to the cumulative distribution threshold of 0.9998 and contrast-enhanced by scaling with a gamma value of 0.7. Unless a time series is desired, these completed local volumes can be composited into a single maximum intensity projection (MIP) of the global volume over time. When displayed as a time series of local volumes, the image may appear incomplete or irregularly stripy in some regions. This occurs when the virion diffuses toward or away from the cells in Z such that cells are visible in the imaging volume for only a portion of the local volume time. Compositing into a global volume is recommended for forming distance maps to avoid such issues.

##### 1.4.1 Multi-Trajectory Processing and Visualization

For analysis and visualization of VLP dynamics across a single area, multiple trajectories residing in a common area were co-registered in a global volume space. The image data associated with each of the individual trajectories was accumulated and processed together as a single global image volume. As a result, minimal interpolation was required due to extended sampling time over the course of the many trajectories. The visualization of many trajectories was distinguished by applying a uniquely hued colormap to each visualized trajectory. The progression of these trajectories over time is shown through darkening of the base color.

#### 1.5 Method Validation Experiments

The following sections explain the methods used to validate 3D TrIm.

##### 1.5.1 Imaging Point Spread Function Measurement

The imaging point spread functions shown in fig. S13A-C were measured using the same image stack data prepared for calibrating the ETL as described above in section 1.2.2. To measure the lateral point spread function, peaks in the plane of highest intensity within a z-stack were fit to a multivariate Gaussian function. The widths of each peak were averaged to determine the FWHM for the lateral PSF at each ETL current. To measure the axial PSF, line cuts were taken at the location of each peak found previously and fit to a univariate Gaussian across the optical axis. As before, peak widths were averaged at each ETL current.

##### 1.5.2 Tracking Precision Validation

To validate our tracking precision, we tracked the same fixed VSV-G VLP over 5 consecutive trajectories of equal length (~ 28 sec). Due to photobleaching, the mean intensities (reported as photon counts per second, or Hz) of each trajectory varied from 100 kHz down to 40 kHz. We evaluated the tracking precision in these trajectories by normalizing the X, Y and Z positions and fitting the positions to a Gaussian (fig. S1A-C). At an average intensity of 100 kHz, the standard deviations are 0.025 μm in X, 0.024 μm in Y and 0.083 μm in Z, respectively. The standard deviation in all three axes increases as the intensities decreases. To better visualize and include more details of the relationship between standard deviation and intensity, we segmented the trajectories, plotting standard deviation as a function of average intensity for every 5 second segment of X, Y and Z, respectively (fig. S1G-I). The integrated intensity trace is shown in fig. S1J, in which the dashed line separates different trajectories and the dotted lines indicate the location of the segments.

##### 1.5.3 Registration Validation

To validate our registration between tracking and imaging we used a field populated with a diffuse spread of 200 nm Tetraspeck beads (fig. S4A). Using 3D-TrIm we acquired ~20 trajectories of microspheres located in the field by tracking the coverslip-bound beads for 30 sec while recording the image and then stepping the piezo stage beyond the previously tracked location to ensure a different microsphere was detected in the tracking volume. We evaluated our registration by fitting the region surrounding the calibrated trackcenter to a Gaussian (fig. S4B-C) and comparing the stage-space Gaussian centroid locations of the imaged beads versus the mean stage position of the bound particle trajectories.

The mean deviation between tracking and imaging centers was 100 ± 59 nm in X and 27 ± 24 nm, less than the lateral pixel size of ~180 nm (fig. S4D). The distribution for the axial deviation was skewed significantly such that the average deviation was 294 ± 109 nm. To correct this, we used this mean value as a calibration-offset and shifted all voxels by this amount. The distribution obtained using the shifted center reduced the mean deviation to 93 ± 59 nm, with both original and corrected distributions shown in fig. S4E.

##### 1.5.4 Spinning Disk Confocal Virus Tracking

Comparative virus tracking experiments to 3D-TrIm were conducted on a spinning disk confocal microscope (Andor Dragonfly 505), equipped with 100x/1.40-0.70 HCX PL APO (Leica 11506210) oil objective and 488 nm and 637 nm laser lines for excitation (40 µm pinhole). To analyze the ability of image-based tracking to capture viral diffusivity in the extracellular matrix of monolayer cells, volumetric imaging was performed over a comparable axial range to that of 3D-TrIm. HeLa cells plated on a coverslip, stained with 100 nM SiR650-actin (as described above) were positioned on a heated stage (37 °C) in a custom-built sample holder and maintained in a buffered solution (live cell imaging solution, ThermoFisher, #A14291DJ). eGFP.Vpr VLP was added to initiate the reaction to a final concentration of $1.9\times10^6$ TU/mL, which approximates to multiplicity of infection ~1.

Simultaneous two-color imaging was performed in dual-camera mode using a 570LP dichroic mirror, at 30 % and 2 % 488 nm and 637 nm laser power, respectively. Images were captured on an Andor iXon Life 888 EMCCD camera and an Andor Zyla PLUS 4.2 Megapixel sCMOS 512×512 with 2×2 binning, both with 40 msec exposure time per frame in the fastest interval mode. Each 8 µm volume comprised of 16 frames and completed on average every 2.32 sec for a total of 80 repeats.

Virus tracking on the spinning disk was also performed in 40 % glycerol (v/v) to determine the trajectory sampling rate in a slower diffusive environment. eGFP.Vpr VLPs diffused at 0.4-2 $\mu m^2/s$ over large axial ranges (>10 µm) in multi-layered epithelial cells. As a comparison the diffusivity of eGFP.Vpr VLP in 40 % glycerol at 37 °C was captured over a 10 µm z-range. Each volume comprised of 21 planes imaged an Andor Zyla PLUS 4.2 Megapixel sCMOS 256×256 with 2×2 binning at 40 msec exposure time per plane (30 % 488 nm laser power) in the fastest interval mode. Each volume completed on average every 2.37 sec for a total of 80 repeats.

##### 1.5.5 Spinning Disk Confocal Data Analysis:

Virus spots were tracked over time to create trajectories using the tracking algorithm in IMARIS software (Oxford Instruments). Briefly, spots in each frame were detected in the virus channel using the spot detection algorithm with and estimated XY Diameter of 0.25 µm and background noise subtraction. Detected spots were filtered based on the *Quality* filter value of 2.15. Spot in consecutive frames were connected using the *Brownian motion* tracking algorithm with a max distance of 3.73 µm and max gap size of 3, only trajectories longer than two frames are shown.

#### 1.6 Sample Preparation and VSV-G VLP Validation Experiments

For these single virus tracking experiments, we incorporated fluorescent protein into VSV-G VLPs by fusing either eGFP, or miRFP670, to HIV-1 Vpr which is packaged within the pseudovirus nucleocapsid. 3D-TrIm trajectories were acquired on cells labeled with SYTO61 (targeted to nucleic acids) or SiR650-actin (targeted to f-actin). The cell labels were chosen to maximize chromatic separation between the tracking and imaging, and the largest contributor to tracking crosstalk was cell autofluorescence by single-photon excitation, which was low enough not to perturb the active-feedback single-virus tracking. We cultured monolayers of either HeLa, BJ fibroblasts, or LDLR-deficient GM701 fibroblasts on glass coverslips. Alternatively, to achieve multi-layered cells, we cultured HT29-MTX cells on inverted matrix support filters (*65*). This selection of cell types offered diversity in morphology, extracellular environment, and cell surface receptor concentration to observe their influences on the early stages of viral contacts.

##### 1.6.1 Plasmid construction

The expression vector eGFP.Vpr, used to generate internally labelled VSV-G VLPs was constructed as follows. The vector pVpr+ backbone, was derived from mCherry-2xCL-YFP-Vpr (Addgene: #105215) and amplified by PCR with the following primers 5′-TTTTCCTGCAGGTGAACAAGCCCCAGAAGACC-3′ (SbfI site underlined) and 5′-TTTTACCGGTCGTCGACTGCAGAATTCG-3′. DNA encoding for the fluorescent protein, eGFP was amplified by PCR with the following primers: 5′-TTGCTAGCGCTACCGGTC -3′ (NheI site underlined) and 5′-TCCGCCTGCAGGCTTGTACAGCTCGTCCATG-3′ (SbfI site underlined) using eGFP-Rab7 (Addgene: #12605) as the template. Both PCR products were digested with NheI + SbfI and ligated together to create eGFP.Vpr, successful insertion was verified by sequencing.

The expression vector pmiRFP70.Vpr, used to generate internally labelled VSV-G VLPs was derived from the same backbone as above, and amplified by PCR with the following primers 5′-TTTTACCGGTCGTCGACTGCAGAATTCG-3′ (AgeI site underlined) and 5′-AAAACCTGCAGGTGAACAAGCCCCAGAAGAC-3′ (SbfI site underlined). DNA encoding for the fluorescent protein, miRFP670 was amplified by PCR with the following primers: 5′-GATCCACCGGTCGCC-3′ (AgeI site underlined) and 5′- TTTTCCTGCAGGGCTCTCAAGCGCGG-3′ (SbfI site underlined) using pmiRFP670-N1 (Addgene: #79987) as the template. Both PCR products were digested with AgeI + SbfI and ligated together to create pmiRFP670-Vpr, successful insertion was verified by sequencing.

##### 1.6.2 Production of VSV-G-pseudotyped Lentivirus Containing Fluorescently Labelled Vpr

Viral vectors were produced by the Duke Viral Vector Core Facility (Department of Neurobiology, Duke University School of Medicine), as described previously (*66*). Briefly, HEK-293T cells grown in 10 cm plates were transfected using a calcium phosphate-based protocol with either 10 µg of psPAX2, 2.5 µg of pREV, 5 µg pMD2.G (VSV-G), and 5 µg of the eGFP.Vpr plasmid (described above) or 10 µg of psPAX2, 2.5 µg of pREV, 5 µg pMD2.G (VSV-G), 15 µg of the transgene/reporter gene pLenti-GFP, and 5 µg of the pmiRFP670.Vpr plasmid (described above), to produce VLPs suitable for single-particle tracking or transduction assays, respectively. In both cases, the media was replaced at 24 h after transfection, and 72 h post-transfection, supernatant containing pseudotyped particles was centrifuged at 450 × g for 10 min and filtered through a 0.45-µm pore-size filter.

Viral titers were determined using a p24 -enzyme-linked immunosorbent assay (ELISA) and reported in transducing units/ml (TU/mL), typically 2-4×$10^7$ TU/mL. The virus solution was stored in aliquots at −80 °C. Prior to single-particle tracking and immunofluorescence experiments the virus solution was buffer exchanged by dialysis (Spectra-Por Float-A-Lyzer G2, MWCO 100 kDa, Millipore Sigma) against PBS (Genesee Scientific, Cat #: 25-507) at 4 °C.

##### 1.6.3 Infection Assays

To assess the infectivity of VSV-G pseudotyped VLPs containing fluorescent Vpr, HeLa cells were inoculated with VSV-G lentiviral vectors encoding the GFP reporter construct pLenti-GFP. HeLa cells (Duke Cell Culture Facility, ATCC # CRL-1958) were grown using complete DMEM which comprised of DMEM Media (Corning, #10-013-CV) supplemented with 10% foetal bovine serum (Millipore Sigma, #F2442), and 1× penicillin-streptomycin (Corning, #30-002-CI). Cells were maintained at 37 °C with 5% $CO_2$. 24 h prior to infection, cells were plated in complete DMEM at $5\times10^4$ cells/well in a 8 well μ-slide, glass bottom (Ibidi, #80827). Directly prior to infection, the media was exchanged with fresh media, where specified containing 8 μg/mL Polybrene (Santa Cruz Biotechnology, #SC-134220). HeLa cells were inoculated with a multiplicity of infection (MOI) of 0.01, 0.05, 0.1, 0.5, 1, 5, or 10 transfecting units per cell of lentiviral vector. The infected cultures were incubated at 37 °C in an atmosphere of 5% $CO_2$.

At 72 h p.i., cells were washed three times with Dulbecco's phosphate buffered saline (DPBS) (HyClone, #SH30264.01) and nucleic acid stained with 2.5 μM SYTO 61 (Life Technologies, #S11343) in live cell imaging solution (LCIS) (Life Technologies, #A14291DJ). The staining proceeded for 30 minutes in the 37 °C $CO_2$ incubator. The staining solution was removed, cells were washed successively three times with DPBS and mounted in LCIS.

HeLa-GFP cells were observed by wide-field fluorescence microscopy (Zeiss: Axiovert 200M) equipped a × 20 objective lens (Plan-NEOFLUAR, NA=0.5, Carl Zeiss) and EMCCD (QuantEM, 512SC) using Xeon Arc lamp with excitation at wavelength 480/30 nm and emission of wavelength >530 nm for GFP, wavelength 605/70 nm and >660 nm for SYTO 61. Nuclei and cell boundaries of GFP-positive cells were identified using the CellProfiler image analysis software (*67*).

##### 1.6.4 Antibody Labelling

To produce a labeled secondary antibody for immunofluorescence assays, 50 μg of goat anti-mouse IgG H&L (Abcam, #ab6708) was dialyzed against 1 L of 1× PBS at 4 °C for 4 h (D-tube mini, MWCO 6-8 kDa, Millipore Sigma, #71504-M). The volume of the recovered antibody was measured and to this 1/10 the volume of 100 mM NaHCO3 (pH = 8.3) was added. Next, to initiate labeling 5-fold molar excess of ATTO655-NHS ester (Millipore Sigma, #76245), dissolved and subsequently diluted in anhydrous DMSO, was added so that the final volume of DMSO was < 2%. The reaction was left to continue at room temperature for 1 h. The solution was applied to a desalting column (Zeba™ Spin, 7K MWCO, ThermoFisher, #89882) pre-equilibrated with 10 mM Tris-HCL, pH = 7.5, and 0.01% $NaN_3$, to terminate the reaction and remove uncoupled dye.

##### 1.6.5 eGFP.Vpr Quantification

Photobleaching experiments were performed on fixed VLP molecules to determine the number of eGFP.Vpr molecules incorporated within each virus-like-particle. VLPs were adhered to autoclaved quartz coverslips (EMS, # 72256-06) coated with poly-L-lysine (#P6282, Millipore Sigma), overnight in PBS at 4°C. Fixed eGFP.Vpr VLPs were held in the laser focus by engaging the tracking system. For photobleaching experiments, the 488 nm laser power was ~400 nW at the objective focus. The intensity level of a single eGFP chromophore was established by tracking the VLP until the intensity reached the background level.

##### 1.6.6 Immunofluorescence

The packaging efficiency of eGFP.Vpr inside the VLP was analyzed by two different immunofluorescence assays, one targeted against the inner capsid the other against the external envelope glycoprotein. VLPs were adhered to autoclaved glass coverslips (VWR, # 48380-046) coated with poly-L-lysine (#P6282, Millipore Sigma), overnight in PBS at 4°C. For HIV-1 p24 Gag labelling, VLPs were fixed for 20 min using 3.7% formaldehyde in PBS. Subsequently, washed (3×, 5min) and permeabilized for 10 min in T-PBS (0.1% Triton X-100 in PBS (pH=7.2)). Coverslips were blocked with buffer containing 10% normal goat serum (MP Biomedicals, #IC19135680), 0.2 M Glycine, and 0.1% Triton X-100 in PBS (pH=7.2) for 90 min at room temperature.

VLPs were stained for capsid protein using mouse anti-HIV-1 p24 gag monoclonal antibody (The following reagent was obtained through the NIH HIV Reagent Program, Division of AIDS, NIAID, NIH: Anti-Human Immunodeficiency Virus 1 (HIV-1) p24 Gag Monoclonal (#24-3), ARP-6458, contributed by Dr. Michael Malim), at 2 ng $\mu L^{-1}$ in blocking buffer 120 min at room temperature. Coverslips were washed (3×, 5 min) in T-PBS to remove any unbound primary antibody. For VSV-G labelling the sample was not fixed, instead the adhered LV particles were washed with PBS and blocked with PBS containing 10% normal goat serum (MP Biomedicals, #IC19135680), for 90 min at room temperature.

VLPs were stained for VSV-G envelope glycoprotein using mouse anti-VSV-G monoclonal (Kerfast, #EB0010), at 2 ng $\mu L^{-1}$ in blocking buffer 120 min at room temperature. Coverslips were washed (3×, 5 min) in PBS to remove any unbound primary antibody. In both scenarios as a secondary antibody, ATTO655-labelled Goat Anti-Mouse IgG (described above) was used at 4 ng $\mu L^{-1}$ in blocking buffer for 120 min at room temperature. Finally, coverslips were washed with either T-PBS or PBS (3×, 5 min), for p24 and VSV-G, respectively, and mounted in PBS. In control experiments the same procedure was followed except either the primary or secondary antibody incubation was omitted.

Immunostained VLPs were imaged on a spinning disk confocal (Andor Dragonfly 505) on a Leica DMi8 inverted microscope using 100x/1.40-0.70 HCX PL APO (Leica 11506210) oil objective and 488 nm and 637 nm laser lines for excitation (25 μm pinhole). Images were captured on an Andor iXon Life 888 1024×1024 EMCCD camera, and the system was controlled by Fusion 2.0. (Duke University Light Microscopy Core Facility NIH Shared Instrumentation grant 1S10RR027867-01).

Intensity based colocalization was performed using IMARIS software (Oxford Instruments) to extract the Pearson's coefficient for each field of view. Independently, eGFP.Vpr centers were identified in MATLAB (MathWorks) by determining the intensity maximum of each foci and the intensity at the corresponding position in the ATTO655 channel (*68*).

##### 1.6.7 Cell culture

HeLa (Duke Cell Culture Facility, ATCC #CRL-1958), fibroblasts (Duke Cell Culture Facility, BJ, ATCC #CRL-2522), hypercholesterolemia fibroblasts (Coriell Institute, GM00701), and mucus-producing differentiated goblet cells (HT29-MTX-E12, Millipore Sigma #12040401) were all grown using complete DMEM which comprised of DMEM Media (Corning, #10-013-CV) supplemented with 10% fetal bovine serum (Millipore Sigma, #F2442), and 1× penicillin-streptomycin (Corning, #30-002-CI). Cells were maintained at 37 °C with 5% $CO_2$ and passaged when they reached ~70% confluency.

##### 1.6.8 Cell staining and microscope sample preparation

24 h prior to microscopy, HeLa and fibroblast cells were plated in complete DMEM at $8\times10^5$ cells/well in a 6 well plate with an autoclaved glass coverslip (VWR, # CLS-1760-025) to ensure 80% confluency on the day of the experiment. HT29-MTX cells were seeded on the underside of a Transwell® permeable filter (Millipore Sigma, #CLS3460) in complete DMEM, which after attachment (4 h at 37 °C and 5% $CO_2$) was reinverted and cultured for 72 h prior to microscopy (*65*). Cells were stained with either nucleic acid stain (SYTO61, ThermoFisher, #S11343) or F-actin stain (SiR-actin, Siprochrome, #SC001). Cells were incubated with 2 μM SYTO61 in complete DMEM for 30 mins at 37 °C and 5% $CO_2$ followed by 3× wash steps (1XPBS). Alternatively, cells were incubated with 250-100 nM SiR-actin in complete DMEM for 6-14 hours at 37 °C and 5% $CO_2$ followed by 3× wash steps (1×PBS). In each case after staining the coverslip was transferred to HEPES pH=7.4 buffered solution (live cell imaging solution, ThermoFisher, #A14291DJ) in a custom-built sample holder and positioned on a heated stage (37 °C). All experiments were performed using live cells.

Live cell volumetric imaging and real-time viral tracking was performed on cells prepared as described above. eGFP.Vpr VLP was added to cells on a heated stage to initiate the reaction to a final concentration of $1.9\times10^6$ TU/mL, which approximates to multiplicity of infection ~1. The average excitation power at the focus was 180 nW and 7.5 mW for the 488 nm and 800 nm beams, respectively.

##### 1.6.9 Analysis of VLP size by 3D SMART

The size and brightness of eGFP.Vpr incorporated VLPs (fig. S9) was evaluated by real-time 3D tracking to extract the diffusion coefficient and particle emission rate. Free virus particles were tracked in HEPES pH=7.4 buffered solution (live cell imaging solution, ThermoFisher, #A14291DJ) at room temperature. The tracking microscope configuration was identical to that used for tracking and imaging as described in section 1.1.2. The average excitation power of the 488 nm laser was 180 nW at the focus.

#### 1.7 Data Analysis

The following sections describe protocols used to quantitatively analyze simultaneously acquired VLP trajectories and live cell imaging data.

##### 1.7.1 Mean Square Displacement Analysis

The diffusion coefficient can be obtained by linear fitting of the mean square displacement (MSD) with lag time (τ). The MSD was calculated using the definition:

$$MSD(\tau) = N^{-1}\Sigma_{n=1}^{N}\big((x(n+\tau) - x(n))^2) + (y(n+\tau) - y(n))^2) + (z(n+\tau) - z(n))^2)\big) \quad (3)$$

Here, $x(n)$, $y(n)$, and $z(n)$ are the coordinates of the trajectory at timepoint $n$. $N$ is the total number of data points, here the entirety of each change-point segment was used. For segments shorter than 30 msec the diffusion coefficient was calculated from the sum of the variances in the x, y, and z coordinates step size. When available, the 95% confidence interval of the diffusion coefficient was calculated from the linear fit of MSD($\tau$) using the MATLAB (MathWorks) function `confint`. Alternatively, for diffusion coefficients of segments less than 30 msec, the error was estimated using the variance of the variance:

$$D_{unc} = \frac{1}{2\tau}\sqrt{\frac{2}{N}}\left(\sigma_x^2 + \sigma_y^2 + \sigma_z^2\right) \tag{4}$$

For trajectories defined as diffusing on cellular protrusions MSD analysis was conducted using only the steps along the fit cylinder z coordinates so that confinement to the cylinder surface does not affect the calculation of D (section 1.7.7).

For single virus tracking only experiments, without cells, the hydrodynamic radius (*r*) of the particles was calculated using the Stokes-Einstein relation:

$$D = \frac{k_B T}{6\pi\eta r} \tag{5}$$

Here, $k_B$ is the Boltzmann constant, *T* is temperature, and $\eta$ is viscosity of solution.

##### 1.7.2 Change Point Analysis

A Gaussian changepoint algorithm was used to extract diffusive changes within a single virus trajectory (*69*). The full method can be found in the work of Montiel and Yang. Here we briefly overview the process. To minimize the calculation time and avoid interference from the piezoelectric feedback the sampling rate of the trajectory was downsampled to 3 msec. In a recursive manner, a log-likelihood ratio test is used to identify change points in a trajectory by separating the trajectory into different segments and calculating the following test statistic $\lambda_N$:

$$\lambda_N = \sqrt{max\left(Nlog(\sigma_N^2) - klog(\sigma_k^2) - (N-k)log(\sigma_{N-k}^2)\right)} \tag{6}$$

Here, $N$ is the total number of data points, $k$ is the point at which the trajectory is split, $\sigma_N^2$ is the step-size variance for the entire trajectory, and $\sigma_k^2$ and $\sigma_{N-k}^2$ are the variances before and after the split point, respectively. The split point $k$ is cycled through all the data points to find the maximum value of the test statistic. This value is compared to the approximate asymptotic critical values derived by Horvath (*70*). A false-positive error of 10% was applied. If the test statistic exceeds the critical value, a change point is identified and the trajectory split into two segments. Changepoint analysis is then run on each segment until no more changepoints are identified. Diffusion coefficients within each segment are determined via mean-squared displacement, as detailed above.

##### 1.7.3 Protrusion Cylindrical Fit

Cylindrical fitting was performed following Enkhbayar et al. (*71*). Briefly, the following function was minimized:

$$f = \Sigma_i(|\vec{x_i} - \vec{o} - (\vec{x_i} \cdot \vec{a})\vec{a}| - r)^2 \quad (7)$$

where $\vec{x_i}$ are the 3D coordinates, $\vec{o}$ is a vector from the origin to the cylindrical axis, $\vec{a}$ is the cylinder axis, and $r$ is the radius of the cylinder. The MATLAB function `fmincon` was used to minimize the function $f$ given the following constraints:

$$\vec{a} \cdot \vec{o} = 1$$
$$|\vec{a}| = 1$$

##### 1.7.4 Cell Boundary Determination

Image segmentation of the cell boundaries was required for calculating virus-cell distances. Membrane stains were considered as a method for identifying the edge of the cell, but the continual recycling of the membrane by the live cells led to a complete internalization of any membrane dye within 30 minutes of the start of the experiment. SYTO (which labels the cytosol and nucleus through DNA/RNA staining) and SiR Actin were stable and reliable indicators of the cell morphology.

Indeed, while the membrane itself was not stained during these experiments, it can be reasoned that the actin filaments and nucleic acid-containing cytoplasm, both types of stains used, sit directly beneath the membrane and their thickness is significantly less than the resolution of our imaging. Given this, it can be reasonably assumed that our visualized cellular images approximate the morphology of the cell within 500 nm.

To segment the cell boundaries in the image, we utilized the morphological top-hat filter with a spherical structuring element of radius 5 pixels using the `imtophat` function in MATLAB (MathWorks) and applying this transformation to the local volume intensity described in section 1.4. This function retains elements smaller than the structuring element and brighter than the background. To remove segmented noise pixels, we employed a second morphological filter, `bwareaopen` in MATLAB, which removes small unconnected objects. Finally, the segmented volume is passed to the `isosurface` function in MATLAB to determine the faces and vertices of the resulting segmentation.

##### 1.7.5 Cell-to-Virus Distance Calculation

To determine the minimum distance, *D*, between each point, *i*, in the cellular image, *c*, and the viral trajectory, *v*, the volume must be segmented using the top hat procedure described above. Then, for each segmented voxel, the distance is compared to every point of the trajectory. The minimum Euclidean distance shown in the formula below is taken as the distance to the nearest point on the trajectory to that cellular voxel.

$$D_i = \min_j\left(\sqrt{\left(c_{x,i} - v_{x,j}\right)^2 + \left(c_{y,i} - v_{y,j}\right)^2 + \left(c_{z,i} - v_{z,j}\right)^2}\right) \quad (8)$$

An illustration of this calculation is shown in fig. S12 where the segmented cell is visualized as a gray surface. The red, orange, and blue spheres represent selected locations of the voxels ($c_{xyz}$). The purple vectors draw a line to the calculated closest point of the trajectory, $\tau_{xyz,min.}$. The spheres are color-mapped based on the calculated minimum distance; only unique contact points are shown. Contact events were defined using the calculated minimum distance as distances within 0.5 μm of the segmented cell surface, on the same size scale as the imaging Z-PSF. The contact events were visualized as purple patches on plots of distance as a function of time (Fig. 2C). However, to aid visualization of these data, these regions correspond to rolling average (50 msec) distance measurements. The distance volumes (such as those shown in Fig. 2B) were created by rendering the segmented cell surface with color mapping by trajectory position to cell-surface distance. These calculated distances, in conjunction with diffusivity calculations, were used to visualize and quantify the nature of viral interaction with the cell surface.

##### 1.7.6 Determination of Trajectory fate

Active-feedback trajectories can be terminated for several reasons. As described above, the particle may diffuse out of the range of the piezoelectric stage, or its intensity may drop due to photobleaching below a threshold set near the background intensity level. Additionally, a brighter particle may diffuse into the tracking volume causing the algorithm to neglect the original dimmer particle and start following the brighter particle. The fate of each trajectory after any of these events occurred was manually determined for a subset of trajectories lasting longer than 20 sec.

Events were determined using a combination of the particle intensity trace (I), diffusivity trace (D), coordinate traces, distance trace, and image intensity, and categorized into the five behaviors below:

| | | | |
|---|---|---|---|
| (1) | Remain freely diffusing | $D \geq 1$ μm$^2$/s, | Any distance |
| (2) | Begins diffusive, binds to cell | $D_{bound} < 10\times D_{free}$ | Dist. < 0.5 μm |
| (3) | Tracking initiates on a surface-bound particle (stationary) | $D \leq 0.1$<br>random, confined motion | Dist. < 0.5 μm |
| (4) | Tracking initiates on a surface-bound particle (mobile) | $1.6\times10^{-4} < D \leq 0.05$<br>linear motion | Dist. < 0.5 μm |
| (5) | Starts diffusive, jumps to bound particle | $D_{bound} < 10\times D_{free}$,<br>$I_{bound} > 1.5\times I_{free}$, | Dist. < 0.5 μm |

The results of this classification are shown in fig. S38.

##### 1.7.7 Volume Render Visualization

3D visualization of intensity and distance volume data was performed using Amira 3D 2021.1. Binarized arrays with registered spatial coordinate data were imported into the program and visualized using cubic interpolation and alpha composition. Alpha mapping was controlled through a gamma curve to reduce the opacity of noisy features. Features were rendered solidified using an opacity threshold value which solidifies only surface regions and not background, typically by using a value of ~ 0.2. Trajectories were imported as comma-delimited spreadsheets containing coordinate, time, and diffusivity data and displayed as Line Set view objects with a line width of 2.

#### 2 Figures and movies

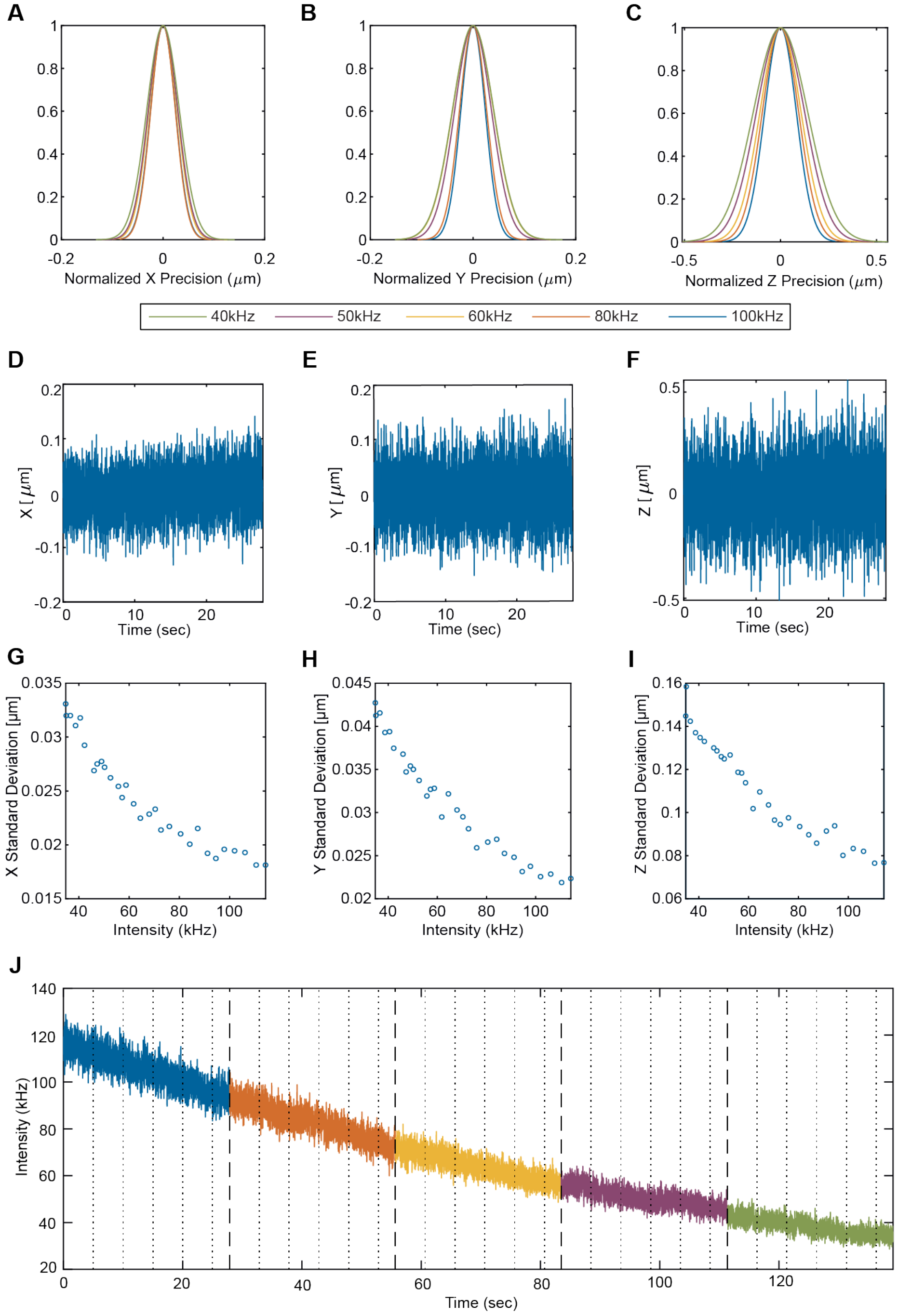

**Fig. S1. Tracking Localization Precision**

Tracking precision was calculated from 5 datasets of the same fixed particle. (**A**, **B** and **C**) Gaussian fit of the normalized X, Y and Z position in different mean intensities (reported as photon counts per second, or Hz). (**D**, **E** and **F**) Representative dataset of the normalized X, Y and Z position of fixed particle. (**G**, **H** and **I**) Standard deviations of X, Y and Z position readouts (in 5 sec segments) increase as intensity decreases, indicating the tendency of losing tracking precision due to the decreasing in intensity. (**J**) Integrated intensity trace of 5 chosen datasets. The line colors show different mean intensity. Dashed lines separate different datasets while dotted lines indicate the 5 sec segmentation in (G-I).

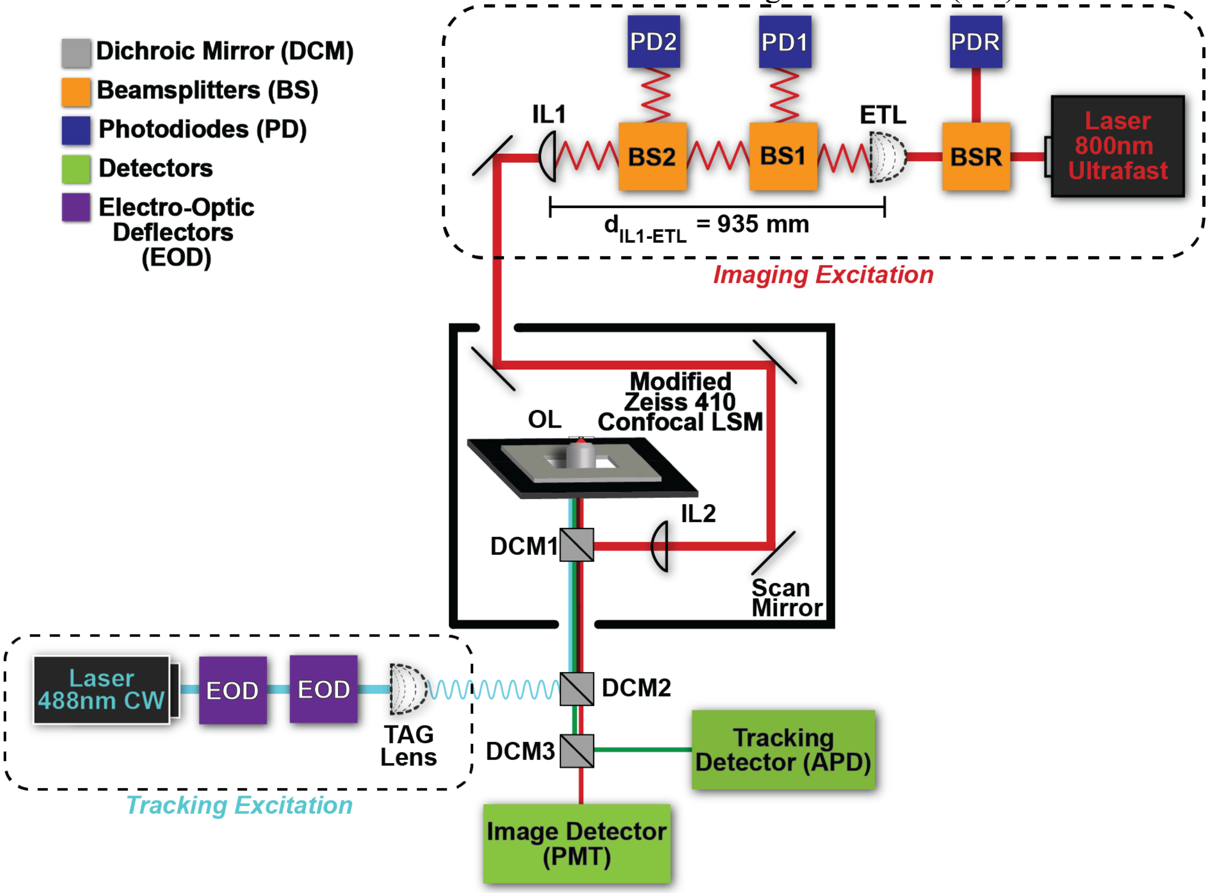

**Fig. S2. Instrument Diagram**

3D-TrIm Instrument Diagram. The 3D-TrIm microscope consists of two excitation sources: one for single-particle tracking and one for imaging. The tracking laser position is modulated using two electro-optic deflectors (**EOD**) and tunable acoustic gradient (**TAG**) lens. The imaging laser focus is modulated by an electrically tunable lens (**ETL**) and the beam is sampled by beam-splitters (**BS**) and evaluated using a system of photodiodes (**PD**). This focus is relayed using two lenses (**IL**). The tracking laser enters the underside of the LSM410 through a dichroic mirror (**DCM2**) where it ultimately couples with the imaging laser through **DCM1** toward the shared objective lens and piezoelectric stage. After exciting the sample, the fluorescence emission pathway is shared between tracking and imaging until separated by **DCM3**.

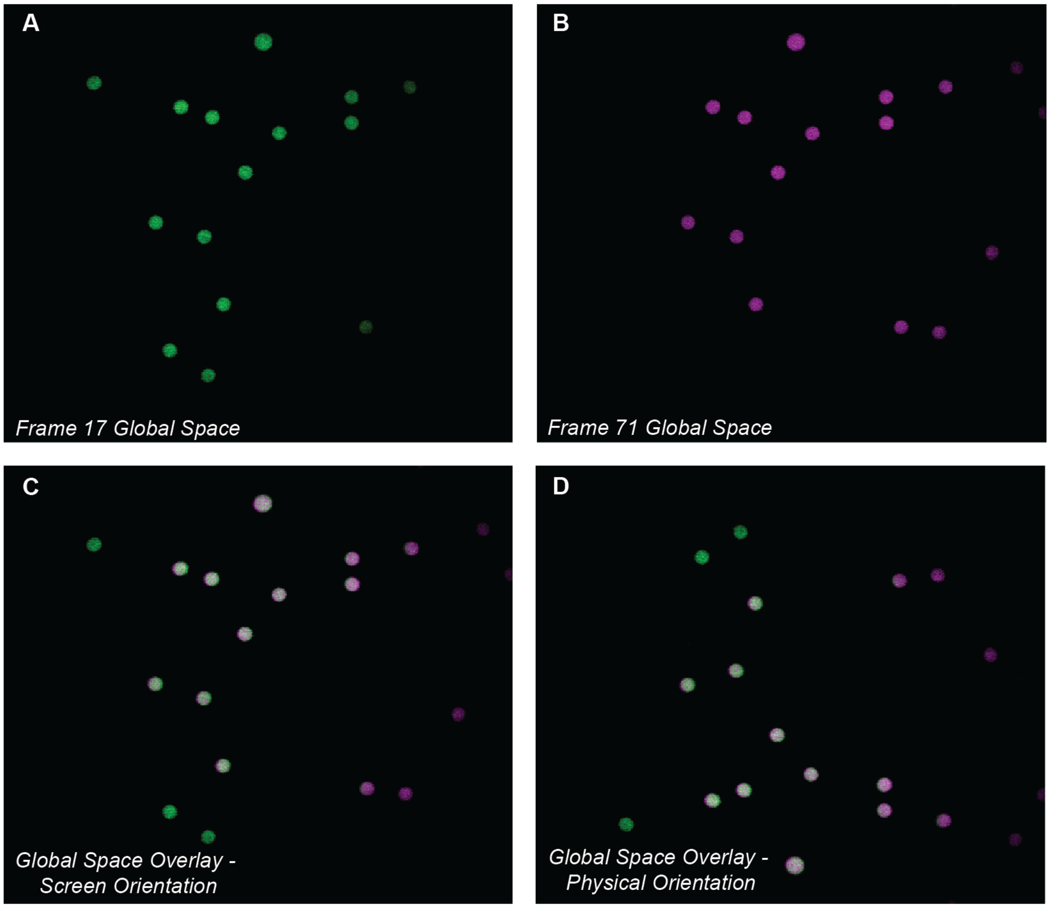

**Fig. S3. Registration Concept**

Registration Concept using 6 µm fluorescent microspheres. This concept involves moving the stage between imaging periods and registering the pixels in the absolute global volume space, instead of the frame space (see: fig. S5). (**A**) Frame 17 registered in global space. (**B**) Frame 71 registered in global space. (**C**) Overlay of global volume space frame 17 and 71. Unlike in the previous overlay, there is no apparent shift due to the registration, rather each frame occupies a different portion of space, so the extrema feature microspheres not shown in the other. This image is reflected and rotated for easy comparison to the originals (**D**) Shows the true orientation in physical space. This is the image orientation that results from registration.

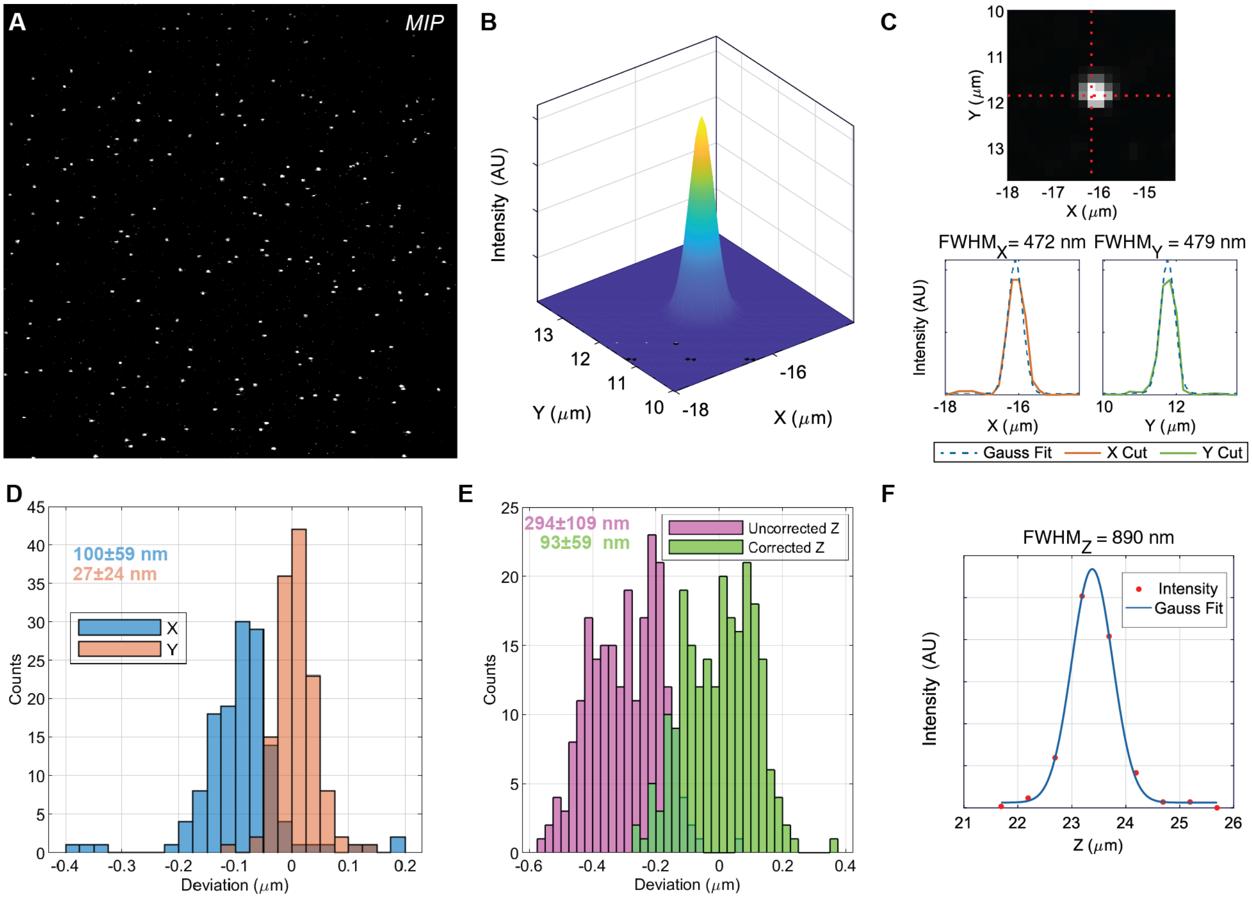

**Fig. S4. Registration Validation**

The process for validating our registration between tracking and imaging involves acquiring 3D-TrIm datasets using a field of microspheres which detect on both tracking and imaging channels. (**A**) Maximum intensity projection (MIP) of microsphere. **(B)** Gaussian fitting near trackcenter to obtain bead centroid in stage-space. (**C**) Fit across Z-axis cutting through peak in (**B**) to find 3D centroid. (**D**) Histogram of the difference between stage position and image centroid (registration error) in X/Y. (**E**) Histogram of Z registration error before/after correction.

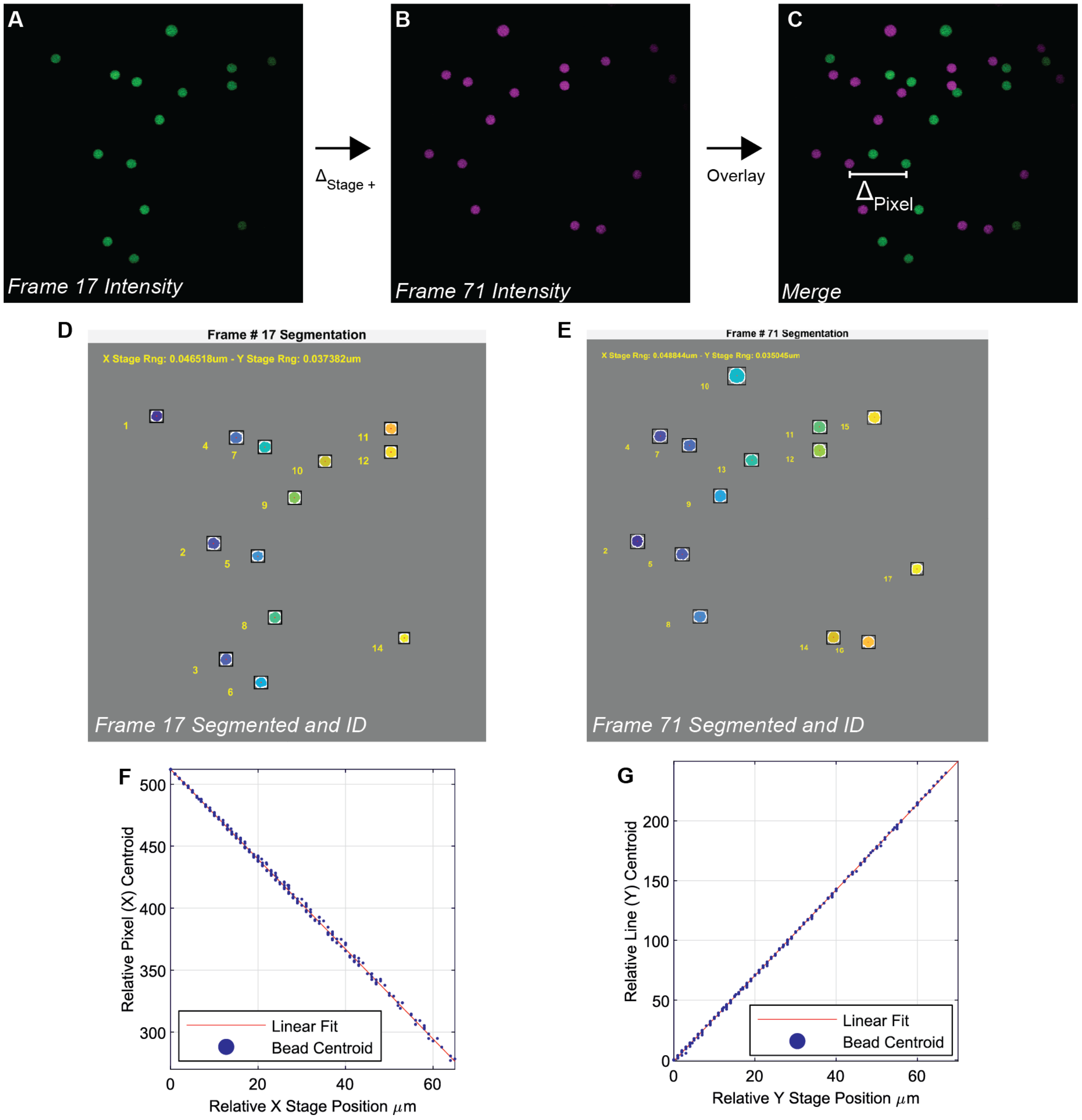

**Fig. S5. Pixel Size Calibration**

Pixel size calibration using image-based tracking of 6 µm fluorescent microspheres (from the same imaging data shown registered in fig. S3). The stage is translated 1 µm between frame acquisitions. (**A**) Intensity of frame 17. (**B**) Intensity of frame 71. (**C**) Frame-space overlay of frames 17 and 71 show displacement along the translation axis. (**D**) Segmented and numbered microspheres on frame 17. (D) Segmented and numbered microspheres on frame 71 shows ability to consistently identify and number beads despite shift. (**F**) Plot of relative pixel displacement versus relative stage displacement along the X-axis. Using relative values enables measurement of beads that are initially off-screen at the stage position of the first frame. (**G**) Plot of relative pixel displacement versus relative stage displacement along the Y-axis.

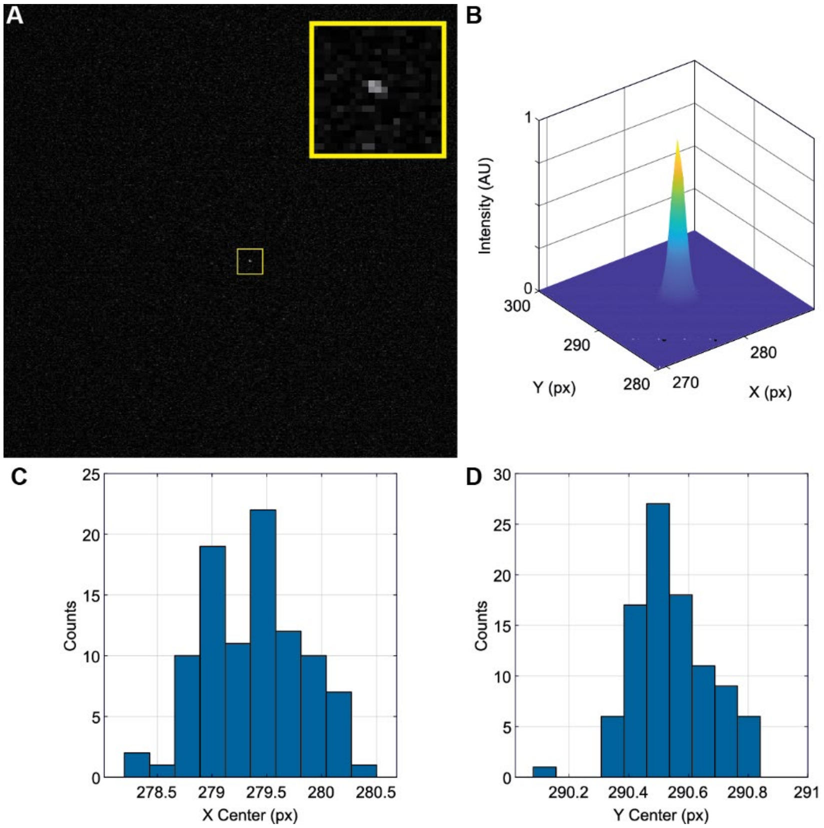

**Fig. S6. Trackcenter Calibration**

TrIm trackcenter registration calibration (**A**) 2D frame image with single microsphere located at center of tracking volume. Inset shows magnified view. This snapshot shows the microsphere being actively tracked and held it in the center of the tracking volume, therefore, detected simultaneously on both tracking and imaging systems. (**B**) Gaussian fitting of intensity is used to find trackcenter location (**C**) Histogram of X centroid coordinates (**D**) Histogram of Y centroid coordinates.

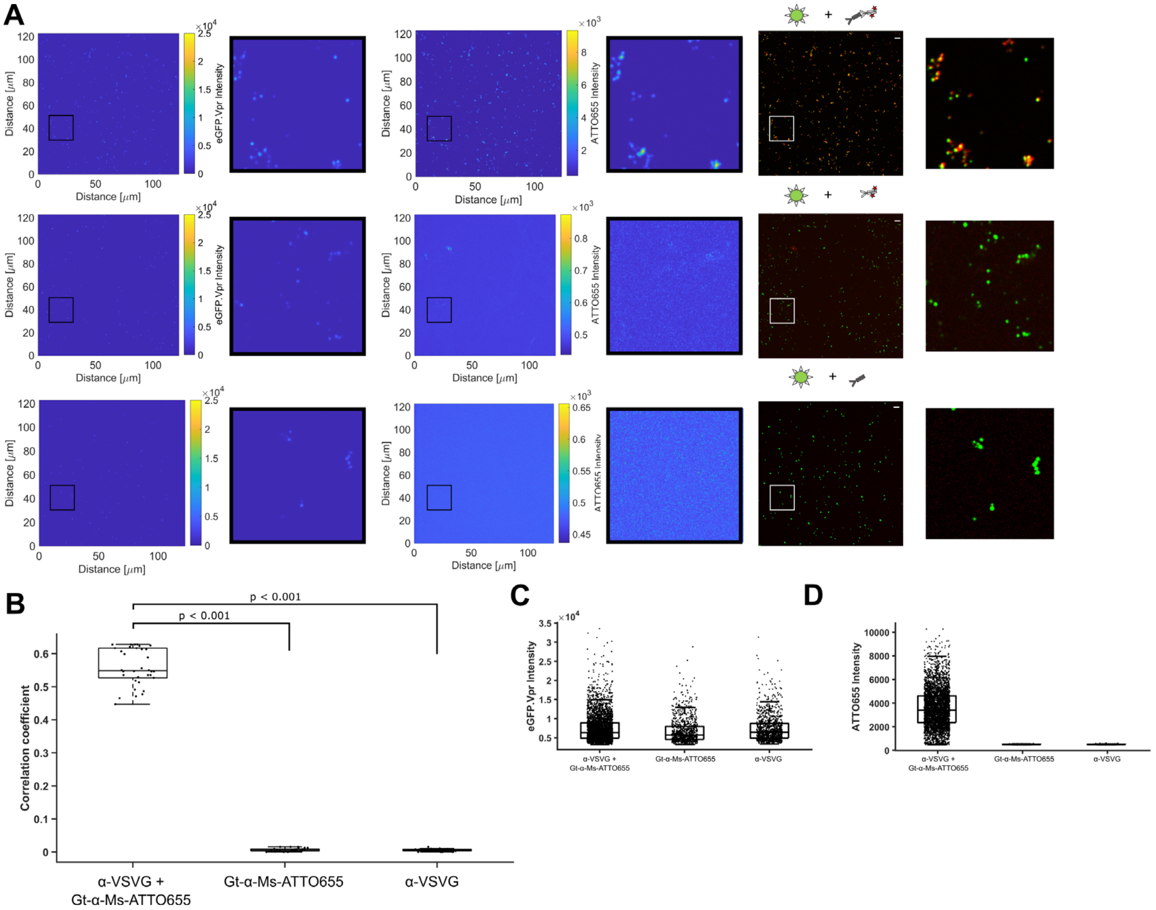

**Fig. S7. eGFP.Vpr is incorporated into VSV-G LVs, assessed by anti-VSV-G immunofluorescence.** (**A**) Left: representative raw intensity images of eGFP.Vpr -VSV-G VLPs. Middle, representative raw intensity images of ATTO655-labled VSV-G. Right: false-colored merge of eGFP.Vpr (green) and VSV-G (red). Top panel, immobilized VLPs incubated with primary and secondary antibodies. Middle panel, negative control, immobilized VLPs incubated with secondary antibody only. Bottom panel, negative control, immobilized VLPs incubated with primary antibody only. (**B**) Intensity-based colocalization analysis, yielding Pearson's correlation coefficient, was perform on each field of view (FoV), representative FoVs shown in (A) (n = 30). Statistical significance was assessed by Kruskal-Wallis test. (**C**) Intensities of eGFP.Vpr spots and the corresponding ATTO655 spot intensity at the same pixel location shown in (**D**). Standard box plots are provided in (B), (C), and (D), with the center representing the median and the top and bottom edges showing the third and first quartiles of the data. The whiskers encompass data points not considered outliers.

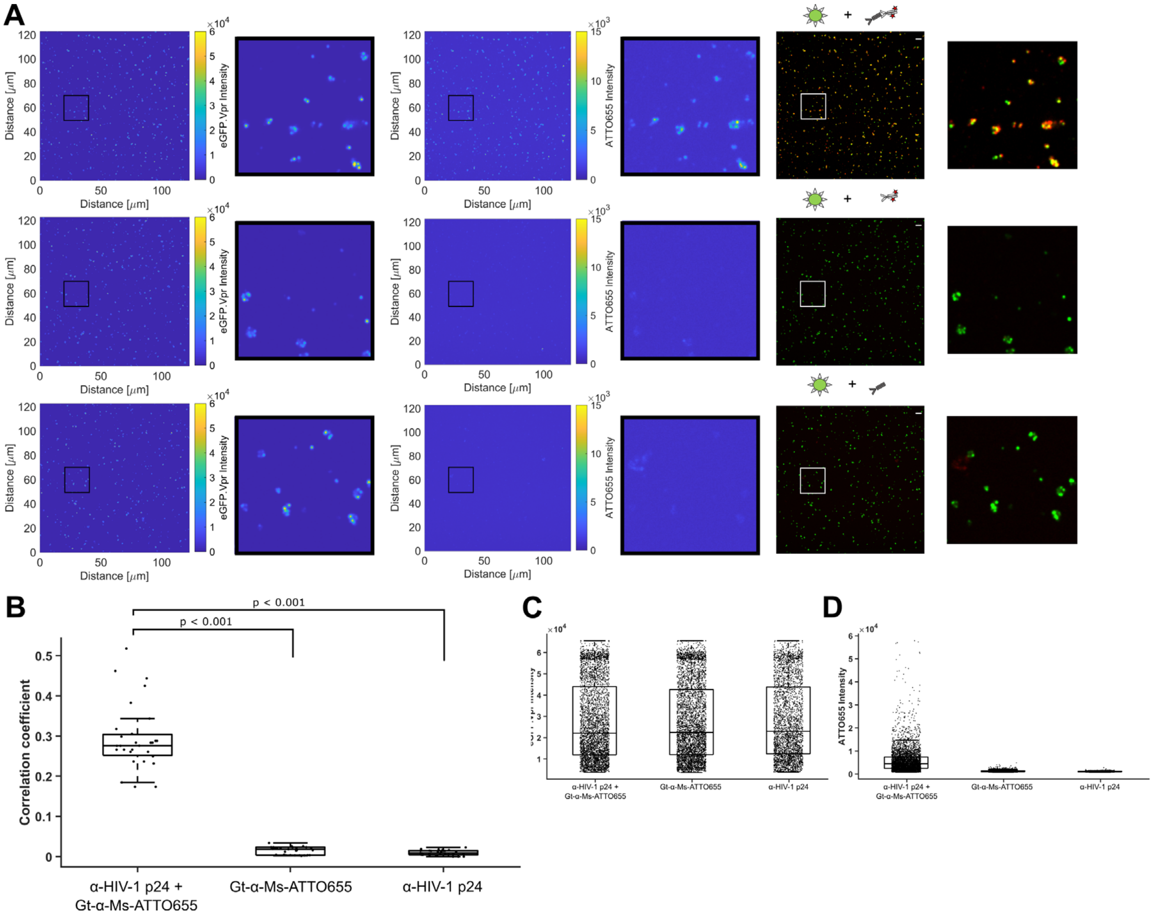

**Fig. S8. eGFP.Vpr is incorporated into VSV-G capsid, assessed by anti-gag immunofluorescence.**

(**A**) Left: representative raw intensity images of eGFP.Vpr -VSV-G VLPs. Middle, representative raw intensity images of ATTO655-labled p24. Right: false-colored merge of eGFP.Vpr (green) and VSV-G (red). Top panel, immobilized VLPs incubated with primary and secondary antibodies. Middle panel, negative control, immobilized VLPs incubated with secondary antibody only. Bottom panel, negative control, immobilized VLPs incubated with primary antibody only. (**B**) Intensity-based colocalization analysis, yielding Pearson's correlation coefficient, was perform on each field of view (FoV), representative FoVs shown in (A) (n = 30). Statistical significance was assessed by Kruskal-Wallis test. (**C**) Intensities of eGFP.Vpr spots and the corresponding ATTO655 spot intensity at the same pixel location shown in (**D**). Standard box plots are provided in (B), (C), and (D), with the center representing the median and the top and bottom edges showing the third and first quartiles of the data. The whiskers encompass data points not considered outliers.

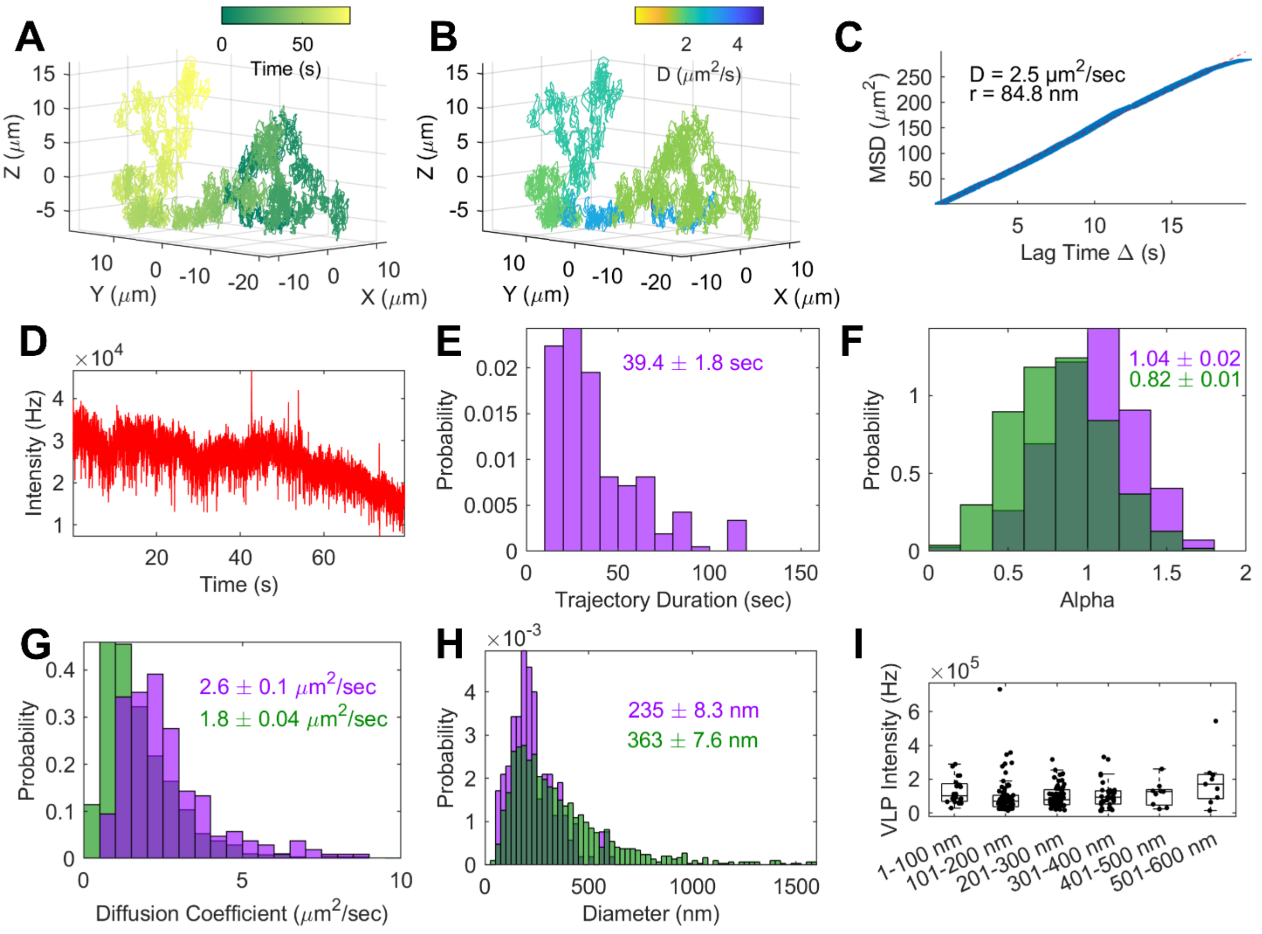

**Fig. S9. Sizing of VLPs via MSD and change-point analysis.**

(**A**) Complete trajectory of free virus. (**B**) Trajectory in (A) separated into distinct diffusive states via change-point analysis. (**C**) Mean squared displacement analysis with measured MSD (blue solid line) and linear fit (red dashed line) of trajectory in (A) yielding a measured hydrodynamic radius of 84.8 nm. (**D**) Tracking intensity trace of virus in (A). (**E**) Average trajectory duration (**F**) Free viruses undergo Brownian diffusion; alpha represents power component of mean squared displacement analysis. (**G**) Diffusion coefficient. (**H**) Hydrodynamic diameter. (F-H, magenta) full trajectory MSD-analysis, (G-I, green) change-point analysis (mean ± SEM, n = 210). (**I**) Vpr.eGFP intensity does not correlate with virus size, diameter from MSD analysis of full trajectory. A standard box plot is provided with the center representing the median, and the top and bottom edges showing the third and first quartiles of the data. The whiskers encompass data points not considered outliers.

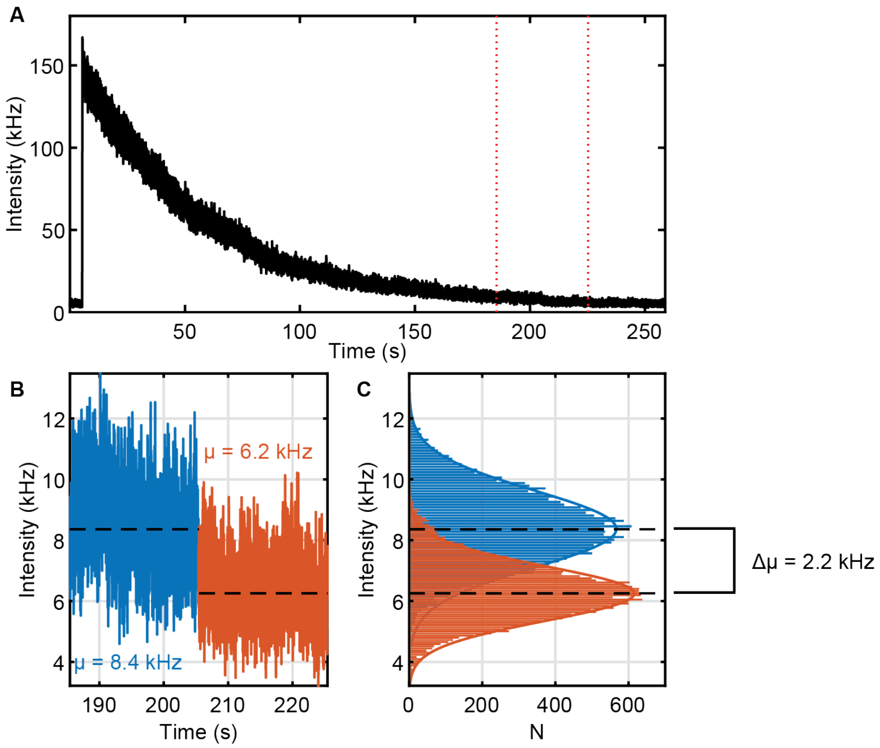

**Fig. S10. Determination of the emission intensity of a single eGFP-Vpr.**

(A) Intensity trace of a eGFP-Vpr VLP immobilized on a glass coverslip. (B) Zoom in on the final bleaching step, with the intensity before the bleach shown in blue and the background intensity shown in orange. (C). Histrogram of the intensities from (B), showing the magnitude of the final bleaching step to be 2.2 kHz for a single eGFP-Vpr.

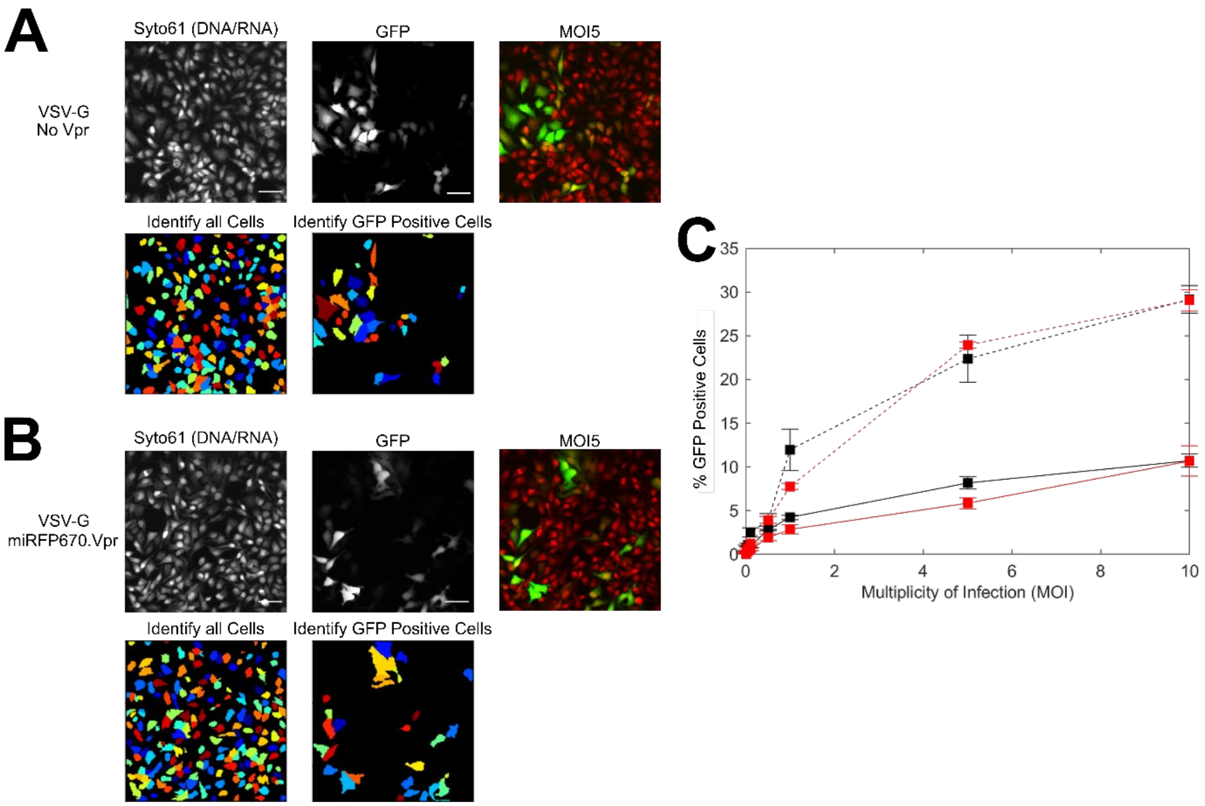

**Fig. S11. Fluorescent-Vpr does not prevent VSV-G gene delivery.**

GFP expression in HeLa after infection with VSV-G carrying GFP-reporter gene, either (**A**) without encapsulated Vpr, or (**B**) with encapsulated miRFP670-Vpr. HeLa cells stained with Syto61, 72 h-posttrasnduction at MOI 5 (top panel). The total number of cells and only cells expressing eGFP were detected using CellProfiler analysis software (bottom panel). Reporter assays were performed using the 20× magnification objective. Scale bar = 50 µm. (**C**) Average ± SEM (n = approximately $3 \times 10^4$ cells) of the percentage of cells expressing eGFP after transduction with VSV-G No Vpr (black) or VSV-G miRFP670.Vpr (red), in the presence (solid) and absence (dashed) of polybrene.

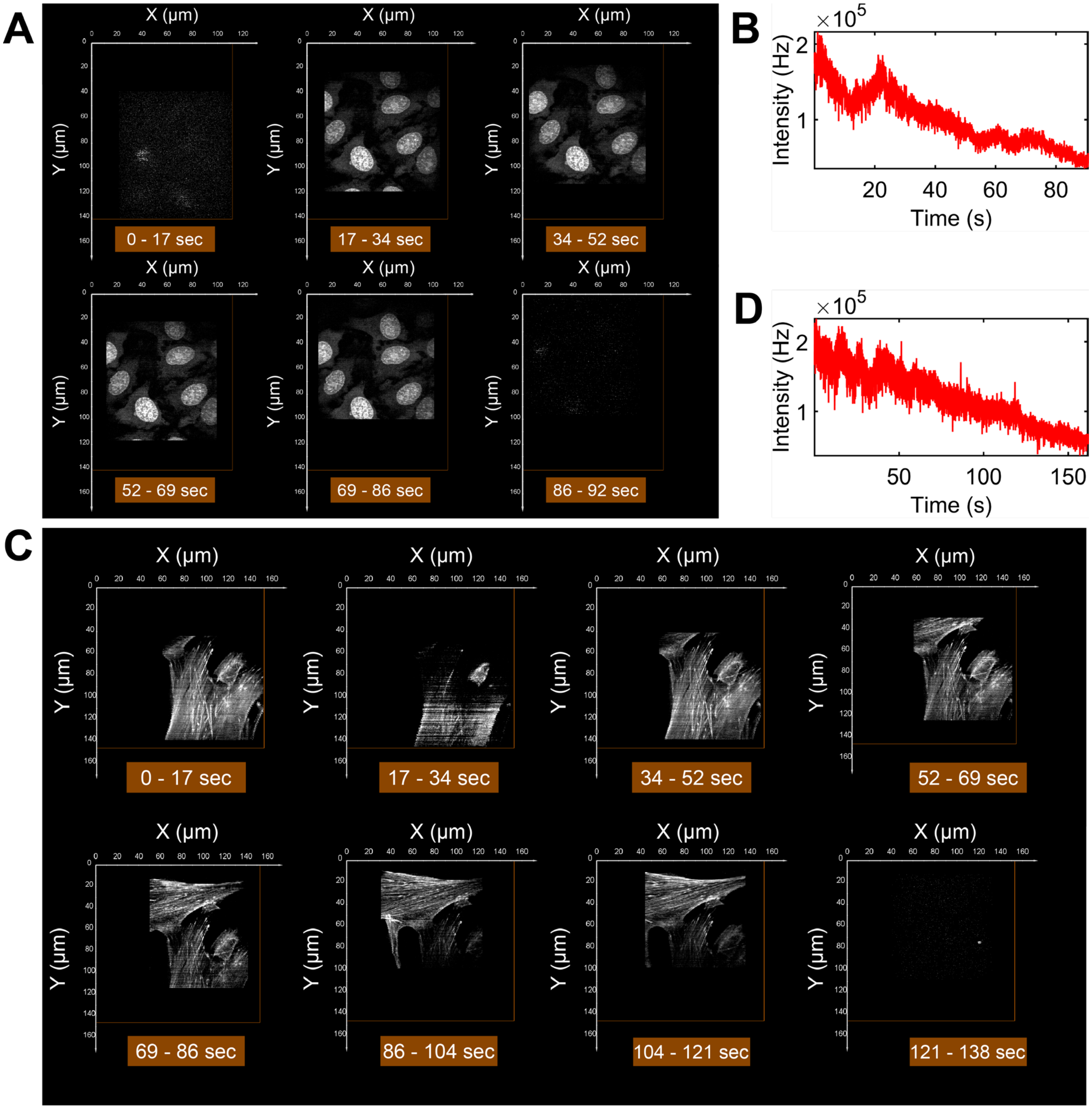

**Fig. S12. High resolution tracking and 3D imaging data, related to Fig. 2.**

(**A**) Local Volume MIPs. (**B**) Tracking intensity trace. (**A** and **B**) associated with VSV-G/HeLa tracking and imaging render, Fig. 2. (**A-E**). (**C**) Local Volume MIPs, local volumes from 138-162 sec have been omitted as trajectory diffuses a greater distance above the cells than the axial range of the ETL. Note in volume 2 from 17-34s there is a noted absence of intensity due to transit in Z of the particle away from the cells. This can appear as striping due to unsampled voxels and is a feature of local volumes, especially when displayed as MIPs . (**D**) Tracking intensity trace. (**C** and **D**) associated VSV-G/GM701 tracking and imaging render, Fig. 2. (**F-I**).

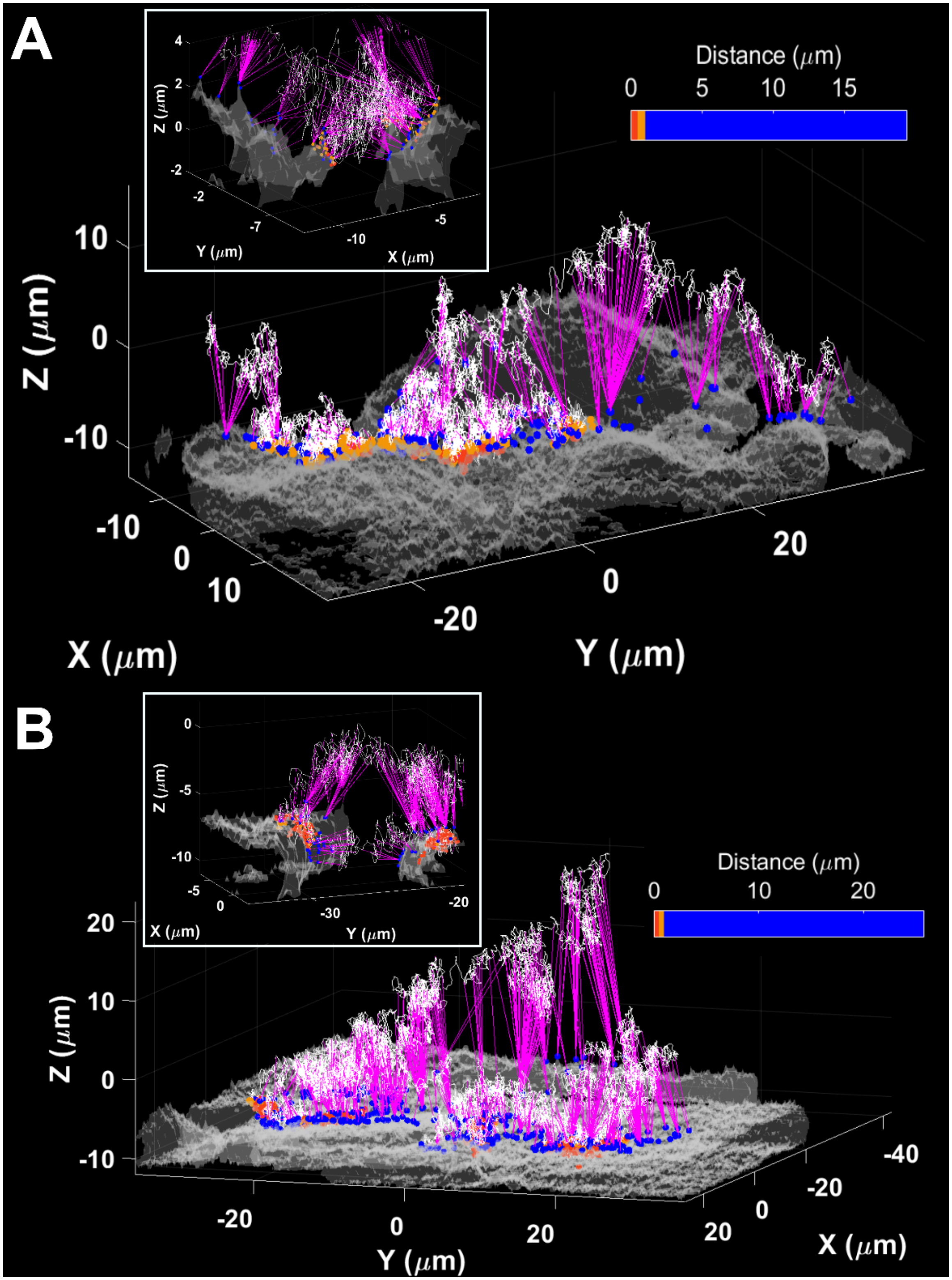
A
Distance (μm)
0
5
10
15
Z (μm)
X (μm)
Y (μm)
B
Distance (μm)
0
10
20
Z (μm)
Y (μm)
X (μm)

**Fig. S13. Virus-to-cell distance map calculation.**

(**A** and **B**) The full trajectory is shown in white. Using the tophat-generated isosurface (gray) from 3DFASTR imaging data the distance from each trajectory point to each surface vertex is calculated as describe in section 1.8.4. The minimum distance is then assigned to each data point (purple lines, only every $50^{th}$ data point is shown for clarity and every $10^{th}$ data point in the insert). The vectors from the vertex to the data point corresponding to the minimum distance are also shown as spheres on the cellular contour. (**A**) related to trajectory shown in Fig. 2. (**A**-**E**), while (B) is related to trajectory shown in Fig. 2. (**F**-**I**).

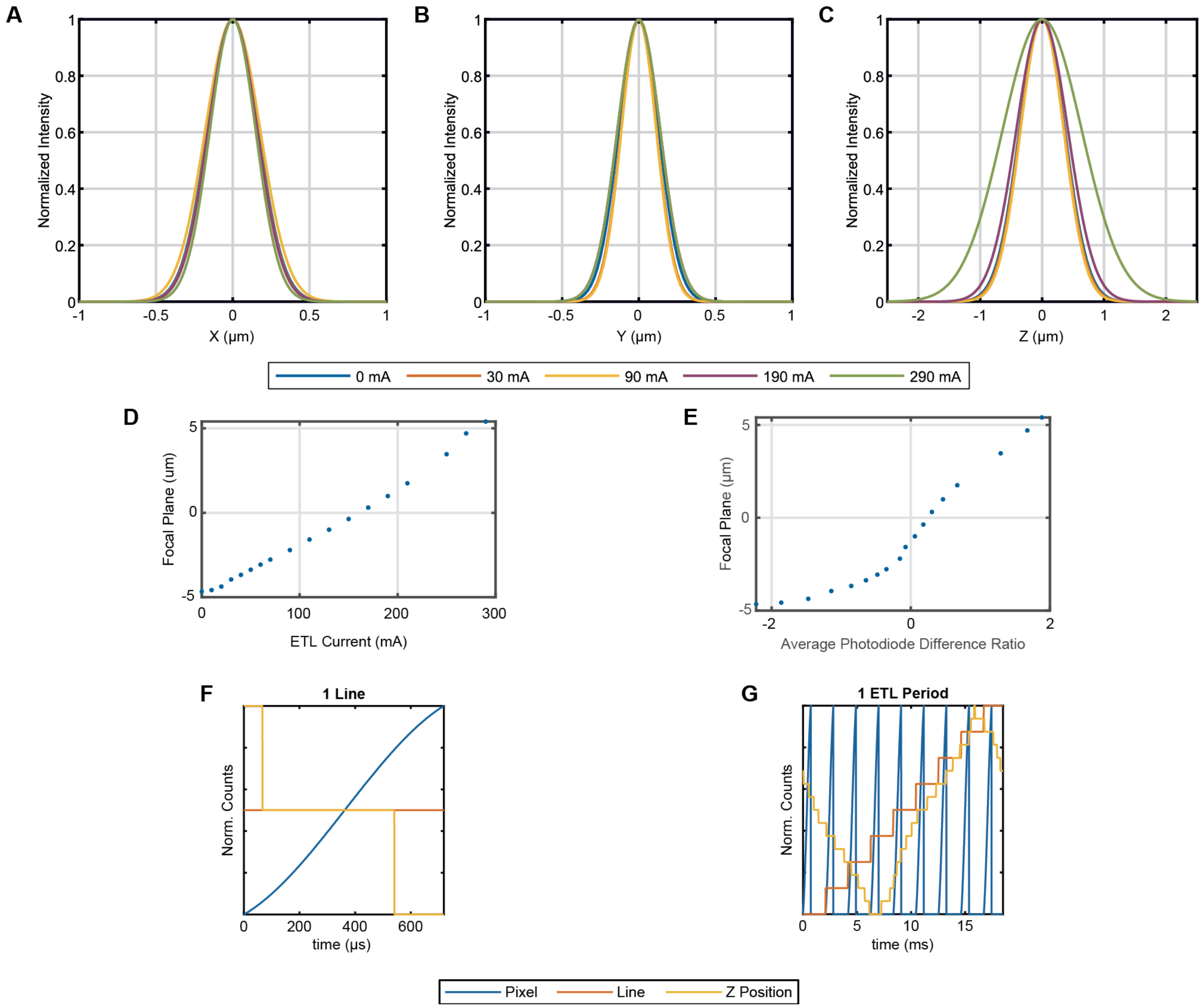

**Fig. S14. Imaging Point Spread Function and 3D-FASTR Improvements**

Imaging PSF from Gaussian fits shown at different ETL DC currents. (**A**) X-axis (**B**) Y-axis (**C**) Z-axis. The Z-axis shows the greatest change in FWHM at high currents. (**D**) Focal response curve of ETL shows nearly linear focal translation with current, the principle requirement to apply the 3D-FASTR theory. In the originally reported system, this curve was logarithmic, requiring modulation to drive an arbitrary waveform to coerce a linear response. (**E**) ETL Calibration Curve. (**F**) Relative scan progress between ETL and LSM pixel and line scans at 1 line (~625 µs) and (**G**) 1 ETL period (~20 ms)

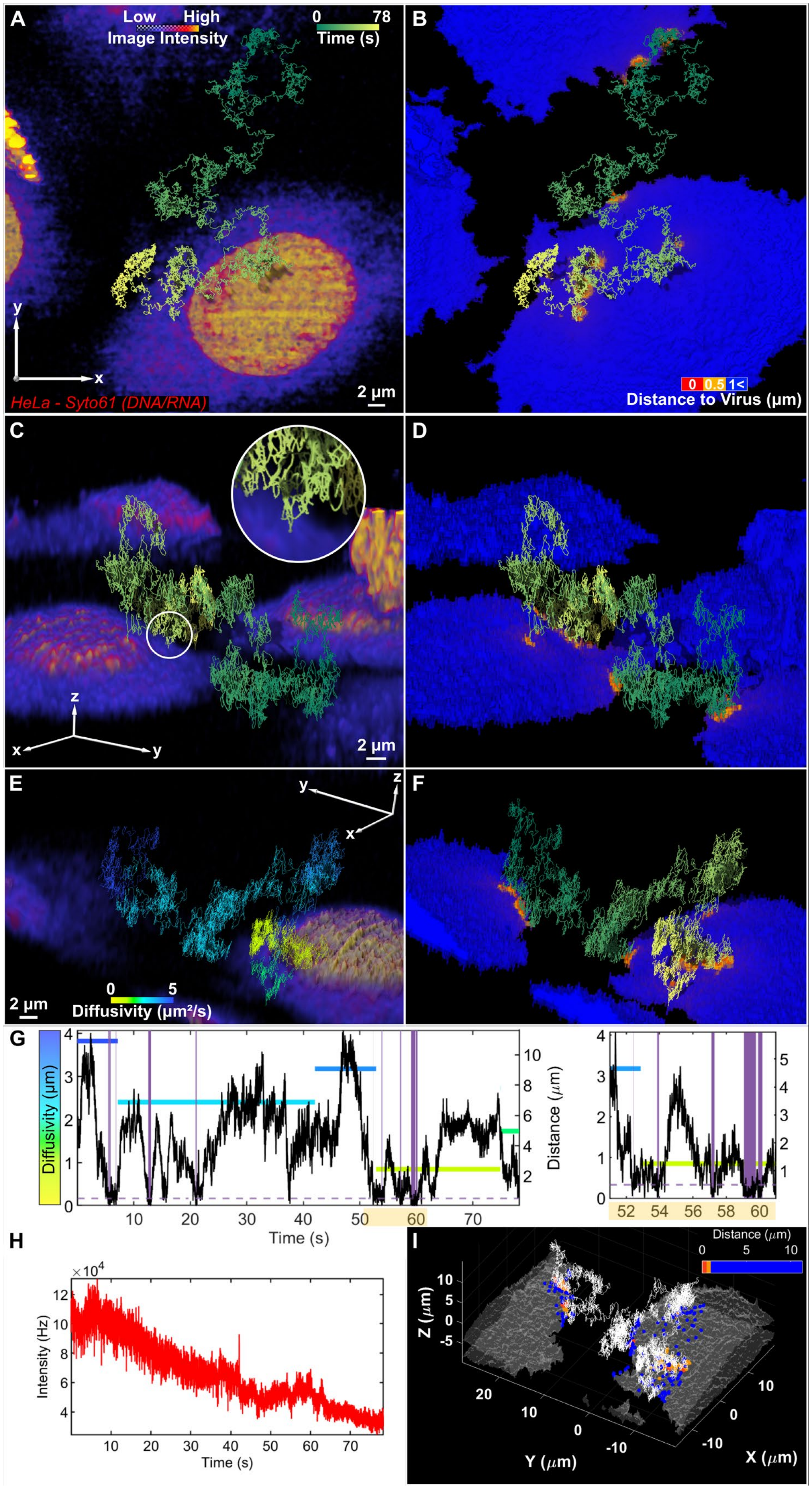

A
Low
High
Image Intensity
0
78
Time (s)
y
x
HeLa - Syto61 (DNA/RNA)
2 µm
B
0
0.5
1<
Distance to Virus (µm)
C
z
x
y
2 µm
D
E
y
z
x
2 µm
0
5
Diffusivity (µm²/s)
F
G
Diffusivity (µm)
Distance (µm)
Time (s)
H
×10^4
Intensity (Hz)
Time (s)
I
Distance (µm)
Z (µm)
Y (µm)
X (µm)

**Fig. S15. Additional example number 1 of VSV-G freely diffusing in the extracellular space.**

(**A** and **B**) Top-down view of virus making multiple close approaches on two adjacent live HeLa cells. (A and B) share the same axes and scale bar. (**C** and **D**) Same trajectory from a different angle. Circular insert exploded view of virus in close contact with the cell surface. (C and D) share the same axes and scale bar. (**E** and **F**) Cross-section with slice along y axis, overlaid with trajectory color-coded by diffusion coefficient in (E) and time in (F). (E and F) share the same axes and scale bar. (A, C and E) cells color-coded by intensity, while (B, D and F) display virus-to-cell distance map on surface of cell. (**G**) Left, correlation between diffusivity and distance from the cell surface of VSV-G VLP, close-approaches (below 0.5 μm) are color patched purple. Dashed purple line, 0.5 μm from cell surface. Right, enlarged view of data between 51-61 sec, matching trajectory segment displayed in circular insert of (C). (**H**) Tracking intensity trace. (**I**) Visual representation of virus-to-cell distance calculation. Isosurface volume render of cell image (gray) overlayed with trajectory sampled at 3 msec (white). Distance vectors are calculated from each trajectory timepoint to the isosurface (generated by top-hat transform) and displayed as color-coded spheres on the cell surface.

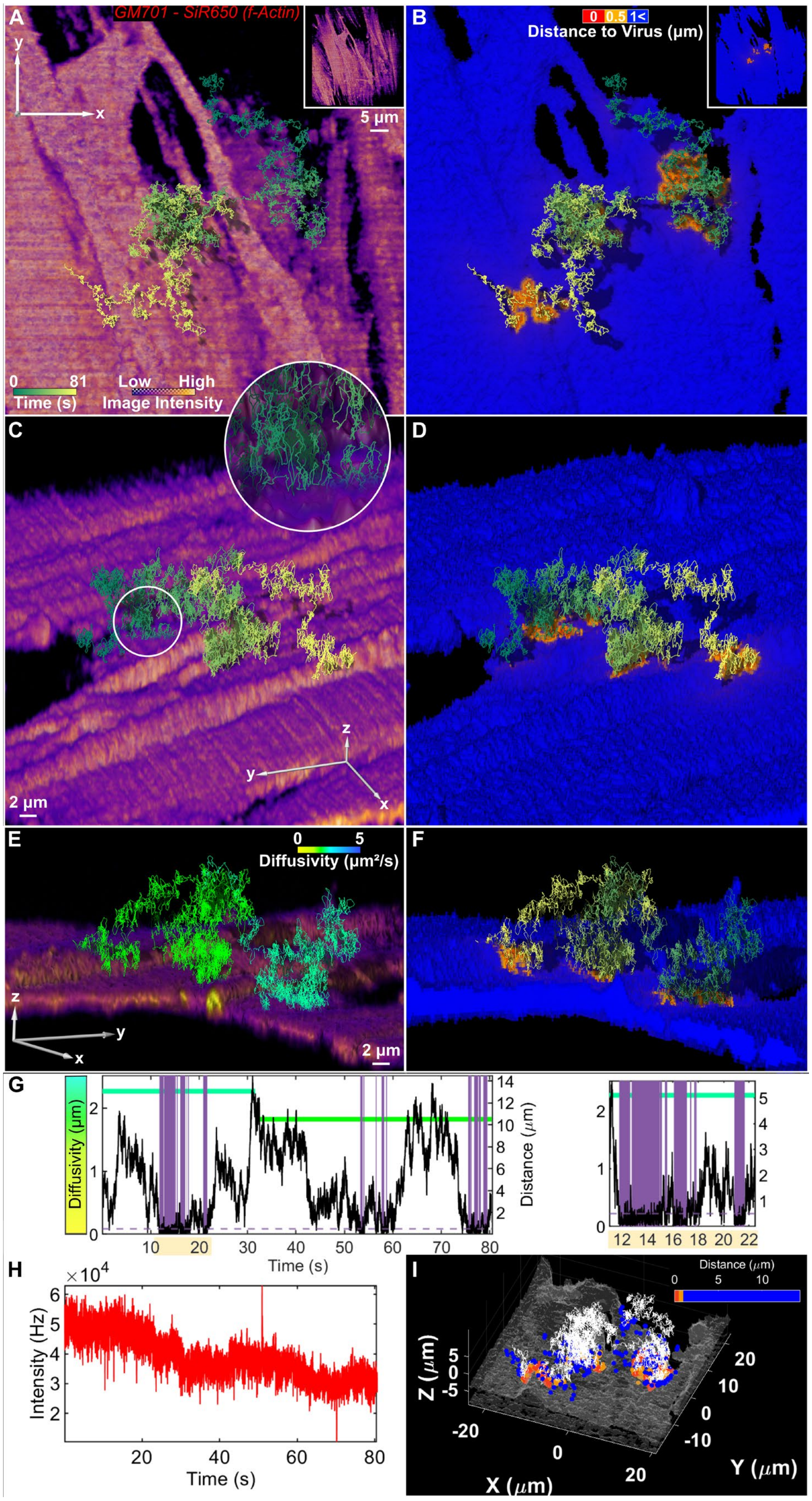
A
GM701 - SiR650 (f-Actin)
5 µm
0 81
Time (s)
Low High
Image Intensity
B
0 0.5 1<
Distance to Virus (µm)
C
2 µm
D
E
0 5
Diffusivity (µm²/s)
2 µm
F
G
Diffusivity (µm)
Distance (µm)
Time (s)
H
×10^4
Intensity (Hz)
Time (s)
I
Distance (µm)
Z (µm)
X (µm)
Y (µm)

**Fig. S16. Additional example number 2 of VSV-G freely diffusing in the extracellular space.**

(**A** and **B**) Top-down view of virus skimming across surface of live GM701 cell. (A and B) share the same axes and scale bar. (**C** and **D**) Same trajectory from a different angle. Circular insert, exploded view of virus in close contact with the cell surface. (C and D) share the same axes and scale bar. (**E** and **F**) Cross-section with slice along x and y axis, overlaid with trajectory color-coded by diffusion coefficient in (E) and time in (F). (E and F) share the same axes and scale bar. (A, C and E) cells color-coded by intensity, while (B, D and F) display virus-to-cell distance map on surface of cell. (**G**) Left, correlation between diffusivity and distance from the cell surface of VSV-G VLP, close-approaches (below 0.5 μm) are color patched purple. Dashed purple line, 0.5 μm from cell surface. Right, enlarged view of data between 11-22.5 sec matching trajectory segment displayed in circular insert of (C). (**H**) Tracking intensity trace. (**I**) Visual representation of virus-to-cell distance calculation. Isosurface volume render of cell image (gray) overlayed with trajectory sampled at 3 msec (white). Distance vectors are calculated from each trajectory timepoint to the isosurface (generated by top-hat transform) and displayed as color-coded spheres on the cell surface.

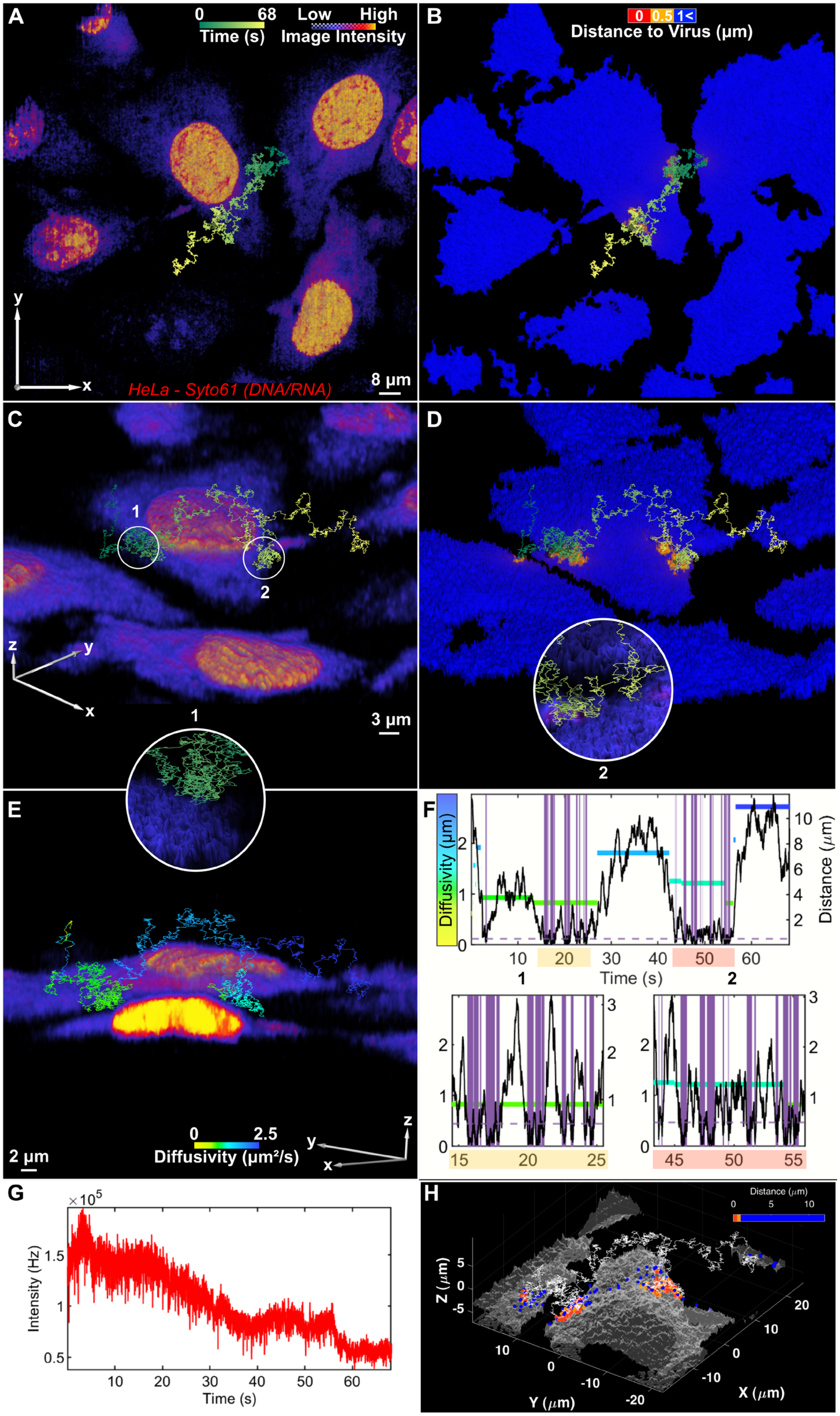
A
0 68
Time (s)
Low High
Image Intensity
HeLa - Syto61 (DNA/RNA)
8 µm
B
0 0.5 1<
Distance to Virus (µm)
C
1
2
3 µm
D
E
0 2.5
Diffusivity (µm²/s)
2 µm
F
Diffusivity (µm)
Distance (µm)
Time (s)
G
Intensity (Hz)
Time (s)
H
Distance (µm)
Z (µm)
Y (µm)
X (µm)

**Fig. S17. Additional example number 3 of VSV-G freely diffusing in the extracellular space.**

(**A** and **B**) Top-down view of virus trajectory pincering live HeLa cell. (A and B) share the same axes and scale bar. (**C** and **D**) Same trajectory from a different angle. Circular insert shows exploded views of virus in close contact with the cell surface. (C and D) share the same axes and scale bar. (**E**) Cross-section with slice along x and y axis, overlaid with trajectory color-coded by diffusion coefficient. (A, C and E) color-coded by image intensity, while (B, D) display virus-to-cell distance map on surface of cell. (**F**) Top, correlation between diffusivity and distance from the cell surface of VSV-G VLP, close-approaches (below 0.5 μm) are color patched purple. Dashed purple line, 0.5 μm from cell surface. Below left, enlarged view of data between 14.5-25.5 sec, associated with trajectory segment displayed in circular insert of (C). Below right, enlarged view of data between 43.25-56 sec, associated with trajectory segment displayed in circular insert of (D). (**G**) Tracking intensity trace. (**H**) Visual representation of virus-to-cell distance calculation. Isosurface volume render of cell image (gray) overlayed with trajectory sampled at 3 msec (white). Distance vectors are calculated from each trajectory timepoint to the isosurface (generated by top-hat transform) and displayed as color-coded spheres on the cell surface.

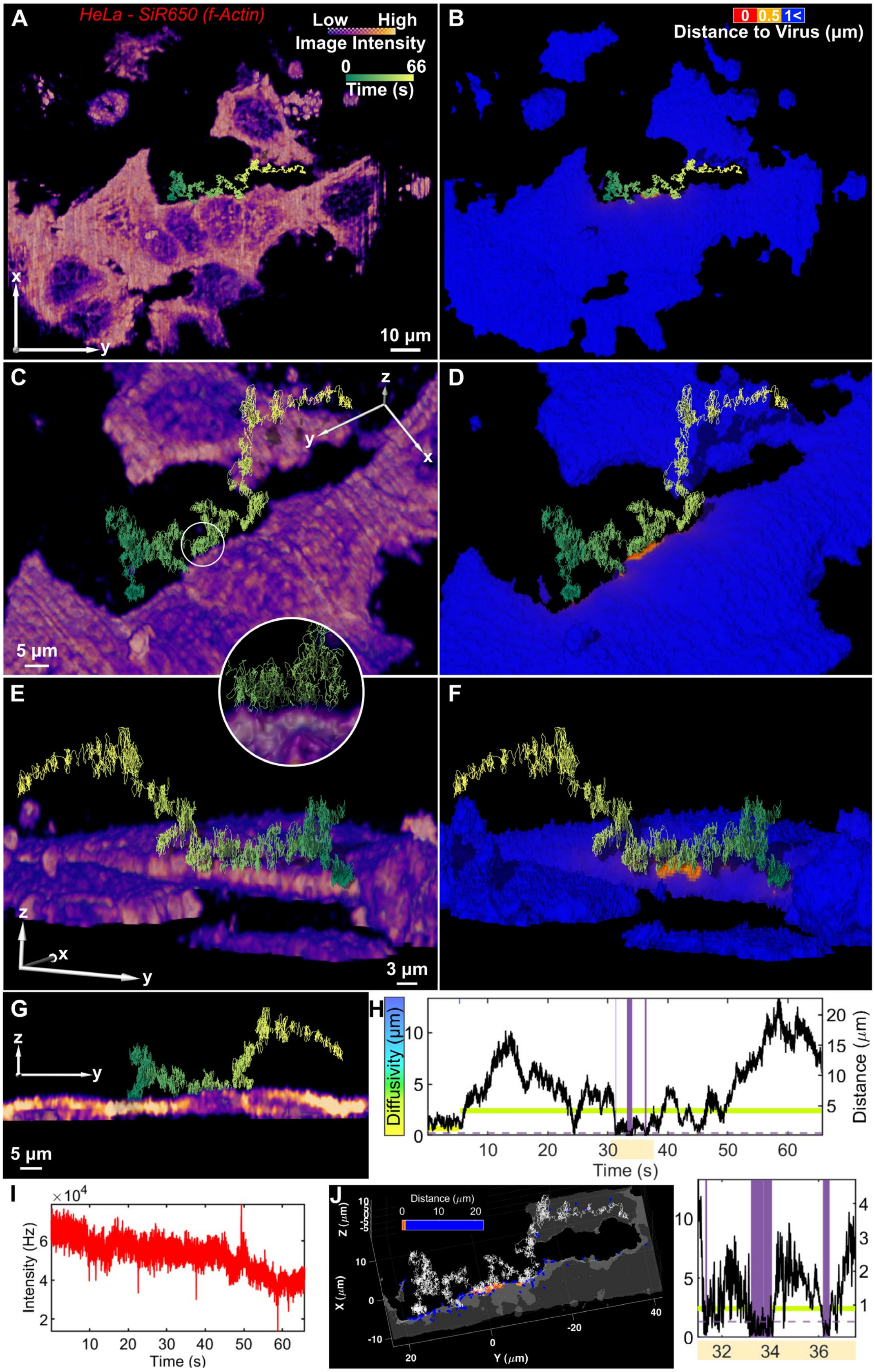

A
HeLa - SiR650 (f-Actin)
Low High
Image Intensity
0 66
Time (s)
10 μm
B
0 0.5 1<
Distance to Virus (μm)
C
5 μm
D
E
3 μm
F
G
5 μm
H
Diffusivity (μm)
Distance (μm)
Time (s)
I
Intensity (Hz)
Time (s)
J
Distance (μm)
Y (μm)
X (μm)
Z (μm)

**Fig. S18. Additional example number 4 of VSV-G freely diffusing in the extracellular space.**

(**A** and **B**) Top-down view of virus hopping along the edge of a live HeLa cell. (A and B) share the same axes and scale bar. (**C** and **D**) Same trajectory from a different angle. Circular insert shows exploded view of virus in close contact with the cell surface. (C and D) share the same axes and scale bar. (**E** and **F**) Same trajectory from reverse angle. (G) Lateral view with slice along y axis. (A, C, E, and G) color-coded by cell intensity, while (B, D, and F) display virus-to-cell distance map on surface of cell. (**H**) Top, correlation between diffusivity and distance from the cell surface of VSV-G VLP, close-approaches (below 0.5 μm) are color patched purple. Dashed purple line, 0.5 μm from cell surface. Bottom, enlarged view of data between 31-37.5 sec, associated with trajectory segment displayed in circular insert of (C). (**I**) Tracking intensity trace. (**J**) Visual representation of virus-to-cell distance calculation. Isosurface volume render of cell image (gray) overlayed with trajectory sampled at 3 msec (white). Distance vectors are calculated from each trajectory timepoint to the isosurface (generated by top-hat transform) and displayed as color-coded spheres on the cell surface.

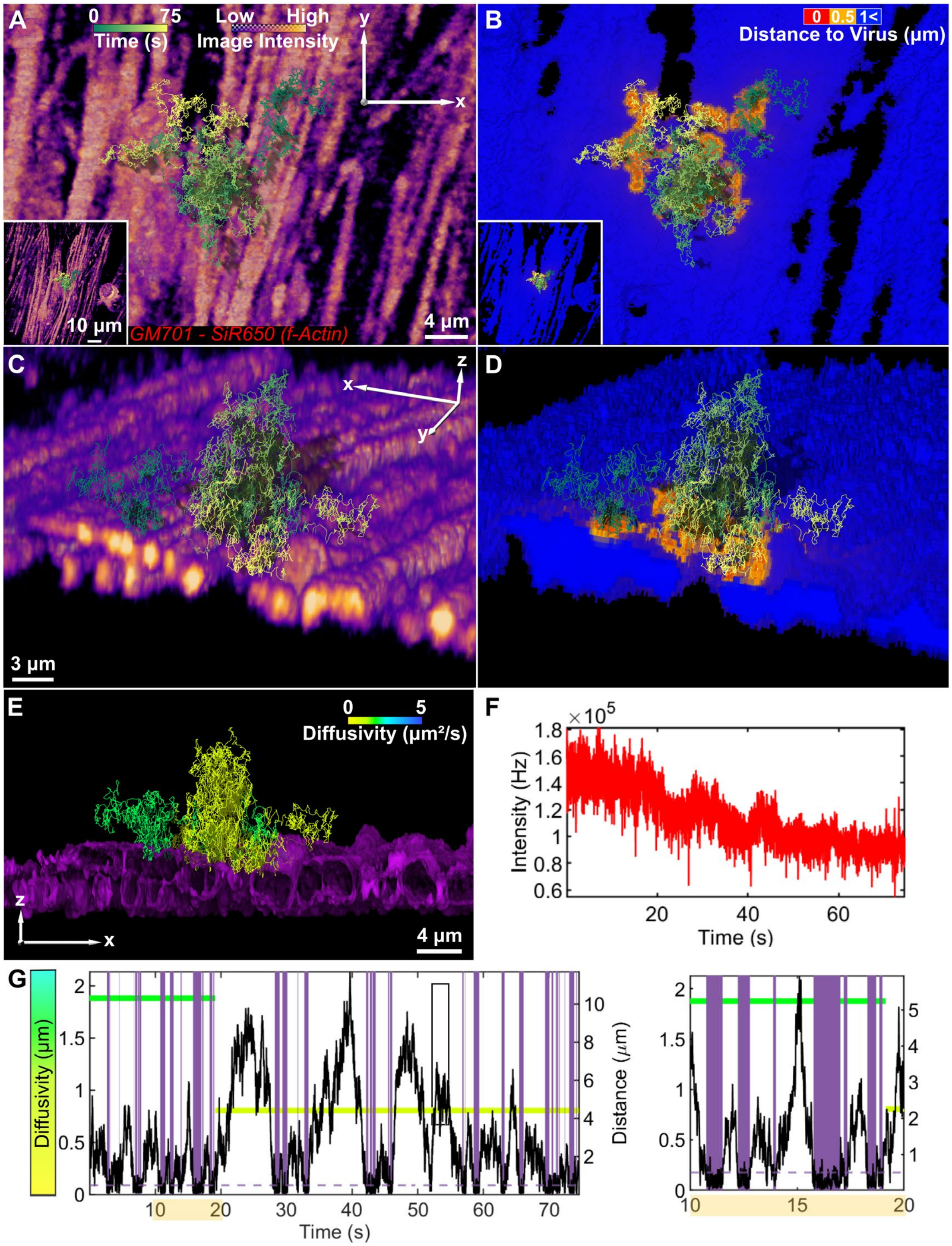

**Fig. S19. Additional example number 5 of VSV-G freely diffusing in the extracellular space.** (**A** and **B**) Top-down view of virus freely diffusing close to the surface of a live GM701 fibroblast cell for long periods without landing. (A and B) share the same axes and scale bar. (**C** and **D**) Same trajectory from a different angle with slice along x axis. (C and D) share the same axes and scale bar. (A and C) color-coded by cell intensity, while (B and D) display virus-to-cell distance map on surface of cell. (**E**) Lateral view with slice along x axis, with trajectory color-coded by diffusivity determined by change-point analysis. Three-dimensional reconstruction with isosurface representation of cell surface to highlight cell surface undulations and virus particle diffusion within surface crevasses (**F**) Tracking intensity trace. (**G**) Left, correlation between diffusivity and distance from the cell surface of VSV-G VLP, close-approaches (below 0.5 μm) are color patched purple. Dashed purple line, 0.5 μm from cell surface. Right, enlarged view of data between 10-20 sec, highlighting multiple close-approaches of the virus with the cell surface.

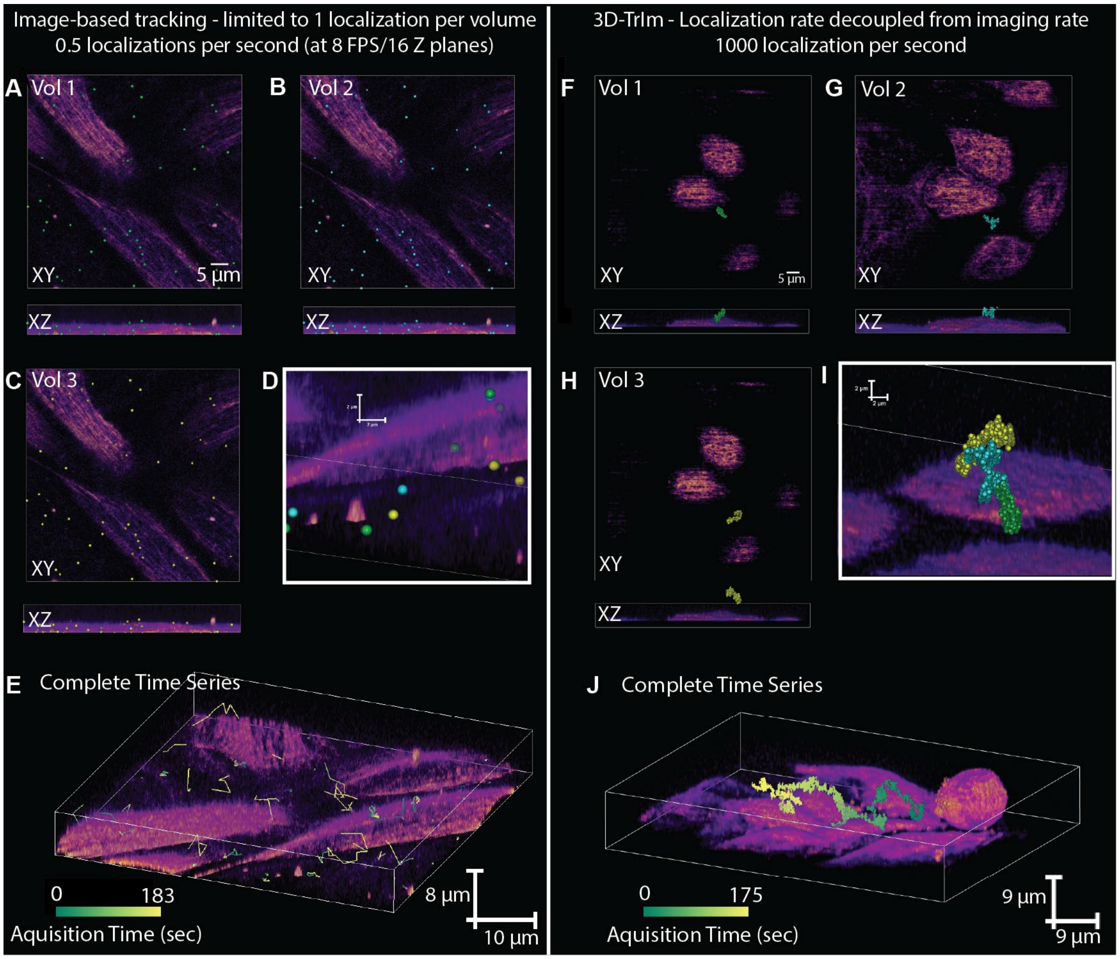

**Fig. S20. Comparison of image-based tracking and imaging (spinning disk) with active-feedback tracking and complementary 3D imaging (3D-TrIm).**

Fluorescently-labeled virus-like particles (eGFP.Vpr VSVG) were introduced to cultures of live HeLa cells stained with SiR650-actin (100 nM). (A-E) Data collected on Andor DragonFly spinning disk confocal microscope. The same area was sampled continuously. Each volume has a depth of 8 µm split into 16 z planes. Virus particles and cells were imaged simultaneously with a camera exposure time of 40 msec. (F-J) Data collected on 3D-TrIm microscope. A single virus particle was tracked continuously (1 msec sampling shown) and the surrounding area was imaged 5 µm below the particle and 3 µm above, to give an approximate 8 µm volume. The resulting trajectories over the entire acquisition period are shown and color-coded by time. Experiments were performed in live-cell image solution with 2% FBS and maintained at 37oC.

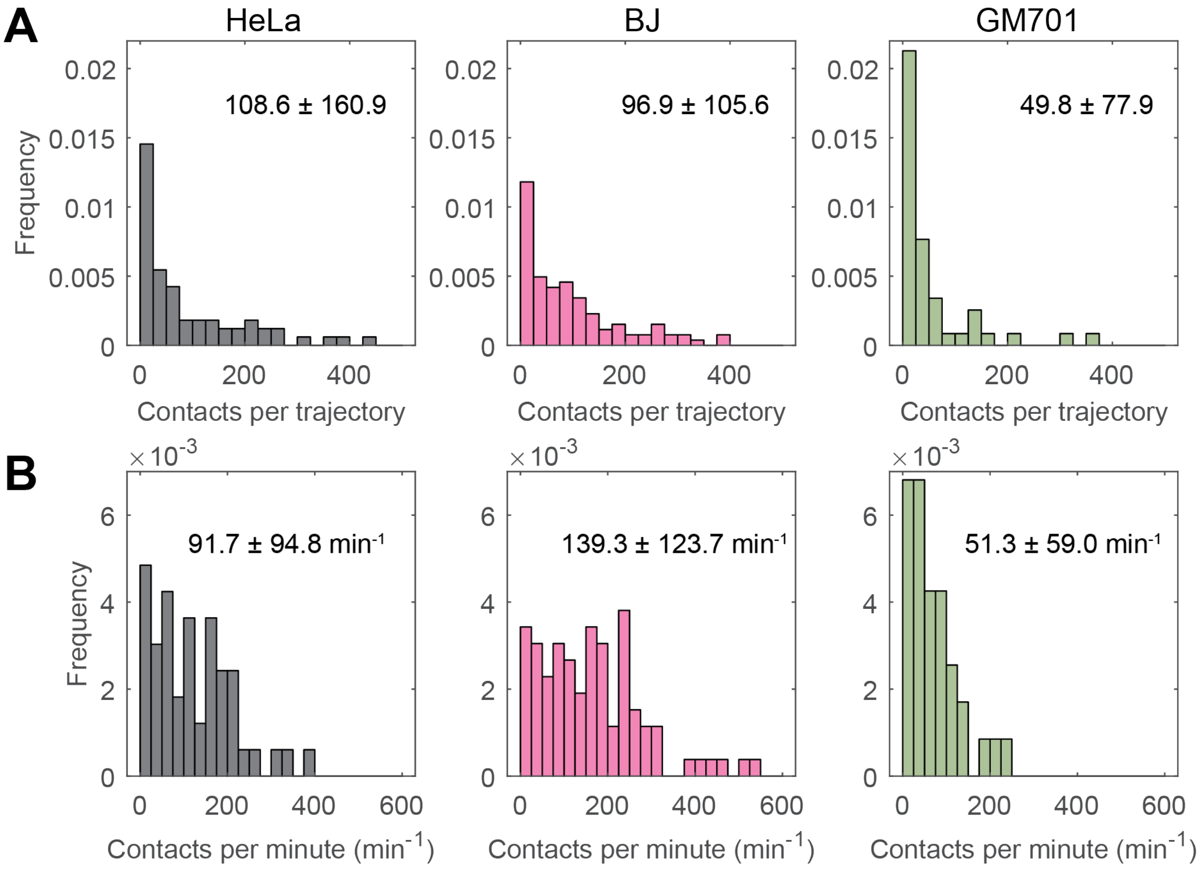

**Fig. S21. Number of transient contact events, related to Fig. 2.**

Number of transient contact events (**A**) per trajectory (**B**) per minute, for HeLa (black, n = 66 trajectories), BJ fibroblasts (magenta, n = 105 trajectories), and GM701 fibroblasts (green, n = 47 trajectories). Mean ± standard deviation. As our methodology gives us the capability to continuously follow VLPs from one event to next in their entirety we were able to calculate accumulative contact time of each trajectory with the cell surface (HeLa: 4.6 ± 6.4 sec, BJ: 4.2 ± 4.9 sec, GM701: 1.9 ± 3.4 sec (mean ± SD)). When comparing the percentage of time in contact with the cell surface our data suggests VSV-G VLPs have a higher affinity for LDLR expressing cells (HeLa: 6.6 ± 7.2 %, BJ: 10.0 ± 8.9 %, GM701: 3.2 ± 3.8 % (mean ± SD)).

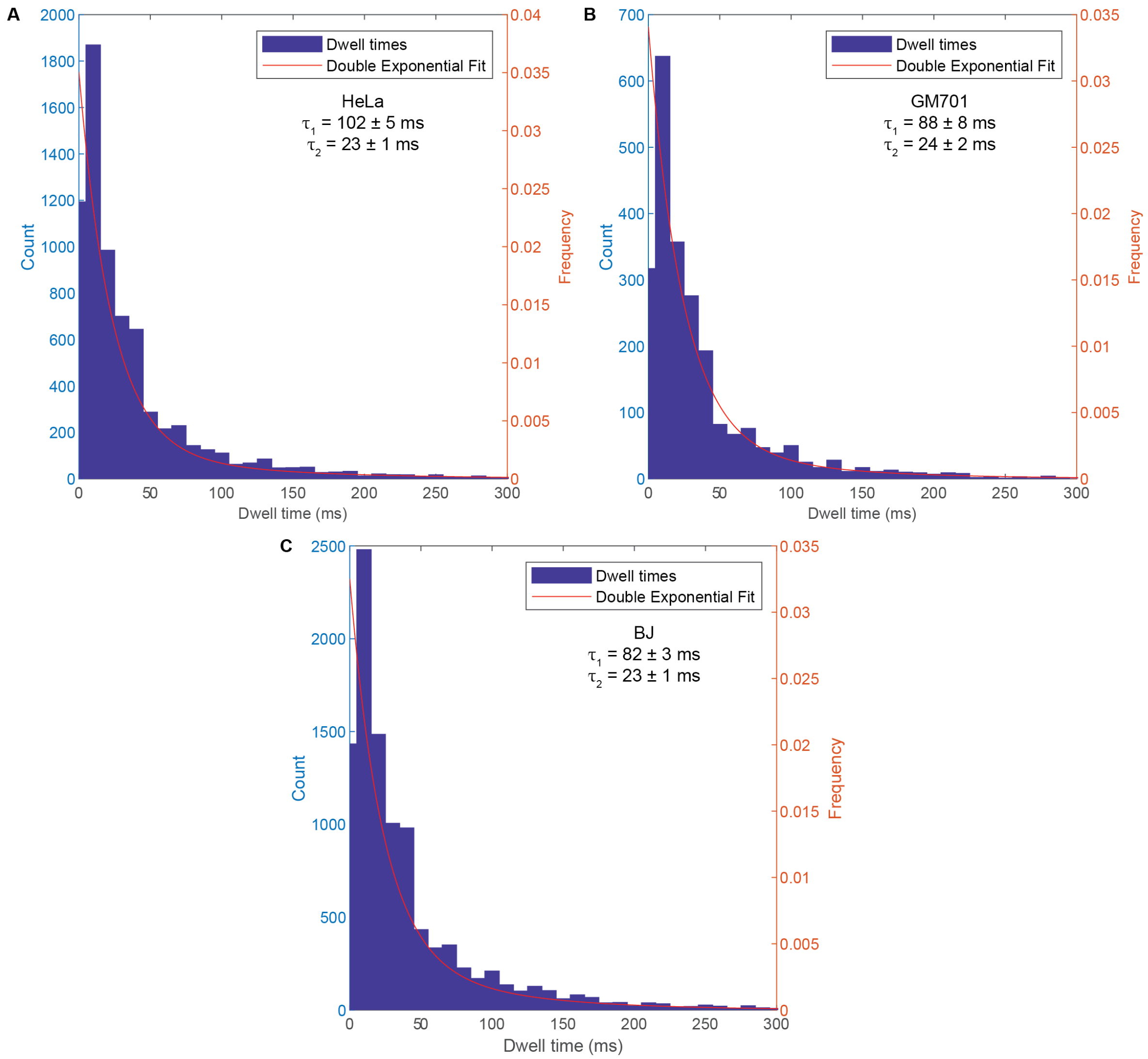

**Fig. S22. Distribution of VLP dwell times near the cell surface, related to Fig.2.**

The dwell times of VLP near the surface of live cells were best fit by a double exponential distribution. (**A**) VLPs near the surface of HeLa cells exhibited short (23 ± 1 msec) and long (102 ± 5 msec) dwell times. (**B, C**) Receptor-negative (GM701, B) and receptor-positive (BJ, C) cells yield the same fast component as HeLa but with the significantly shorter slow component, suggesting a role of cell morphology in these longer dwell time events.

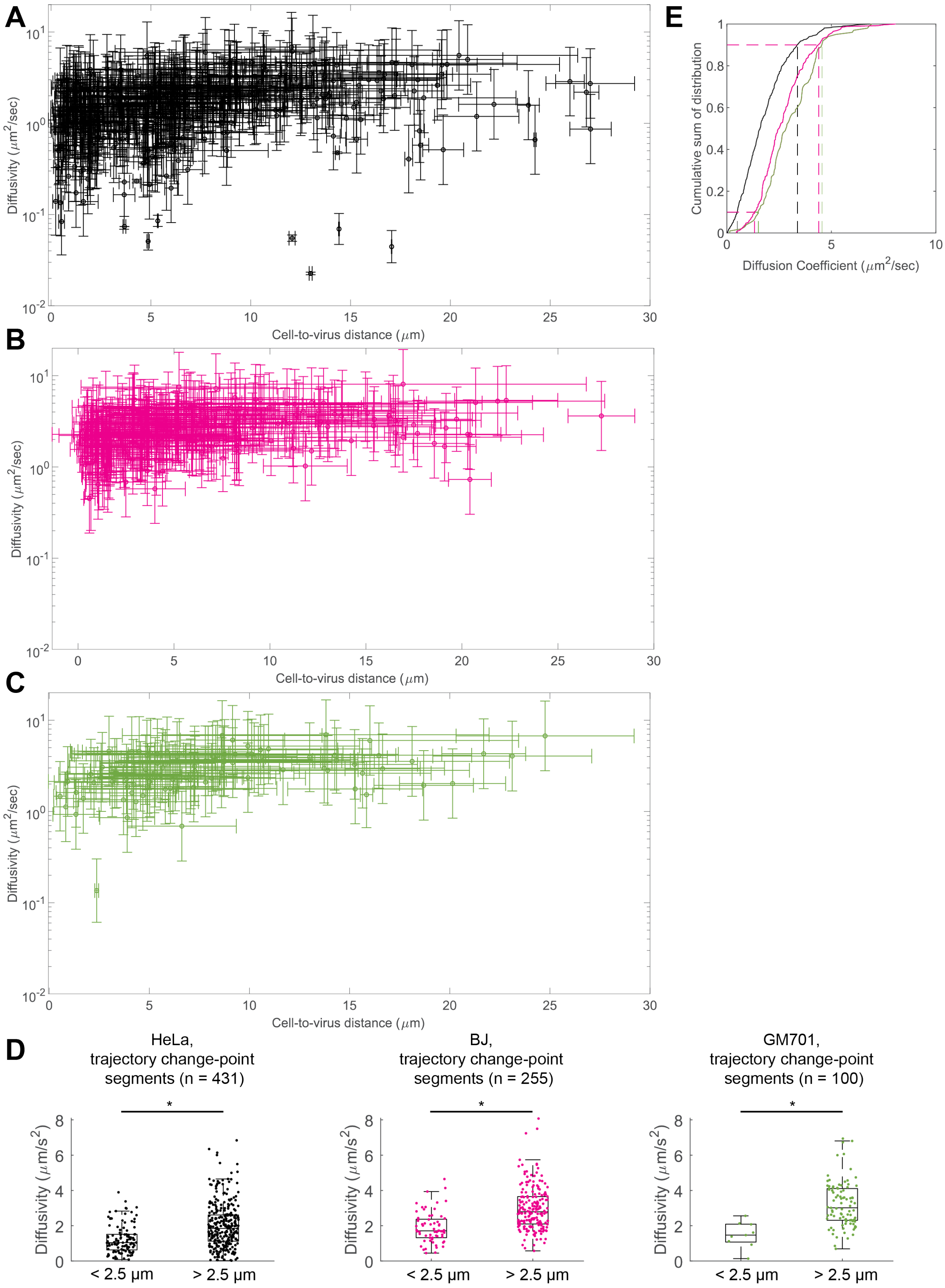
A
B
C
D
E
Diffusivity ($\mu m^2$/sec)
Cell-to-virus distance ($\mu m$)
Cumulative sum of distribution
Diffusion Coefficient ($\mu m^2$/sec)
HeLa,
trajectory change-point
segments (n = 431)
BJ,
trajectory change-point
segments (n = 255)
GM701,
trajectory change-point
segments (n = 100)
Diffusivity ($\mu m/s^2$)
< 2.5 µm
> 2.5 µm
*

**Fig. S23. Dissecting the relationship between VSV-G VLP diffusivity behavior and distance from the cell surface, related to Fig. 2.**

The diffusivity of trajectory segments derived from change-point analysis is spatially registered with respect to the cell surface (section 1.7: Diffusion coefficient and distance measurement calculations). (**A**) HeLa cells, stained with SiR650-actin (n = 66 trajectories and n = 413 segments). (**B**) BJ fibroblast cells, stained with SiR650-actin (n = 105 trajectories and n = 255 segments). (**C**). GM701 fibroblast cells, stained with SiR650-actin (n = 47 trajectories and n = 100 segments). Error bars representing the standard deviation in the diffusivity and distance measurements. (**D**) Distributions of VSV-G VLP diffusivity within 2.5 µm of the cell surface or above 2.5 µm. HeLa (below 2.5 µm = 1.24 ± 0.82 $\mu m^2/s$, n = 107 segments and above 2.5 µm = 2.00 ± 1.25 $\mu m^2/s$, n = 306 segments). BJ (below 2.5 µm = 1.91 ± 0.88 $\mu m^2/s$, n = 65 segments and above 2.5 µm = 2.98 ± 1.23 $\mu m^2/s$, n = 190 segments). GM701 (below 2.5 µm = 1.49 ± 0.73 $\mu m^2/s$, n = 9 segments and above 2.5 µm = 3.22 ± 1.28 $\mu m^2/s$, n = 91 segments). Statistical significance was assessed by Kruskal-Wallis test on the p-value at $p < 0.01$. Standard box plots are provided with the center representing the median, and the top and bottom edges showing the third and first quartiles of the data. The whiskers encompass data points not considered outliers. (**E**) Cumulative sum distributions of diffusion coefficients for VSV-G VLP trajectory segments on HeLa (black), BJ (magenta), and GM701 (green). Dashed lines intersect at $10^{th}$ and $90^{th}$ percentiles, respectively.

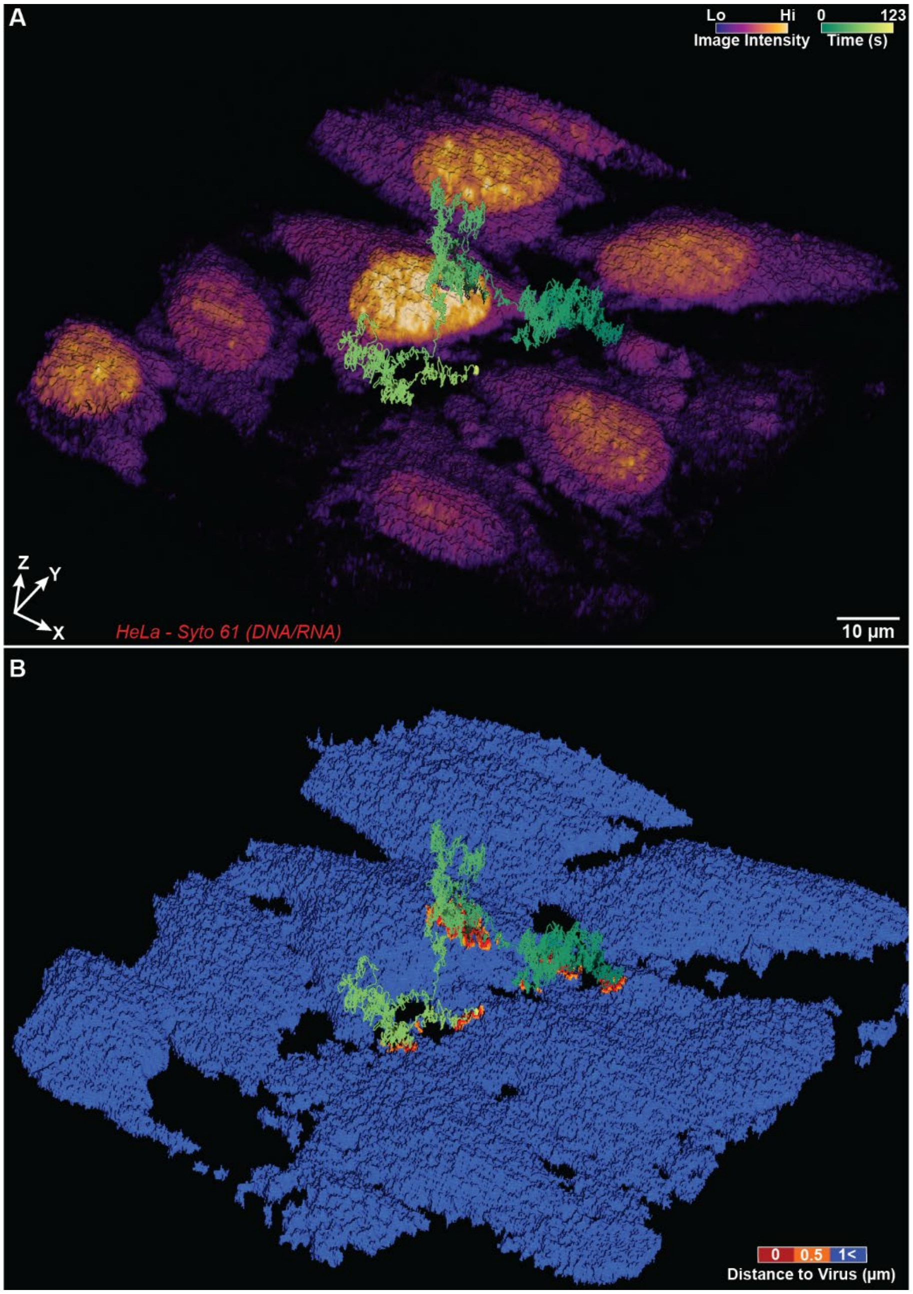

**Fig. S24. Global Volume Renders Related to Fig. 3A-D**

Global volume space render of Viral Binding Trajectory 1. (**A**) Intensity render of 3D-TrIm trajectory data. (**B**) Corresponding distance map showing virus-to-cell distances.

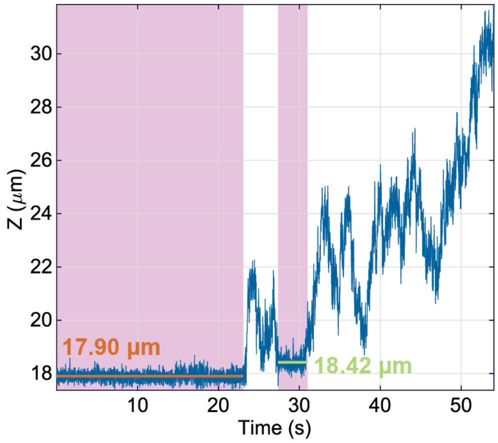

**Fig. S25. Axial Trace of Viral Trajectory Related to Fig. 3E-G**

Axial trace of viral trajectory shows two binding events, the first of which occurs 520 nm lower than the second, agreeing with the image shown in Fig. 3G which depicts the trajectory embedded inside the groove between two actin fiber bundles.

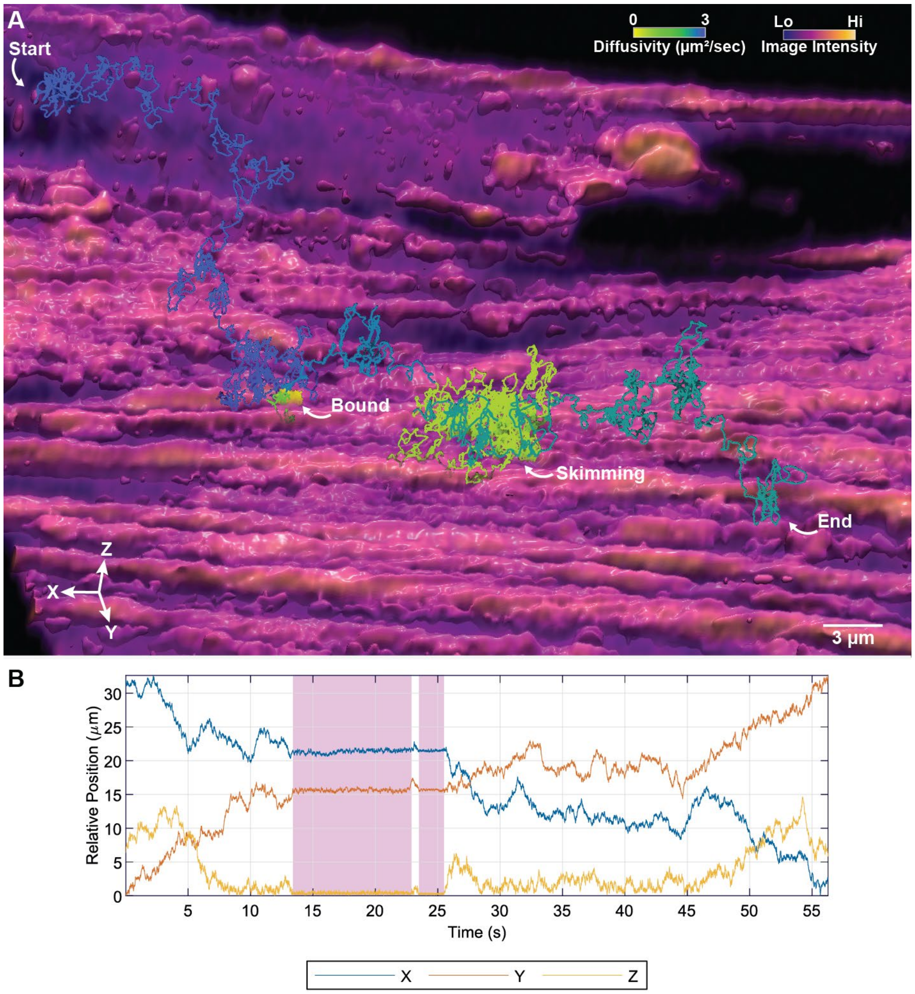

**Fig. S26. Additional Example Number 1 of VSV-G Binding Event**

**(A)** 3D-TrIm render of BJ Fibroblast cells stained with SiR-650 (f-Actin). (**B**) Distance-diffusivity trace of viral trajectory. This trajectory begins with rapid extracellular diffusion before binding to the surface. A small ~1 second detachment occurs before re-binding for several seconds and detaching again where skimming is observed with inhibited diffusivity compared to the initial free diffusion, followed by free diffusion that has greater diffusivity than observed while skimming, but lower than the initial free diffusion.

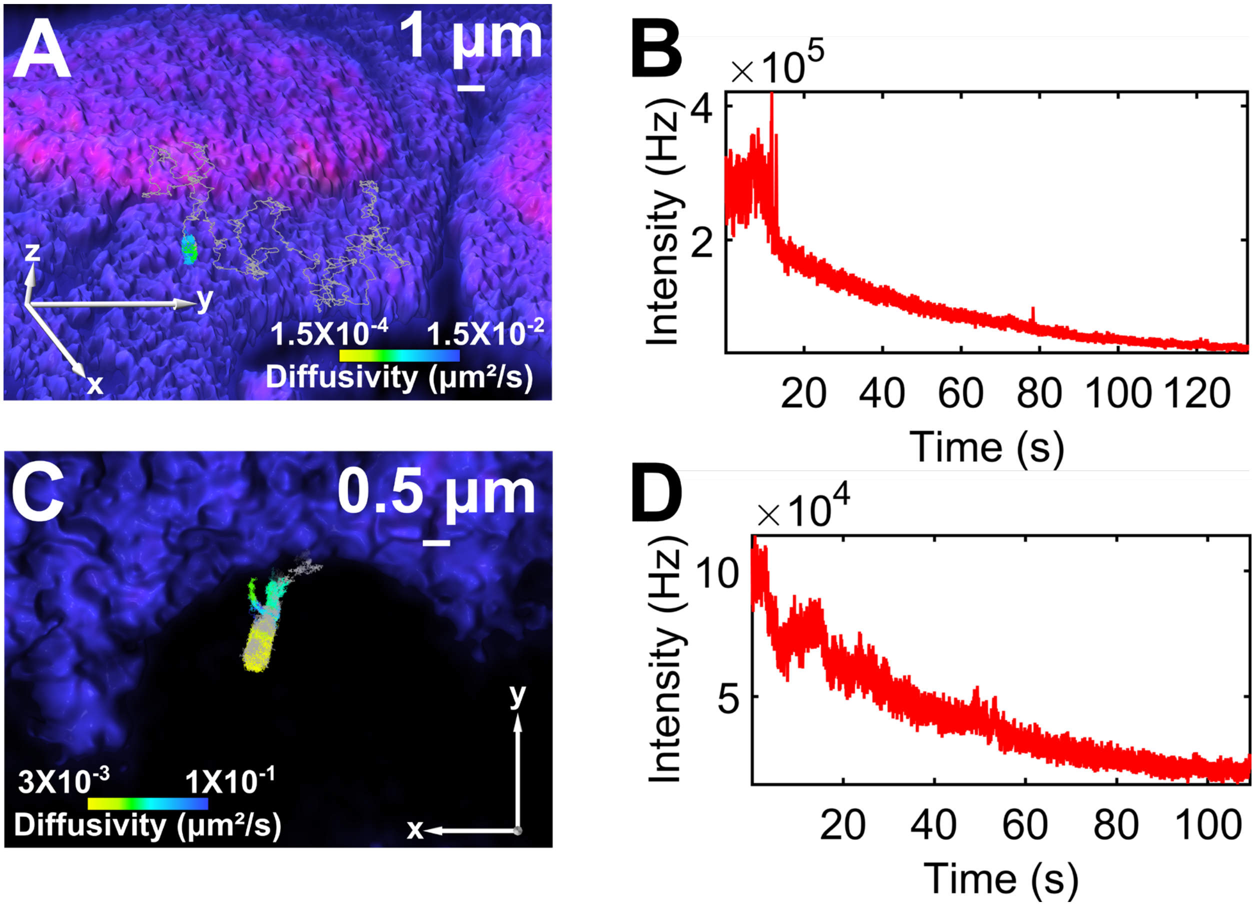

**Fig. S27. High resolution tracking and imaging data, related to Fig. 4.**

(**A**) Magnified view of trajectory shown in Fig. 4A-E. Free diffusion and membrane-bound behavior can be detected using change-point analysis to separate the trajectory into distinct diffusive states, free portion greater than $1.5\times10^{-2}$ $\mu m^2/s$ colored gray. (**B**) Tracking intensity trace. (**C**) Magnified view of trajectory shown in Fig. 4. (F-I) with the trajectory separated into distinct diffusive states via change-point analysis. Segments of the trajectory not used in the protrusion cylinder fit colored gray. (**D**) Tracking intensity trace.

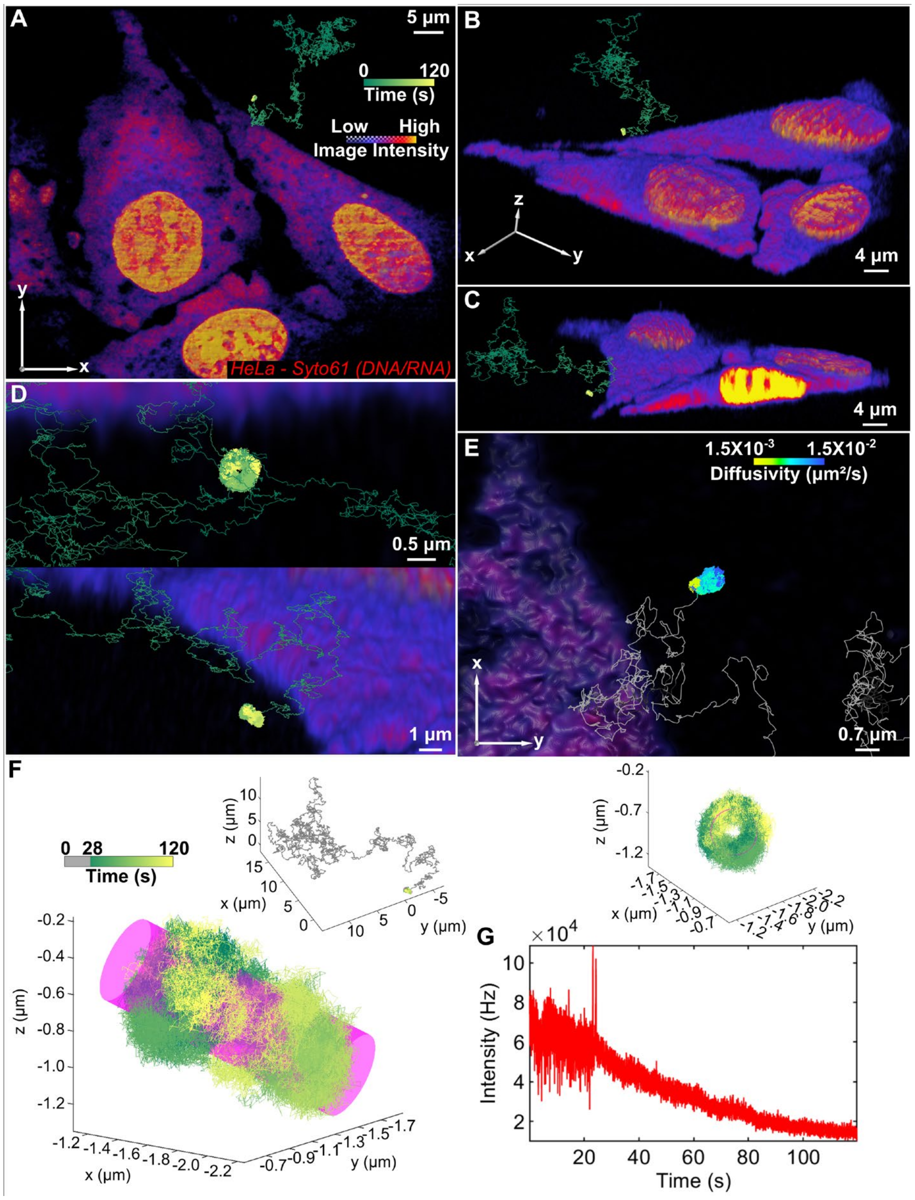

**Fig. S28. Additional example number 1 of virus interaction with protrusion.** (**A**) Top-down view of VSV-G landing on filopodium protruding from the surface of a live cell. 3D volume rendering of live HeLa cells (stained with SYTO61) co-registered with high resolution virus trajectory from 4D tracking and imaging data (**B** and **C**) Same trajectory from different orientation. (**D**) Magnified views showing cylindrical nature of trajectory and proximity to cell surface. (**E**) Top-down view, with trajectory colored by diffusion coefficient, free portion greater than $1.5\times10^{-2}$ $\mu m^2/s$ colored gray. (**F**) Cylindrical fitting of the bound portion of the trajectory in (A-D) (28-120 sec), fitted cylinder (magenta) has a radius of 122 ± 8.4 nm after taking into consideration the tracked particle diameter. (**G**) Tracking particle intensity trace.

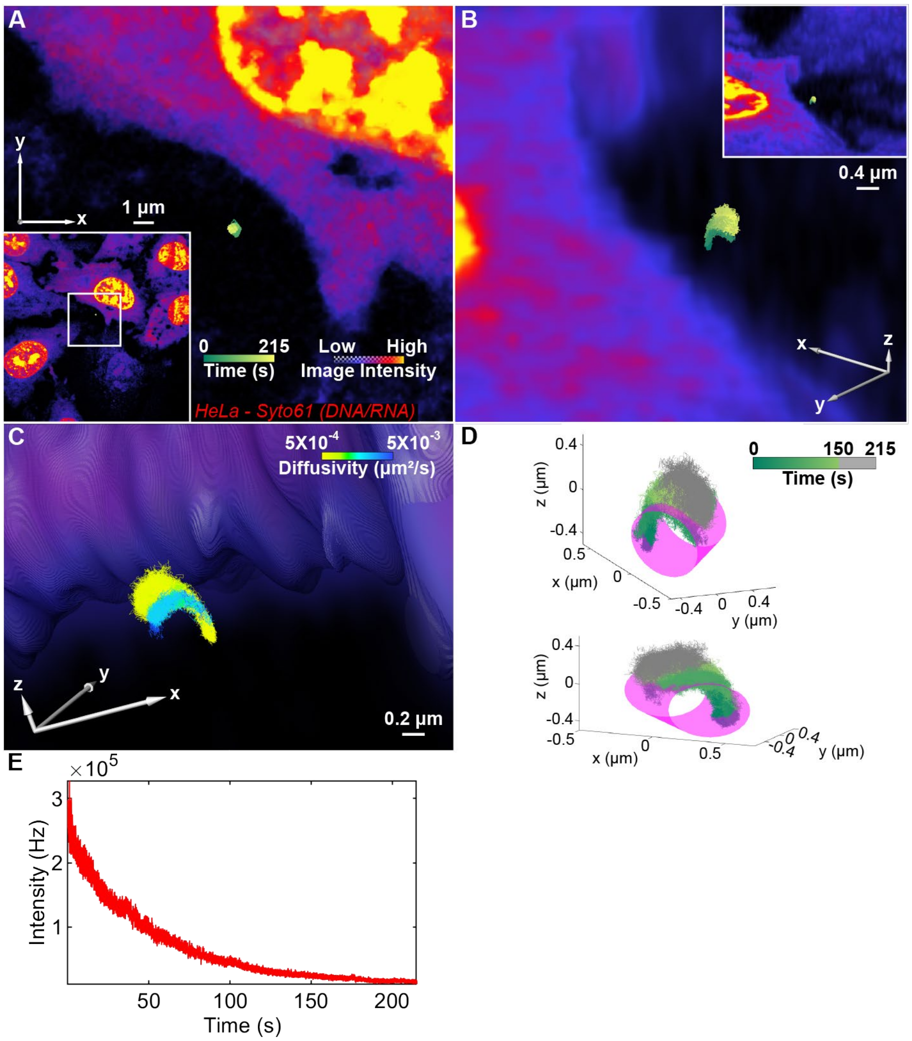

**Fig. S29. Additional example number 2 of virus interaction with protrusion.**

(**A**) Top-down view of VSV-G traveling along filopodium. 3D volume rendering of live HeLa cells (stained with SYTO61) co-registered with high resolution virus trajectory from 4D tracking and imaging data (**B** and **C**) Same trajectory from different orientation, with trajectory colored by time (**B**) and diffusion coefficient (**C**). (**D**) Cylindrical fitting of the bound portion of the trajectory in (**A**-**C**) (1-150 sec), fitted cylinder (magenta) has a radius of 168 ± 8.4 nm after taking into consideration the tracked particle diameter. (**E**) Tracking particle intensity trace.

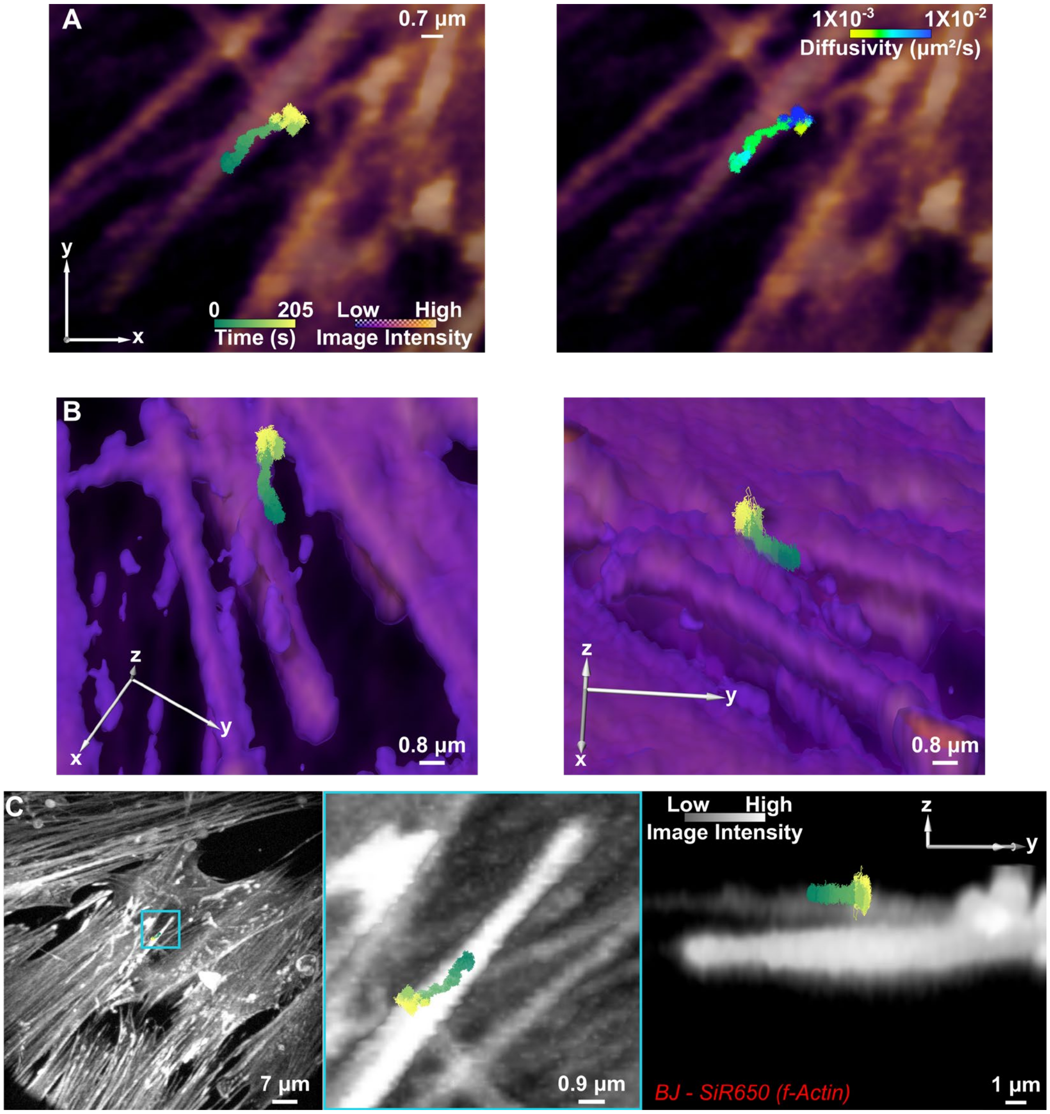

**Fig. S30. High resolution tracking and simultaneous 3D imaging localizes VSV-G VLP diffusion along fibroblast protrusion, additional example number 1.**

(**A**) Left, top-down view of VSV-G VLP 'surfing' on actin-stained protrusion of BJ fibroblast. Trajectory colored by time. The lower, brighter, z-planes have been omitted to help visualize the protrusion the VLP is localized to. Right, same view but trajectory color-coded by diffusivity. (**B**) Snapshots of 3D reconstruction. The sustained linear motion correlates with the lateral surface of the protrusion, large changes in diffusivity where detected. (**C**) (left) XY MIP, (middle) magnified, and (right) YZ MIP.

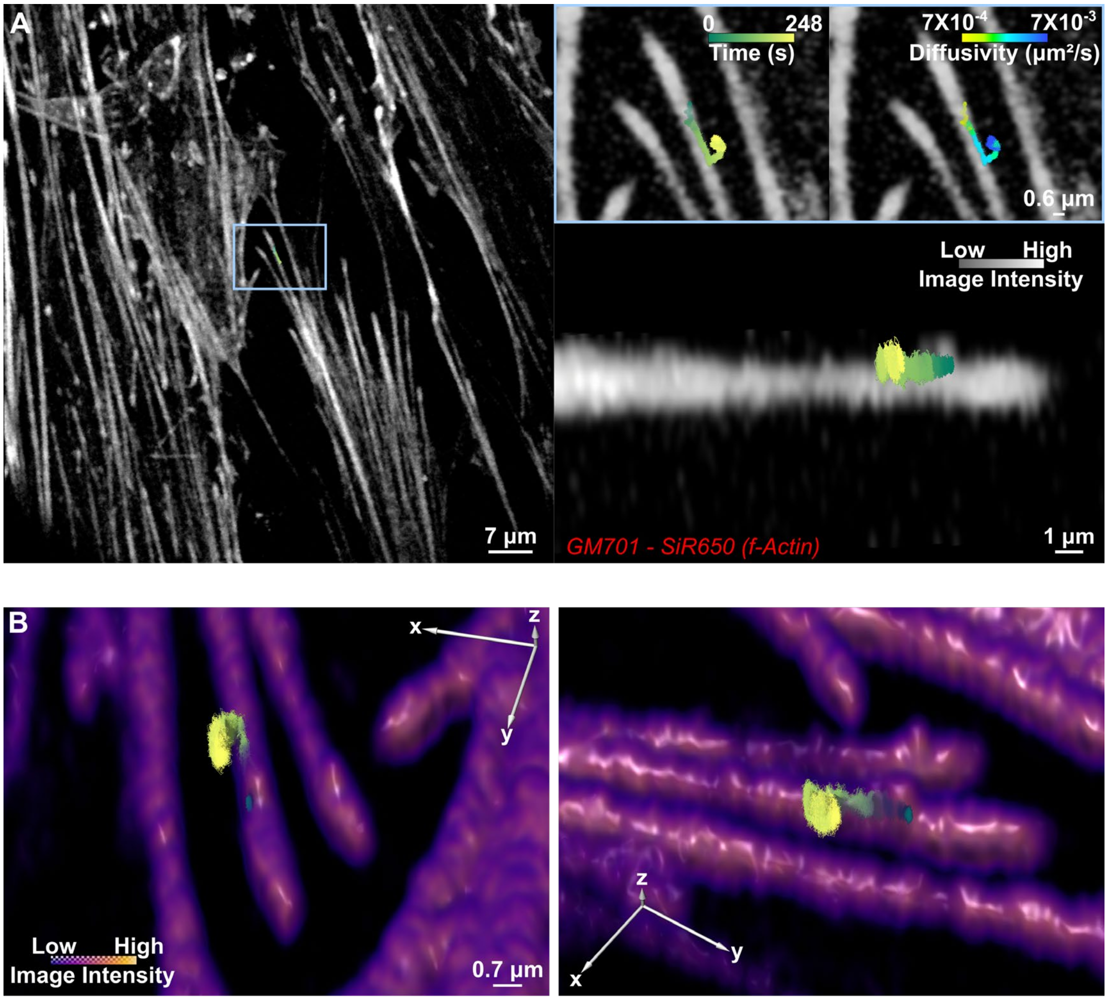

**Fig. S31. High resolution tracking and simultaneous 3D imaging localizes VSV-G VLP diffusion along fibroblast protrusion, additional example number 2.**

(**A**) Top-down MIP, of VSV-G VLP 'surfing' on actin-stained protrusion of LDLR deficient GM701 cell. Right top, magnified views with trajectory colored by time, or diffusion coefficient, respectively. Right, bottom lateral view. (**B**) 3D reconstruction, snapshots from different orientations. The sustained linear motion correlates with the lateral surface of the protrusion as the VLP moves towards the body of the cell before diverging and undergoing more random but confined diffusion on unstained material.

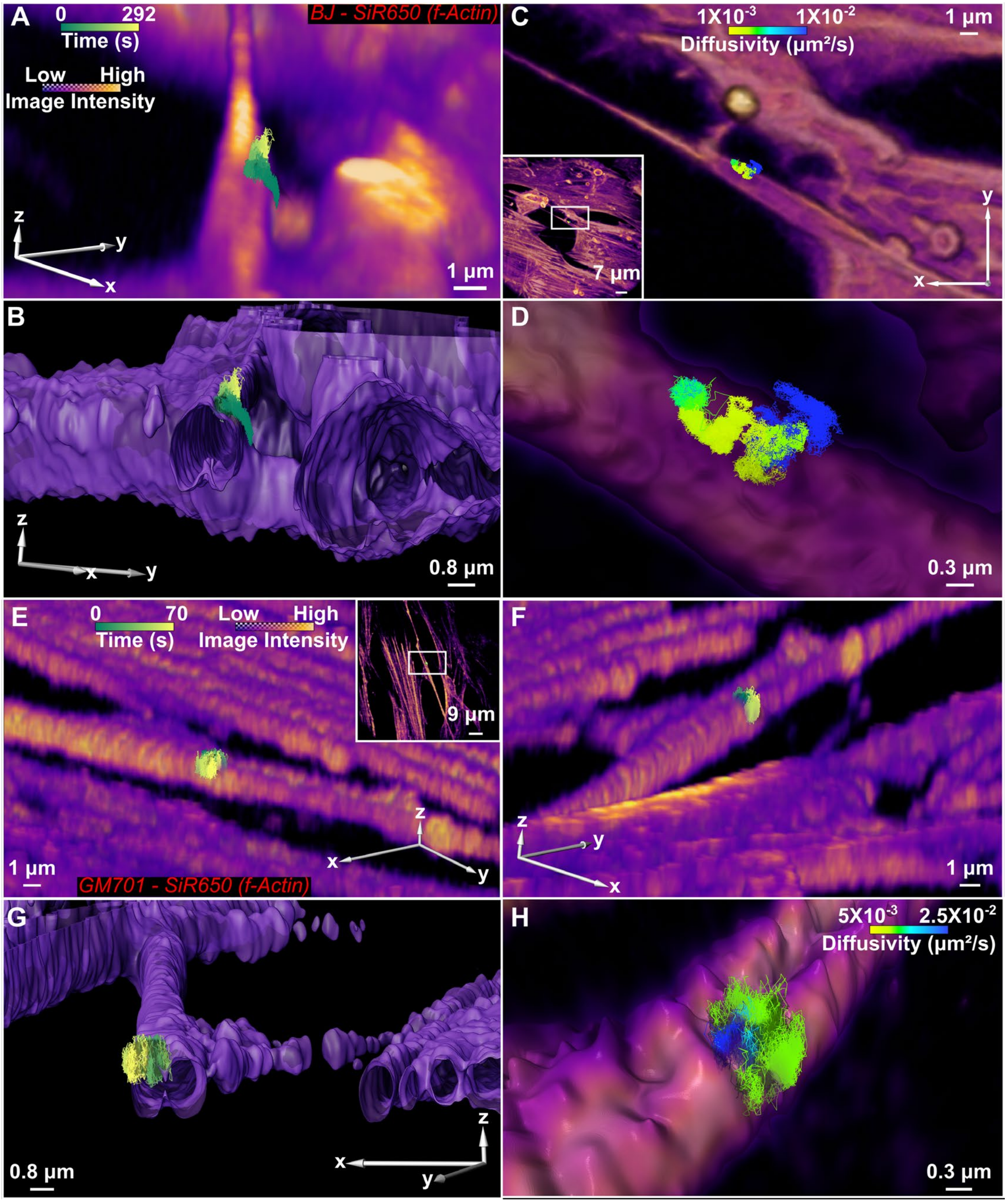

**Fig. S32. VSV-G VLP membrane diffusion.**

(**A**) VSV-G VLP captured diffusing on membrane of fibroblast protrusion. In contrast to other examples the diffusion is more random but does not circumnavigate the cylinder. (**B**) Same trajectory from a different angle with isosurface 3D reconstruction of the cell surface. (**C**) Top-down view. (**D**) Magnified top-down view. (**A** and **B**) Trajectory color-coded by time, while in (**C** and **D**) color-coded by diffusion coefficient. (**E**) A different trajectory on LDLR-deficient fibroblast protrusion. Again, linear motion is not observed, and neither is the circling behavior observed on micro-protrusions. (**F**, **G**, and **H**) Same trajectory from different angles. Trajectory color-coded by time in (**F** and **G**), while in (**H**) color-coded by diffusion coefficient. In (**G**) the render is sliced through to better show the contour of the trajectory.

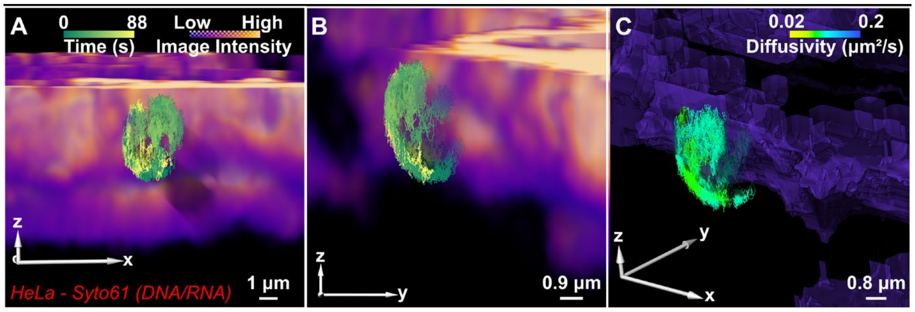

**Fig. S33. Long-term tracking allows the virus to trace out spherical features on the cell surface.**
(**A**) Axial view of VSV-G traveling diffusing on membrane bleb of SYTO61-stained HeLa cells, with an alternate orientation shown in (**B**) 3D volume rendering for data shown in (A), except trajectory colored diffusion coefficient. (**C**) 3D isosurface rendering for data shown in (A and B). In addition, trajectory color-coded by diffusion coefficient.

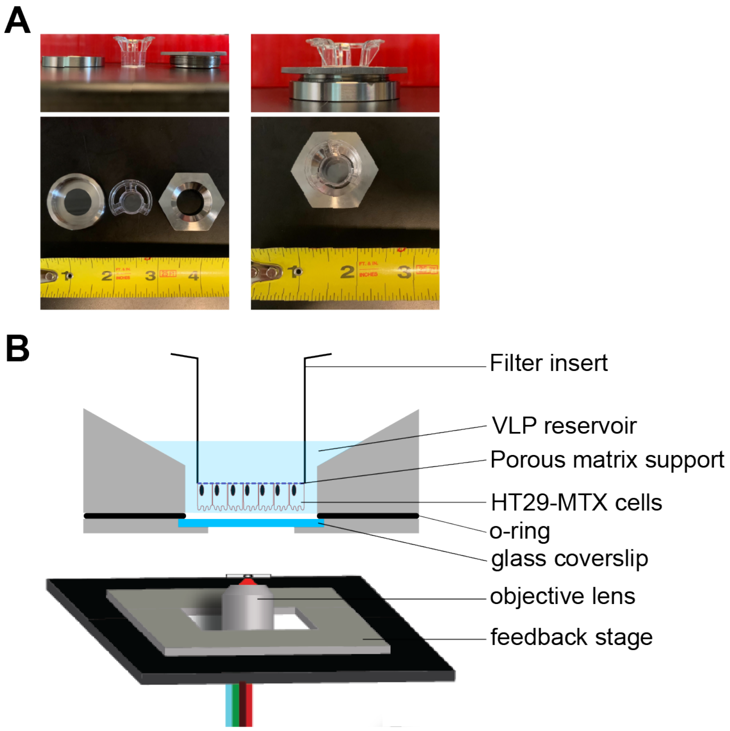

**Fig. S34. Sample holder used for filter-grown HT29-MTX cells, related to Fig. 5.**

(**A**) Sample-holder and filter-insert before (left), and after (right) assembly. (**B**) Schematic showing inverted cells inside chamber, this orientation promotes robust single-particle tracking.

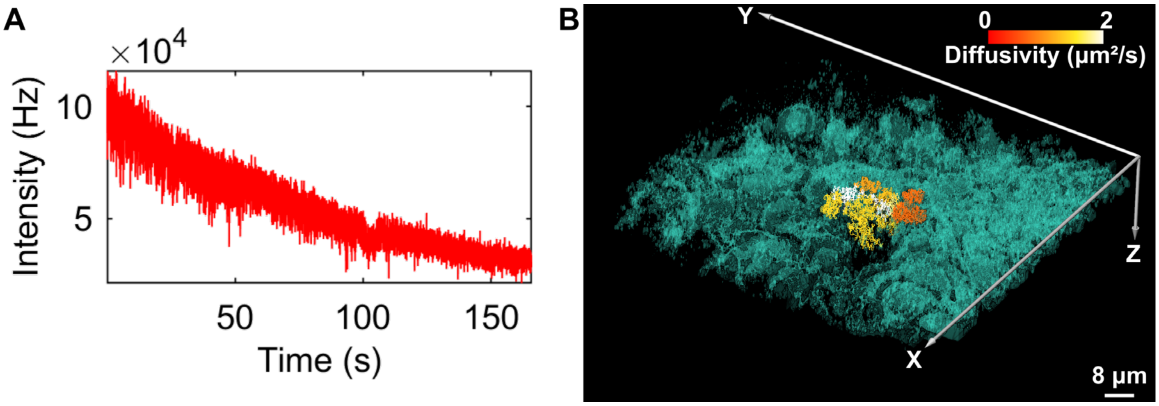

**Fig. S35. Accompanying data to support simultaneous tracking and imaging (TrIm) of VSV-G VLP burrowing through HT29-MTX cells, related to Fig. 5.**

(**A**) Tracking intensity trace. (**B**) 3D reconstruction from a 4D data set covering 10 local volumes, at 10 FPV (see movie S6) with trajectory split into unique segments of diffusion behavior via change-point analysis.

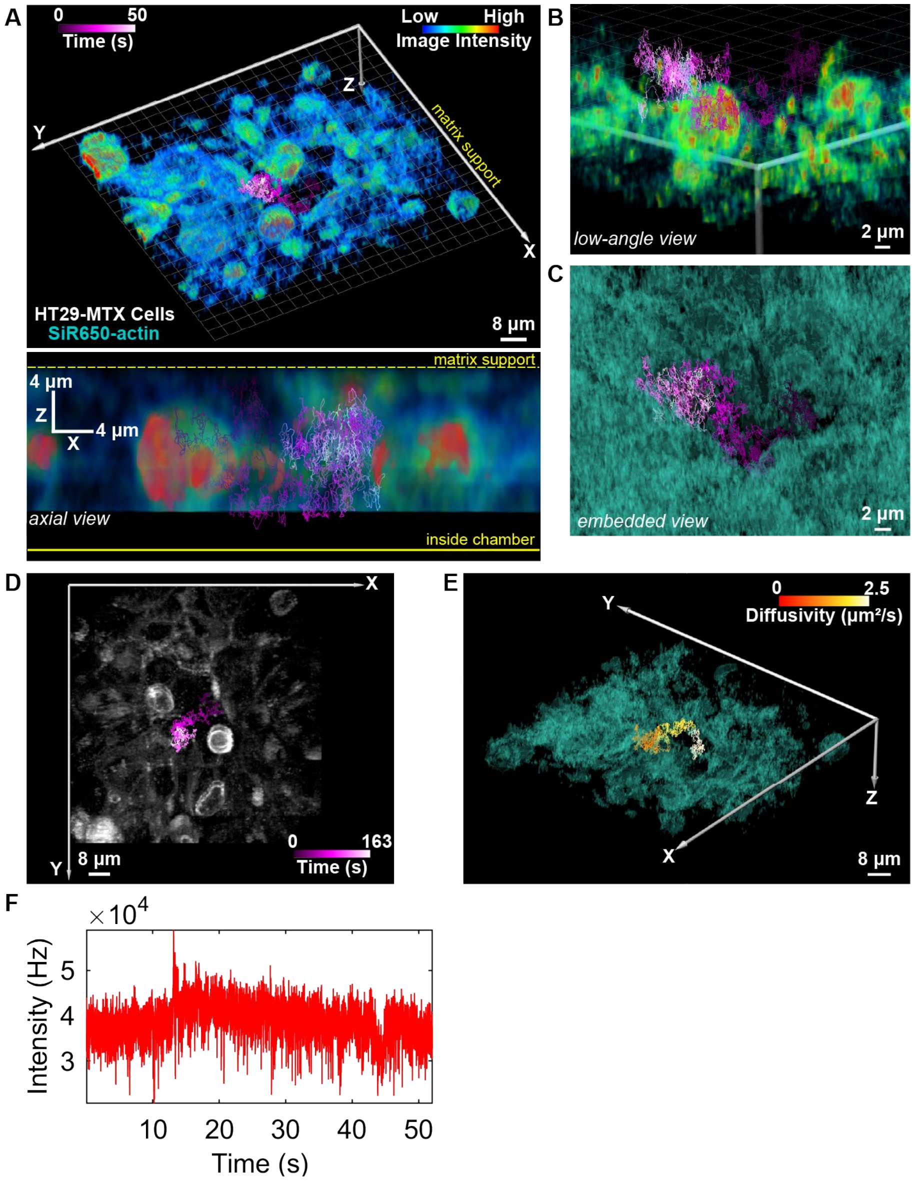

**Fig. S36. Additional example of VSV-G VLP circumnavigating HT29-MTX cells**

(**A**) 3D reconstruction from a 4D data set covering 3 local volumes, at 16 FPV of suspended HT29-MTX cells grown on inverted matrix stained with SiR650-actin with high resolution trajectory. Below, axial view. (**B**) Magnified view. (**C**) Cell boundary reconstruction highlighting sample density. (**D**) XY MIP. (**E**) 3D reconstruction of cellular imaging data in (A) with trajectory split into unique segments of diffusion behavior via change-point analysis. (**F**) Tracking intensity trace.

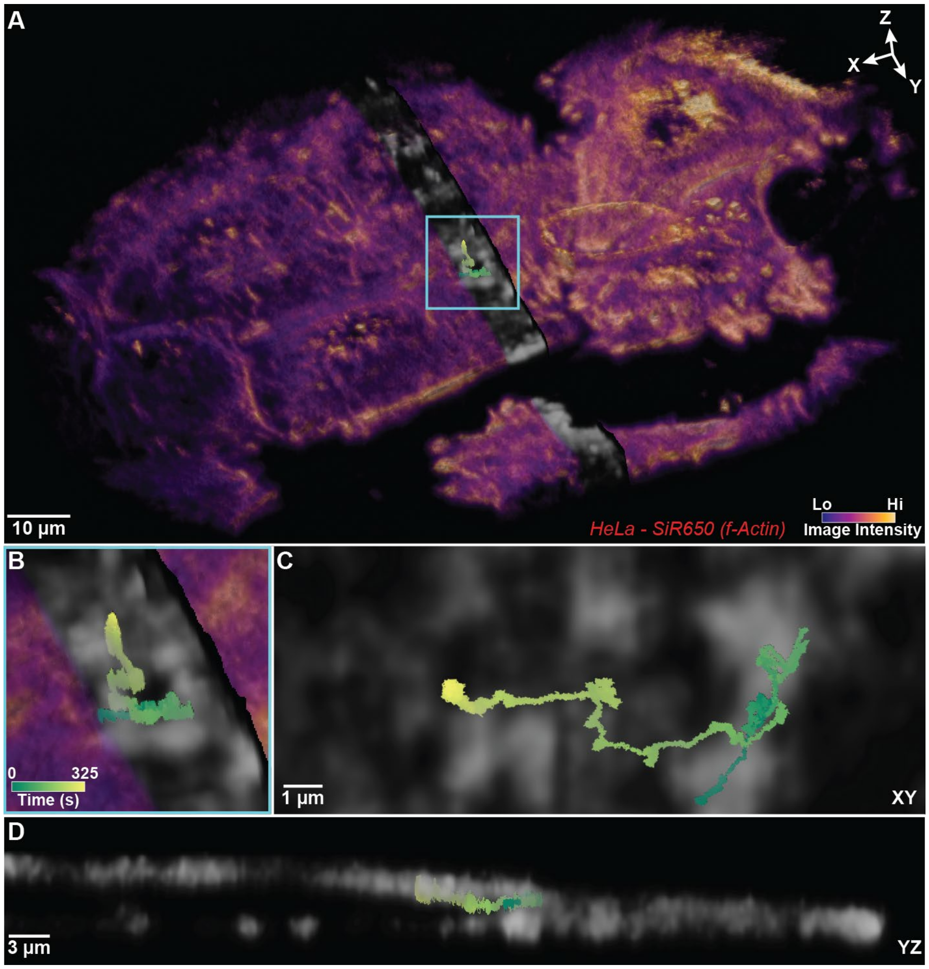

**Fig. S37. Internalized Viral Trafficking**

(**A**) Whole-volume visualization of HeLa cells stained with SiR-650 labeling *f*-actin. Two slice planes were used to section the YZ plane and create a channel which renders the interior volume section semi-transparent, revealing the trajectory embedded beneath the cell surface. (**B**) Close-in view of VLP trajectory. (**C**) Top-down XY view of internalized viral trajectory shows high-intensity, actin-rich regions that the trajectory moves around. (**D**) YZ slice view demonstrates that VLP is truly internalized inside the depths of the cell (see also: movie S7).

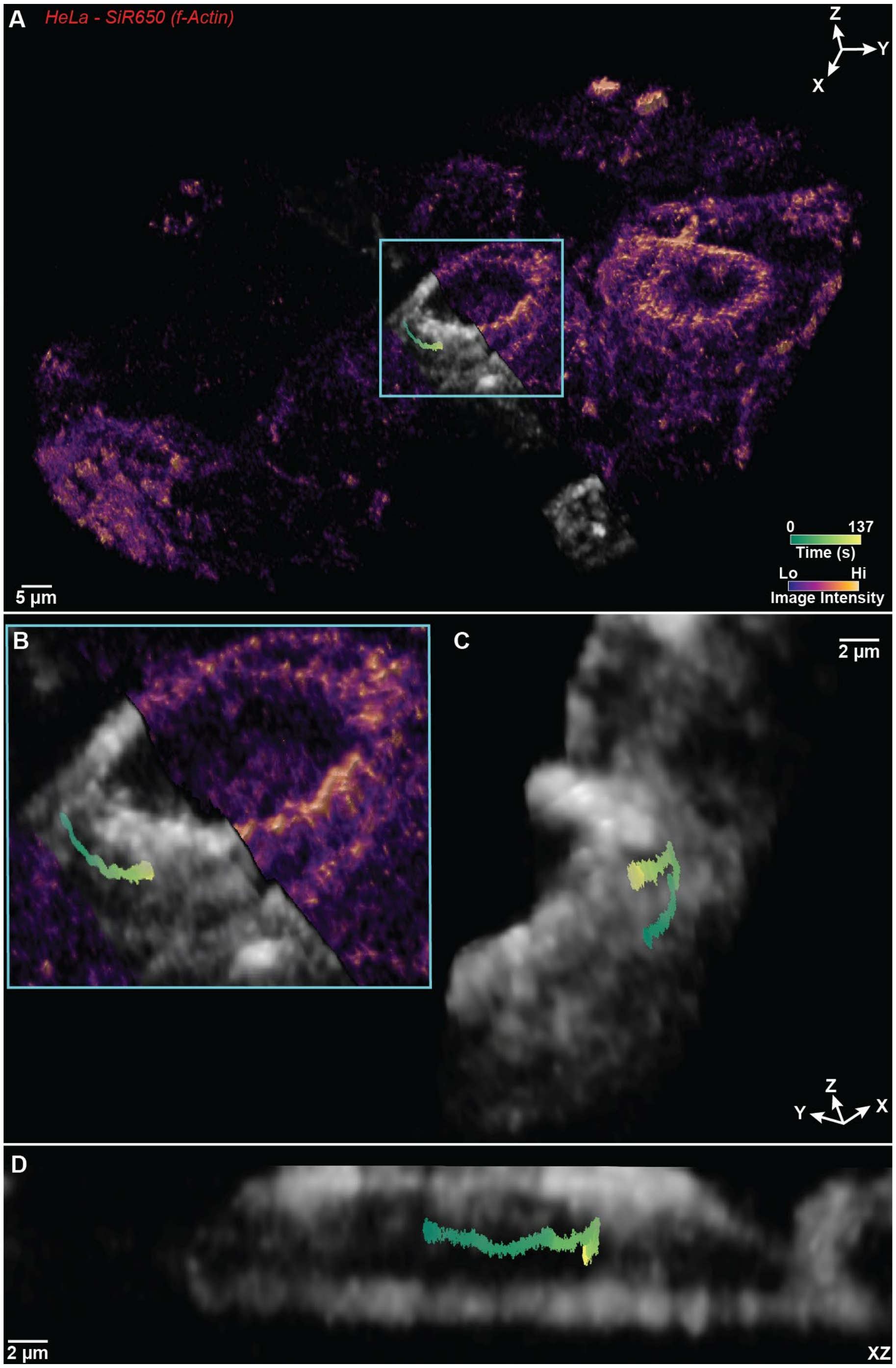

**Fig. S38. Linear Trafficking of Internalized VLPs.**

Wide view of HeLa cells stained with SiR-650-actin. A channel of lower opacity is used to reveal the embedded trajectory which appears to curve around the periphery of the nucleus. (**B**) Inset view of (**A**). (**C**) View of peri-nuclear region near trajectory shows sharp angle change with linear motion after. (**D**) XZ view shows viral trajectory embedded inside cell.

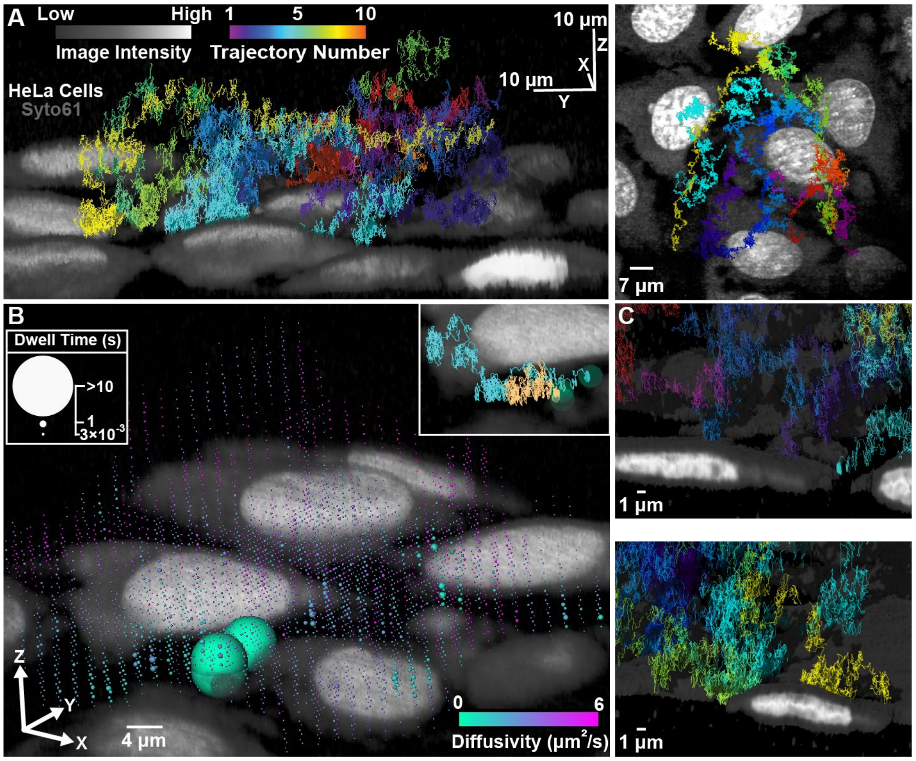

**Fig. S39. Multi-trajectory acquisition.**

(**A**) **Left:** 10 VSV-G VLP trajectories collected in the same area of live HeLa cells stained with SYTO61 (see also: fig. S38 and movie S7). **Right:** top-down (xy) MIP with multiple co-registered VLP trajectories. Colorbar represents trajectory number, with each trajectory uniquely colored. (**B**) The dwell time of 27 VSV-G VLP trajectories (starting from free diffusion) collected in the same area as (A) were binned into voxels and represented as spheres, color-mapped by diffusivity. This correlated analysis reveals commonly visited regions and highlights the slower diffusion experienced by the VLP population nearer the cell surface. **Inset:** two different VLPs land in close proximity on the same cell. (**C**) Exploded views showing multiple VLPs contact the same cell.

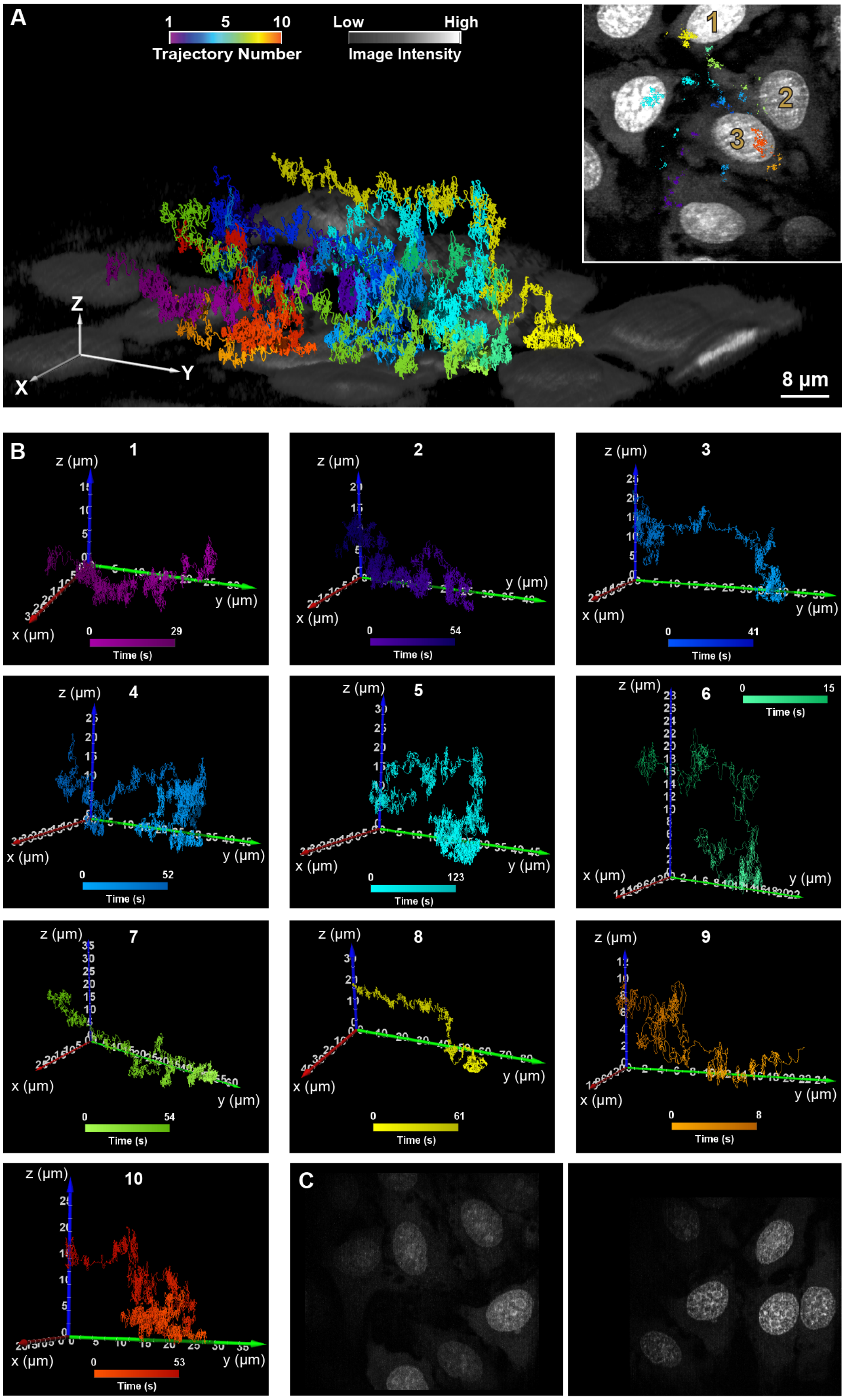

**Fig. S40. Multi-trajectory acquisition.**

(**A**) Complete field of view showing 3D reconstruction of live HeLa cells stained with SYTO61, co-registered with 10 VSV-G VLP trajectories. Insert, hot-spot analysis revealed cell 1 was visited trajectories 6,7, and 8 an accumulative total of 14.3 sec. Cell 2 visited by trajectories 4, 5, and 7 a cumulative total of 8.6 sec. Cell 3 contacted by trajectories 2, 3, 4, 5, 9, and 10 an accumulative total of 64.5 sec. Colorbar represents trajectory number, with each trajectory uniquely colored. In addition, the light to dark hue represents trajectory time. (**B**) Trajectories plotted individually, and color coded by time. (**C**) XY maximum intensity projections of local volumes from trajectories #4 and # 38 in a single area which represent the first and final trajectory where the virus remained in imaging proximity of the cells for at least one complete volume acquisition time. These MIPs demonstrate the ability for 3D-TrIm multi-trajectory visualization after 35.37 minutes of continuous tracking and imaging.

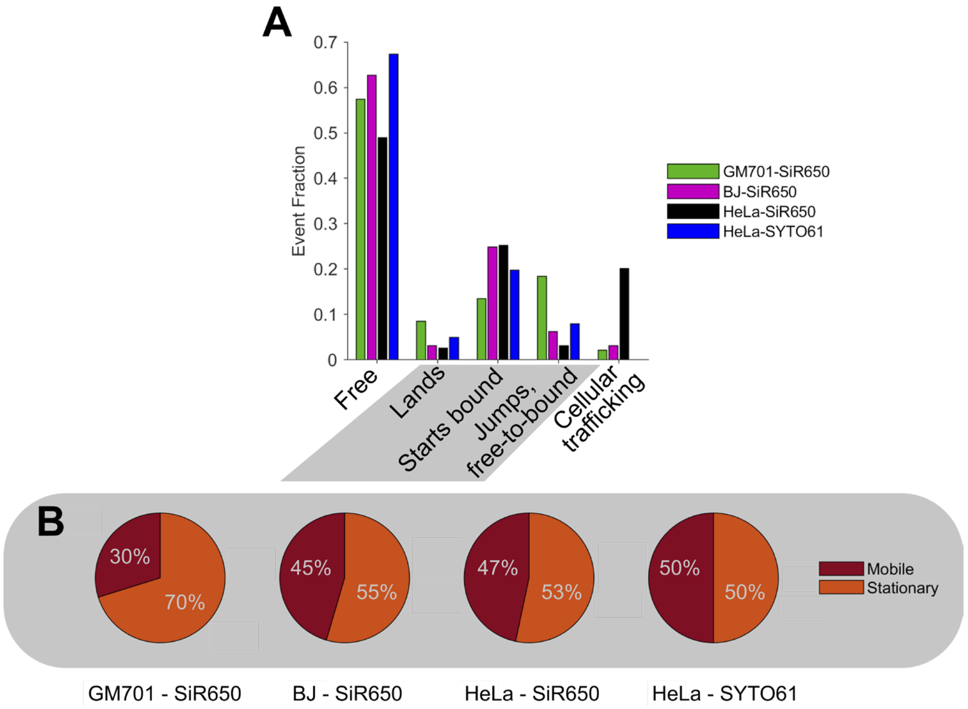

**Fig. S41. VSV-G VLP Fate Determination.**

(**A**) Trajectories classified into five distinct behaviors as described in section 1.8.5. VLPs either remain freely diffusing throughout trajectory duration (free), or land on a cell (lands), or begin on the cell surface (starts bound), or there is a jump between particles from those freely diffusing the those on the cell surface (jumps, free-to-bound), or finally the VLP trajectory begins inside the cell and is mobile (cellular trafficking). (n = 141, 225, 194, 101 trajectories for GM701-SiR650-actin, BJ-SiR650-actin, HeLa-SiR650-actin, HeLa-SYTO61, respectively). (**B**) Of those trajectories in (A) that were classified as having a bound segment (lands, starts bound, or jumps from free-to-bound) these were further classified as mobile or stationary by inspecting the diffusion coefficient as described in section 1.7.6.

**Movie S1. 3D-TrIm Method Animation Demo, related to Fig. 1.**

Demonstration of 3D-TrIm operating principle. Animation sequence begins with overview of experimental setup in which a heated sample containing virus-like particles (VLP) and live cells are mounted on a piezoelectric stage with an objective lens shared by both tracking and imaging microscope sources. This overview is followed by an animation of 3D-SMART real-time tracking, demonstrating how a pair of Electro-Optic Deflectors (EOD) create a lateral Knight's Tour grid pattern, followed by the use of a Tunable Acoustic Gradient (TAG lens) to scan a focal range above and below the center of the focal volume. A final animation demonstrates the principle of 3D-FASTR point-scan imaging.

**Movie S2. VSV-G Exploring the extracellular matrix, related to Fig. 2.**

(**A** and **B**) 3D reconstruction of real-time VSV-G VLP trajectory in extracellular matrix of live GM701 cells (stained with *f*-actin label SiR650-actin), from a 4D data set covering 10 local volumes, at 16 FPV. Trajectory (~ 162 sec) is segmented into 25 segments per second (25 frames per second when playback rate is 1×) and color mapped by time. The progress bar shows how the trajectory is further categorized: (1) Free diffusion period (playback rate: 2×): 0-14 sec, 18-38 sec, 44-62 sec, 70-108 sec. (2) Skimming period (playback rate: 1×): 14-18 sec, 38-44 sec, 62-70 sec, 108-122 sec. (3) Detachment (playback rate: 2×): 122-162 sec. Sphere represents the VLP position in the current frame (refreshing rate is consistent with the trajectory, i.e., 25 fps at 1× playback rate). Image volumes formed from maximum intensity projection over time from local volumes acquired over 16 frame-times. In (A) cells are color-coded by imaging intensity while in (B) cells are color-coded depending on distance of the virus from the cell surface. Panels (A and B) share the same trajectory color scale, camera angle and camera path, however, (A) is magnified compared to (B).

**Movie S3. VSV-G Searching Behavior, related to Fig. 3A-C.**

3D reconstruction of live HeLa cells (stained with nucleic acid label SYTO61) co-rendered with virus trajectory. Trajectory (~ 123 sec) is segmented into 25 segments per second (25 frames per second when playback rate is 1×) and color mapped by time. The progress bar shows how the trajectory is further categorized: (1) Free diffusion period (playback rate: 2×): 14-30 sec, 41-70 sec. (2) Skimming period (playback rate: 1×): 0-14 sec, 30-41 sec. (3) Immobilized period (playback rate: 4×): 70-123 sec. The end points of trajectory segments are labeled with spheres (refreshing rate is consistent with the trajectory, i.e., 25 fps at 1× playback rate), representing the position of the viral particle. The diffusional status of viral particle is further illustrated by the color change of the sphere (Gray: free motion, Cyan: skimming behavior, Purple: immobilization). Image volumes formed from maximum intensity projection over time from local volumes acquired over 16 frame-times. Cells are color-coded by image intensity.

**Movie S4. VSV-G Searching Behavior, related to Fig. 3E-G.**

3D reconstruction of live GM701 cells (stained with *f*-actin label SiR650-actin) co-rendered with virus trajectory. Trajectory (~ 54 sec) is segmented into 25 segments per second and color mapped by diffusivity. The playback rate is constant 1×. Image volumes formed from maximum intensity projection over time from local volumes acquired over 16 frame-times. Cells are color-coded by image intensity. We observed the VLP repeatedly landing and detaching.

**Movie S5. Virus Interaction with Protrusion, related to Fig. 4.**

3D reconstruction of live HeLa cells (stained with nucleic acid label SYTO61) co-rendered with virus trajectory. Trajectory (~ 133 sec) is segmented into 25 segments per second (25 frames per second when playback rate is 1×) and color mapped by time. Image volumes formed from maximum intensity projection over time from local volumes acquired over 16 frame-times. Cells are color-coded by image intensity. Shown in the progress bar, the trajectory is further categorized into two segments: (1) Free diffusion (playback rate: 1×): 0-16 sec. (2) Membrane bound diffusion on protrusion: 16-133 sec.

**Movie S6. VSV-G diffusing through multi-layered epithelial cells, related to Fig. 5.**

3D reconstruction of suspended HT29-MTX cells grown on inverted matrix support (stained with *f*-actin label SiR650-actin) co-rendered with virus trajectory. Trajectory (~ 166 sec) is segmented into 25 segments per second (25 frames per second) and color mapped by time. The playback rate is constant 4×. This movie shows how the global volume intensity is accumulated from local volumes acquired over 4 frame-times. Cells are color-coded by image intensity. Axes grid represents approximate position of matrix support on which the cells are suspended.

**Movie S7. Internalized Viral Trafficking, related to Fig. S37.**

3D reconstruction of live HeLa cells (stained with f-actin label SiR650) co-rendered with virus trajectory. Trajectory (~ 325 sec) color mapped by time. Global image volume formed from maximum intensity projection over time from local volumes acquired over 16 frame-times. Cells are color-coded by image intensity. Initially, the cells are opaque and the trajectory is not visible, a cut-through reveals the complete internalized trajectory. Viewing this embedded trajectory from above reveals high-intensity, actin-rich regions which the trajectory appears to navigate. Finally the camera is rotated to a lateral view to show the position of the trajectory within the 3D cell.

**Movie S8. Multi-Trajectory Overlay, related to Fig. S39.**

3D reconstruction of a single assembled area containing live HeLa cells (stained with nucleic acid label SYTO61) co-rendered with 10 virus trajectories. The total imaging data were acquired during the 10 trajectories and co-registered to form a single common area. Image volumes formed from maximum intensity projection over time from the assembled volumes acquired over 16 frame-times. Cells are color-coded by image intensity. 10 trajectories combined (~ 490 sec in total) are segmented into 25 segments per second (25 frames per second when playback rate is 1×). The trajectories are color mapped with respect to trajectory number (the color legend in lower left indicates the initial color of each trajectory). Each trajectory is further color mapped by time, as indicated in the progress bar. Trajectory #5 is related to movie S3.
